# Reliability of structural brain change in cognitively healthy adult samples

**DOI:** 10.1101/2024.06.03.592804

**Authors:** Didac Vidal-Piñeiro, Øystein Sørensen, Marie Strømstad, Inge K. Amlien, Micael Anderson, William F.C. Baaré, David Bartrés-Faz, Andreas M. Brandmaier, Anne Cecilie Bråthen, Pablo Garrido, Paolo Ghisletta, Håkon Grydeland, Richard N. Henson, Rogier A. Kievit, Max Kormacher, Simone Kühn, Ulman Lindenberger, Athanasia M. Mowinckel, Lars Nyberg, James M. Roe, Markus H. Sneve, Cristina Sole-Padulles, Leiv-Otto Watne, Kristine B. Walhovd, Anders M. Fjell

## Abstract

In neuroimaging research, tracking individuals over time is key to understanding the interplay between brain changes and genetic, environmental, or cognitive factors across the lifespan. Yet, the extent to which we can estimate the individual trajectories of brain change over time with precision remains uncertain. In this study, we estimated the reliability of structural brain change in cognitively healthy adults from multiple samples and assessed the influence of follow-up time and number of observations. Estimates of cross-sectional measurement error and brain change variance were obtained using the longitudinal FreeSurfer processing stream. Our findings showed, on average, modest longitudinal reliability with two years of follow-up. Increasing the follow-up time was associated with a substantial increase in longitudinal reliability while the impact of increasing the number of observations was comparatively minor. On average, 2-year follow-up studies require ≈2.7 and ≈4.0 times more individuals than designs with follow-ups of 4 and 6 years to achieve comparable statistical power. Subcortical volume exhibited higher longitudinal reliability compared to cortical area, thickness, and volume. The reliability estimates were comparable to those estimated from empirical data. The reliability estimates were affected by both the cohort’s age where younger adults had lower reliability of change, and the preprocessing pipeline where the FreeSurfer’s longitudinal stream was notably superior than the cross-sectional. Suboptimal reliability inflated sample size requirements and compromised the ability to distinguish individual trajectories of brain aging. This study underscores the importance of long-term follow-ups and the need to consider reliability in longitudinal neuroimaging research.

## 1. Introduction

Reliability and validity are fundamental to scientific progress. Reliability refers to the consistency of repeated measurements, while validity refers to the extent to which a measure captures what it intends to capture (Lavrakas, 2008). Reliability places an upper limit on validity (Spearman, 1904) and has severe implications for interpretation and statistical power in individual differences research (Parsons et al., 2019; Zuo et al., 2019). In humans, structural magnetic resonance imaging (MRI) features are key to understanding the aging brain and how individuals differ. Cross-sectional estimates are likely to be invalid measures for capturing interindividual differences in brain *aging* (i.e., brain change), as they largely reflect lifelong differences between individuals (Raz and Lindenberger, 2011; Vidal-Pineiro et al., 2021). As a result, there is an increasing availability of longitudinal cohorts. Yet, estimating differences in intraindividual change (longitudinal) is often less reliable than those made on level (e.g. cross-sectional) as variance in change tends to be considerably smaller (Hertzog et al., 2008). This sets limits to the validity of individual differences of brain change estimates. Here, we attempt to estimate the reliability of longitudinal brain change for structural MRI brain features, and the follow-up time and number of observations required to achieve different levels of reliability in cognitively healthy adults.

Measurement reliability is classically defined as the portion of variance attributed to true scores (i.e., between-subjects) relative to the total variance (between and within-subject variance) (Allen and Yen, 2001). Reliability is indirectly related to statistical power in experimental designs (e.g., control vs. treatment) where the interest typically is in precision and thus in reducing both between and within-subject variance (Hedge et al., 2018; Zimmerman and Zumbo, 2015). Yet, this index is key in individual differences research (Brandmaier et al., 2018b; Hedge et al., 2018; Zimmerman and Zumbo, 2015), and consequently in any attempt to understand how interindividual variations in brain change are related to genetic, cognitive, or environmental factors. Using brain change estimates with sub-optimal reliability may have severe consequences for the interpretation, comparability, and reproducibility of results (Parsons et al., 2019). Low reliability reduces statistical power and increases uncertainty in the parameter estimates, leading to false negative results and attenuated estimations of the effects, but also potentially producing false positives and artificially inflated effects when combined with other sources of bias (Button et al., 2013; Loken and Gelman, 2017; Spearman, 1904). Low reliability hampers the validity of the results and lowers the reproducibility across studies and as such, in aging neuroimaging, limited longitudinal reliability is considered an important factor for the lack of converging evidence (Oschwald et al., 2019).

In neuroimaging research, reliability is often assessed as test-retest reliability via repeated scans acquired within the same session, after repositioning, or after some time (Brandmaier et al., 2018b; Hedges et al., 2022; Madan and Kensinger, 2017; Parsons et al., 2024). Core measures of brain structure such as thickness, area, and volume have almost invariably shown high test-retest reliability, e.g., intra-class correlation coefficients (ICC) often > .8, across different scanners, sequences, processing pipelines, and populations (Hedges et al., 2022; Iscan et al., 2015; Liem et al., 2015; Madan and Kensinger, 2017; Sederevičius et al., 2021) (c.f. Parsons et al., 2024). Mimicking this approach is more challenging for longitudinal reliability as it requires two assessments at each time point (c.f. Takao et al., 2022, 2021). Alternatively, the reliability of brain change can be analytically derived by estimating the *true* and error variance of the brain *slopes* as in the growth rate reliability (GRR) index (Rast and Hofer, 2014; Willett, 1989) or in its generalization to latent variable models, i.e., effective curve reliability (ECR) (Brandmaier et al., 2018a). *True variance* is defined by slope variance, that is, the degree to which the individuals vary in their slopes while error variance of the slopes is dependent on measurement error variance, the duration of the study, the number of observations, and the spacing of these observations (Brandmaier et al., 2018a; Hertzog et al., 2008; Rast and Hofer, 2014; Willett, 1989). Here, we follow a similar approach to estimate the reliability of longitudinal brain change.

Increasingly prevalent data-sharing practices mean that the majority of neuroimaging research is authored by researchers who lack the ability to influence data collection plans (Milham et al., 2018). This lack of control is particularly important as practical constraints on collecting longitudinal neuroimaging data, such as economic costs, resources, time, retention, and selective attrition issues, often favor a particular kind of neuroimaging design in terms of number of observations, interval, and sample size. That is, most research in the aging neuroimaging field is performed with (relatively) open data with a limited follow-up time and number of observations. As such, we directed our attention to the reliability *a posteriori*, i.e., in already collected data that is widely used in *secondary* analysis in the neuroimaging field. See elsewhere for *a priori* assessments of longitudinal reliability (Brandmaier et al., 2018a, 2015; Hertzog et al., 2008; Rast and Hofer, 2014; von Oertzen, 2010), where reliability can be optimized before data acquisition by modifying spacing, follow-up time, and number of observations. Here, we estimated the reliability of longitudinal brain change for regional and global cortical thickness, area, and volume features, as well as for subcortical structures, using data from multiple cohorts and the longitudinal stream of FreeSurfer (Reuter et al., 2012a). We explored the impact of total follow-up time and number of observations on longitudinal reliability (see **Figure 1** for schematic representation) as well as the consequences on required sample sizes and the ability to accurately define and distinguish individual trajectories of aging. Finally, reliability is, ultimately a property of the measurement, influenced by the measure, but also partially dependent on the sample (e.g. Appelbaum et al., 2018; Parsons et al., 2019), and in neuroimaging, also of the processing stream. Hence, we illustrated the dependence of the reliability estimates on the population of interest, the processing pipeline, and explored to what extent the results are replicated across different cohorts. We offer a **supporting app** (https://vidalpineiro.shinyapps.io/longrho_shinyapp/) to aid researchers in estimating the reliability of longitudinal change.

**Figure 1.**
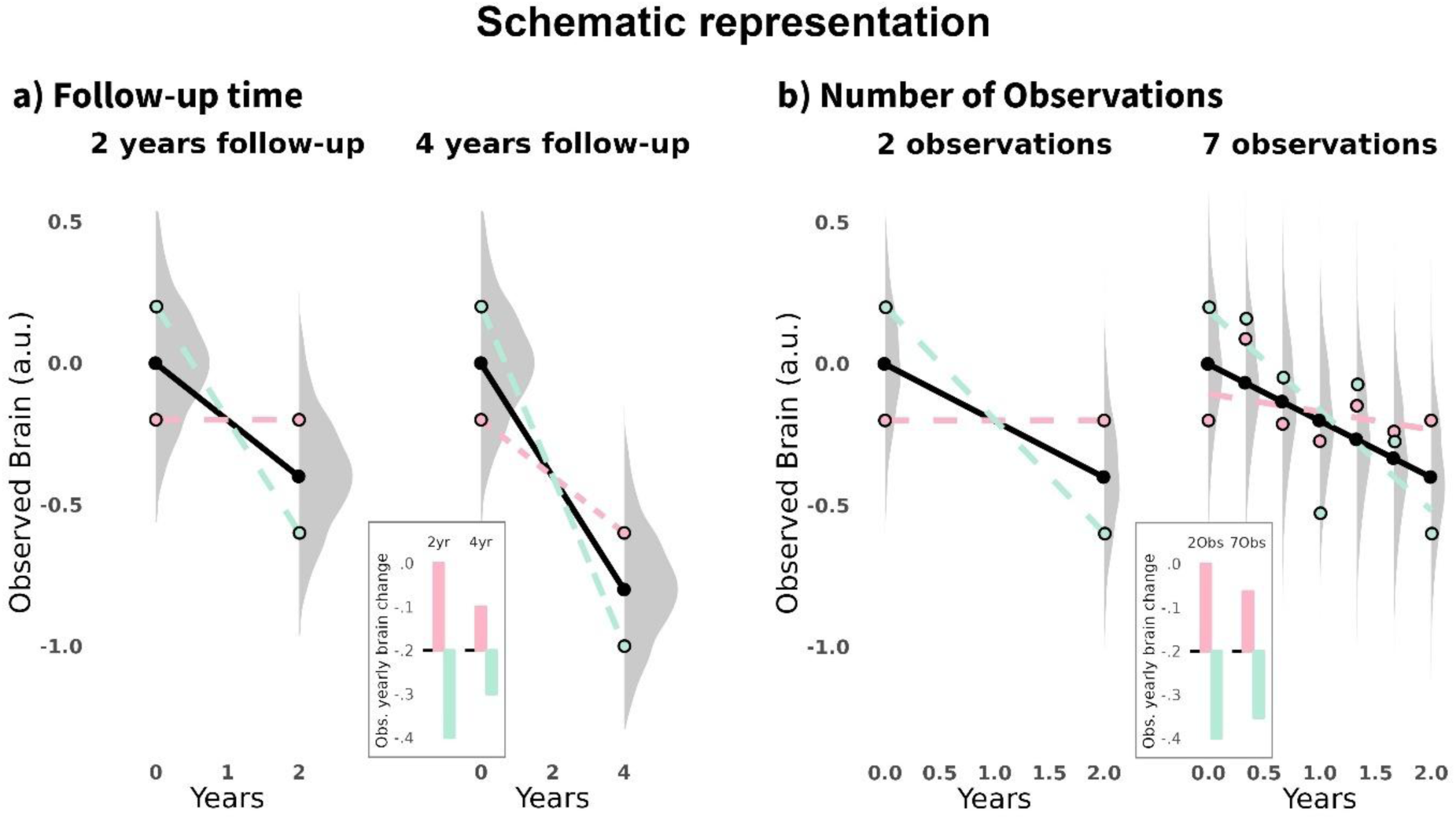
Schematic representation of time and follow-up effects on reliability. Hypothetical scenario illustrating a participant scanned twice at each time point, represented by the green and red lines. Measurement error causes deviations in the estimated slopes from the true trajectory (black line), which represents actual change over time. In the main plots, points represent observed cross-sectional measurements, lines estimated longitudinal (linear) trajectories, and density plots represent the distribution of possible values for a given cross-sectional observation. The boxes show the observed yearly brain change. a) Effects of follow-up time: Extending the follow-up time from 2 years to 4 years reduces the impact of cross-sectional measurement error on yearly change estimates. b) Effects of increasing the number of observations which leads to reductions of measurement error on yearly change estimates.

## 2. Methods

### 2.1. The growth rate reliability index

See **table 1** for a summary of key concepts, definitions, and measurements. To simplify estimations of longitudinal reliability, we made some initial assumptions. First, let us assume individuals have their specific, independent responses from each other. The repeated measures for each individual (i) measured at a given set of occasions (j) (j_1_,…, j_n_), can be expressed as follows (eq. 1) and change can be expressed in terms of linear trends and captured by slope estimates. We will, for simplicity, and in line with most available longitudinal neuroimaging data, assume equispaced measurements and the same number of measurements across individuals. We assume that the slopes of change (*β*_2*i*_) are linear, normally distributed in the population with mean δ and variance 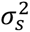 and similarly for (crosssectional) measurement errors (*ε*_*i*,*j*_) with a mean equal to 0 and variance σ_ε_^2^. Although not of direct interest in what follows, the intercepts (*β*_1*i*_) are also assumed to be normally distributed with a nonzero mean and variance.

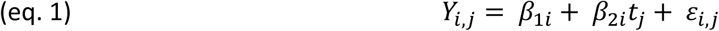

**Table 1.**
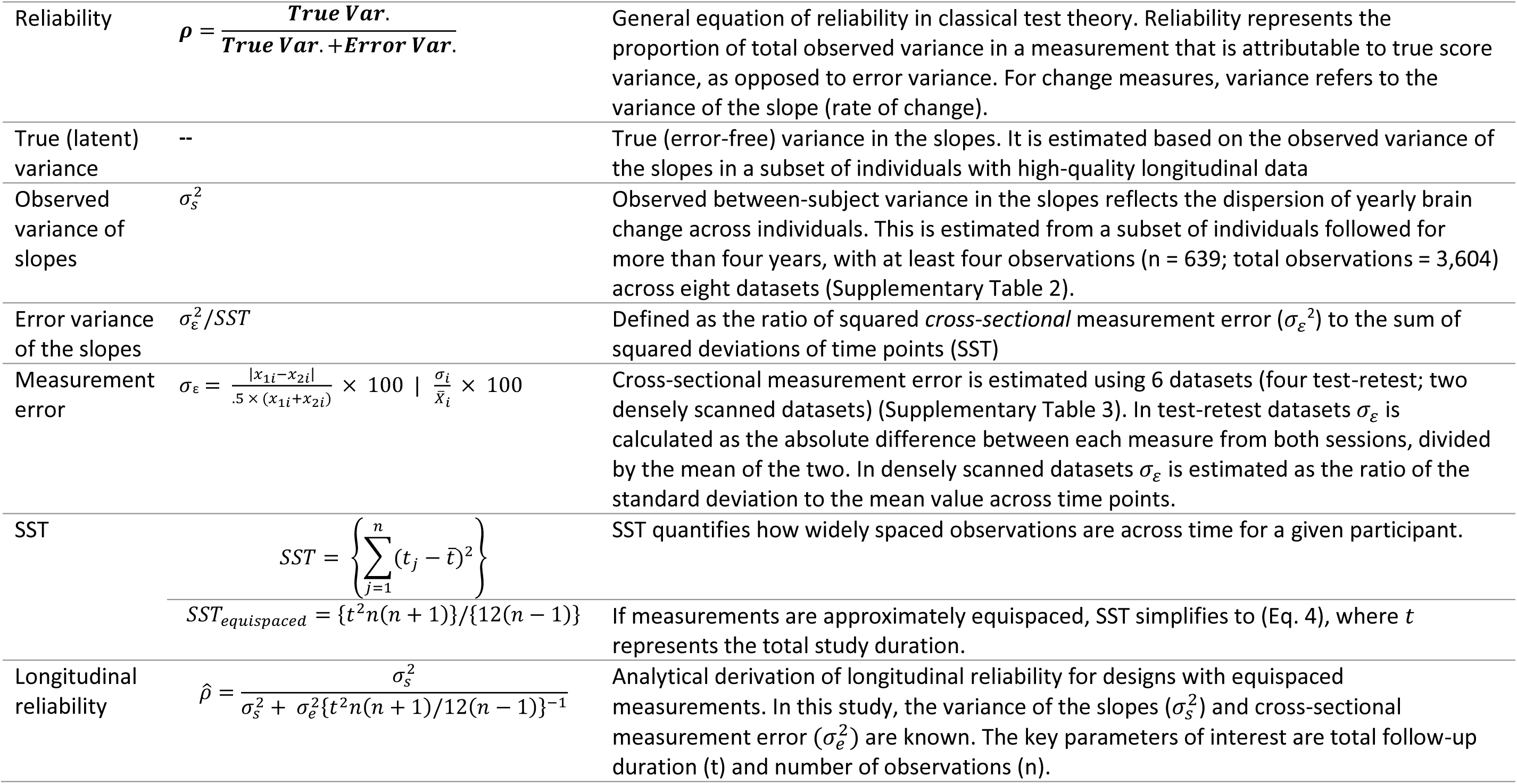
Summary Panel. Key study measures, definitions, and measurements. Var. = Variance.

Also, as defined by classical test theory, the reliability coefficient (*ρ*) is defined as the ratio of the true score variance by total variance. Total variance is defined as the sum of the true score and the error variance (eq. 2).

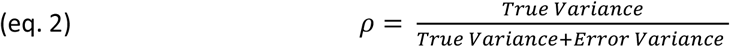

In a longitudinal analysis, and assuming linear changes, *true* variance can, generally, be estimated using linear regression separately on individuals with three or more measurements which allows separation from measurement error (c.f. Brandmaier et al., 2024). We used the variance of slopes 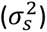 as the estimate of *true (latent)* variance. Slope variance refers to the *observed* variability in individual brain change slopes, while true variance refers to the latent, unobserved variability, free from measurement-related distortions. Note that 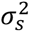, is subject to two opposing influences. On one hand, it is overestimated due to error propagation, while on the other, it is influenced by study design and sample characteristics including drop-out, motivation, mortality, etc., which set constraints to the sample’s variance. That is, attrition bias leads to underestimation of the variance of slopes. We assumed that 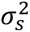 from data with long follow-ups and a high number of observations provides a close approximation of the *true* variance of linear change as the impact of error propagation is minimized. Our simulations showed that the overestimation of the variance of the slopes is approximately 10% (**Supplementary Information** and **Supplementary Figure 1)**. The extent to which the variance of the slopes is underestimated due to attrition bias is more challenging to estimate. See *Discussion* for an in-depth discussion.

The error variance of the slope, which quantifies the uncertainty associated with a given slope estimate, is defined by the ratio of the squared *cross-sectional* measurement error (*σ*_*ε*_) to the sum of squared deviations of time points (SST) (Willett, 1989). In this case, SST captures how widely spaced the observations are for a given participant across time (eq. 3). The SST summation simplifies to (eq.4) if the measurements are approximately equally spaced (*τ* denoting the total duration of the study) (Fitzmaurice et al., 2012) (von Oertzen, 2010).

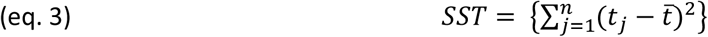

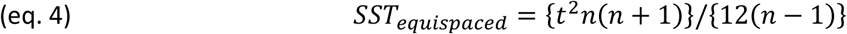

Hence, reliability for longitudinal brain change 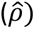 can be computed as follows (eq. 5) and represents a simplification of both the Growth Rate Reliability (GRR) index (Willett, 1989) and the Effective Curve Reliability (ECR) (Brandmaier et al., 2018a), which quantify the ability to distinguish differences in slope parameters (Rast and Hofer, 2014). In addition to the total follow-up time (*t*) and number of observations (n) - the parameters of interest in this study - we need information on the variance of slopes 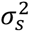 and (cross-sectional) measurement error *σ*_ε_, estimated, for example, from test-retest data. Note from eq. 5 that higher reliability across features is determined by a greater ratio of variability of the slopes to measurement error (**Supplementary Figure 2)**.

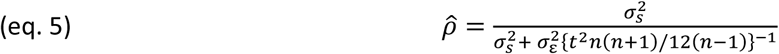

### 2.2 Parameter selection

Two different multi-cohort datasets were used to obtain the parameters of slope variance 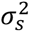 and measurement error (*σ*_*ε*_). The studies were approved by the relevant ethical committees and conducted in accordance with the Declaration of Helsinki. In both cases, data consisted of structural T1-weighted (T1w) scans that were collected using 1.5, 3, and 4 T scanners. T1w scans were preprocessed with the longitudinal FreeSurfer v.7.1 stream (Reuter et al., 2012a) (Dale et al., 1999; Fischl et al., 1999). Cortical thickness, area, and volume data (modalities) were summarized based on the Desikan atlas (Desikan et al., 2006) (|N| = 34 regions of interest [ROIs] per hemisphere) while left and right Lateral Ventricle, Thalamus, Caudate, Putamen, Pallidum, Hippocampus, and Amygdala volumes were extracted based on the *aseg* atlas. The combination of modality (e.g., thickness) and region (e.g., entorhinal cortex) are henceforth referred to as *features*. See **Supplementary Methods** for more information and **Supplementary Table 6** for MRI acquisition parameters.

#### 2.2.1 Slope variance

To obtain 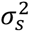, we used a multicohort longitudinal dataset (n = 11 datasets, n = 3611 unique individuals, n = 10964 observations) consisting of cognitively healthy adult participants. The datasets include the LCBC (Walhovd et al., 2016), Umeå (Nyberg et al., 2010), and UB (Rajaram et al., 2017; Vidal-Piñeiro et al., 2014) datasets (from the Lifebrain Consortium) (Walhovd et al., 2018), COGNORM (Idland et al., 2020), the Alzheimer’s Disease Neuroimaging Initiative (ADNI) database (https://adni.loni.usc.edu) (Mueller et al., 2005), The Australian Imaging, Biomarker & Lifestyle (AIBL) Study of Ageing (Ellis et al., 2009), Harvard Aging Brain Study (HABS) (Dagley et al., 2017), UKB (https://www.ukbiobank.ac.uk/) (Miller et al., 2016), PREVENT-AD (Breitner et al., 2016; Tremblay-Mercier et al., 2021), OASIS3 (LaMontagne et al., 2019), and Wayne (Daugherty and Raz, 2016; Raz et al., 2012) datasets. Observations concurrent with cognitive impairment and Alzheimer’s dementia were excluded. 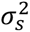 was estimated using a subset of these individuals followed > 4 years and with 4 or more observations (n = 639, observations = 3604) from 8 of these datasets. See **Supplementary Figures 3, 4,** and **Supplementary Tables 1, 2** for the sample’s descriptive statistics and visualization. See **Supplementary Table 5** for data availability. See **Supplementary Information** for a detailed description of the datasets, sample description, and image preprocessing. Values of each neuroimaging feature were fitted using generalized additive mixed models (*gamm4 R-package*) (Wood, 2017) that included age as a smooth term, sex as a covariate, and random intercepts for cohort, site (scanner), and participant. This step removes non-linear age trends at the sample level and harmonizes data across datasets and scanners. For each individual and feature, we estimated the rate of brain change by regressing the *gamm* residuals on follow-up time. Slopes of change were converted to percentage change scores based on the individuals’ mean values. This step was performed so features from different modalities are directly comparable despite possibly minor differences in reliability compared to *raw* scores. Next, extreme outliers, defined by values > 5 mean absolute deviation (MAD) around the fitted mean (multiplied by a MAD to SD scaling factor), were discarded. We used the (squared) standard deviation of the slopes as the measure of interest.

#### 2.2.2. Cross-sectional measurement errorc

Measurement error σ_ε_ was estimated as the average error across six different test-retest cohorts consisting of cognitively healthy adult participants, namely, the S2C (Walhovd et al., 2024), the preventAD (Orban et al., 2015), OASIS (Marcus et al., 2007), and GSP (Holmes et al., 2015) reliability subsets, and the HNU1 (Chen et al., 2015) and Maclaren (Maclaren et al., 2014) test-retest datasets (n = 341, observations = 1036). Three of the datasets partially overlapped with datasets used for estimating slope variance. See **Supplementary Table 4** for the sample’s descriptive statistics and visualization. See **Supplementary Table 5** for data availability. See **Supplementary Information** for a detailed description of the datasets, sample description, and image preprocessing. Briefly, the datasets consisted either of test-retest designs, performed on different days (≤ 3 months) (|N| = 4) (mean interscan interval = 77.2, 20.1, 111.4, and 82.8 days per dataset, respectively) or of cohorts of densely scanned participants over short periods (|N| = 2) (mean time between first and last observations = 33.1 and 31.0 days per dataset, respectively). Extreme outliers (>5 MAD around the mean to the between-subject average for test-retest and to the within-subject average for densely scanned designs) were discarded. Measurement error (*σ*_*ε*_) was estimated for a given subject (i) as the absolute difference between each measure estimated from both sessions divided by the mean of the two for test-retest designs (eq. 6), while for densely scanned designs, we computed the coefficient of variation (eq. 7), where *σ*_*i*_ denotes standard deviation and *X̄*_*i*_ the mean value across timepoints. The mean across subjects was estimated in each cohort while the mean across cohorts was the parameter of interest.

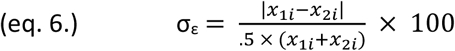

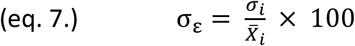

#### 2.2.3 Follow-up time and number of observations

We explored follow-up durations (t) between 2 and 12 years (sampled every two years) and 3, 5, 7, and 9 observations (n). Most of the existing longitudinal MRI data falls below the upper follow-up and number of observations limits. We have included estimates of longitudinal reliability for 3 or more observations to enable comparisons with the empirical estimations which require 3 or more observations. A somewhat arbitrary lower limit of 2 years was set, as shorter follow-up times are rarely used for studying individual differences in *brain aging*. Rather, the available datasets are often part of experimental designs. In any case, reliability estimates outside the reported bounds can be explored in the **supporting app**.

### 2.3 Higher level analysis

All the analyses were carried out in the R environment (R Core Team, 2023). Values in parenthesis represent standard deviations (SD), unless otherwise stated. We chose (left) hippocampus volume, and entorhinal thickness to illustrate the different results in specific features. Both measures are widely used in the context of cognitive neuroscience of aging, especially in relation with episodic memory function. Hippocampus is amongst the features with higher longitudinal reliability, while entorhinal thickness ranks poorly. Visualizations were made with the *ggplot2* (Wickham, 2016) and the *ggseg* (Mowinckel and Vidal-Piñeiro, 2020) R-packages.

#### 2.3.1 Effects of follow-up time, modality, and number of observations

A three-way ANOVA was carried out with all estimates of longitudinal reliability with modality, total follow-up time, and number of observations as predictors.

#### 2.3.2 Agreement across datasets for parameters and reliability estimates

Next, we explored whether measurement error (σ_ε_) and slope dispersion parameters (σ_s_) were comparable across datasets using single (ICC(2,1)) and mean reliability (ICC(2,k)) (McGraw and Wong, 1996; Shrout and Fleiss, 1979). The first index provides information on the reliability when using a single source for extracting parameters while the latter provides the reliability of the mean measurement. Similarly, we explored the reliability of the longitudinal reliability estimates using independent pairs of error and slope dispersion parameters. For overall consistency, we used k = 6 for ICC for mean reliability, as this represents all possible combinations of independent pairs. We used a random subset (n = 500) of possible combinations. We assessed global and local consistency, that is, reliability for the entire model and the model when constrained to a given number of observations and follow-up time. These indices assess the generalizability and representativeness of the parameters used, and the reported reliability estimates across different datasets.

#### 2.3.3 Consequences of longitudinal reliability (I): Sample Size Estimates

We estimated the required sample size given the longitudinal reliability predicted as a function of feature, number of observations, and follow-up time. We assumed a Pearson’s correlation with different *true* effects, 80% power, and perfect reliability for the other variable, and derived the attenuated correlation based on estimated reliabilities (Spearman, 1904). See the **supporting app** for sample size estimations of two-sample t-tests, and three-level ANOVAs following the formulas described elsewhere (Kanyongo et al., 2007; Zuo et al., 2019).

#### 2.3.4 Consequences of longitudinal reliability (II): Misclassification of individual trajectories

Next, we illustrated the degree to which we correctly identify individuals with differing aging trajectories as a function of feature, number of observations, and follow-up time. Hence, we defined three *hypothetical* individuals: a *normal ager*, a *maintainer*, and a *decliner* which decline 0, +1, and −1 standard deviations faster than the population average. Mean annual change – estimated as the yearly brain change between 60 and 80 using GAMM derivatives (*gratia r-package)* (Simpson, 2024) - and its variability across individuals was available from the dataset used for estimating slope variance (section 2.2.1). For each *hypothetical* subject, feature, number of observations, and follow-up time, we computed the probability density functions of the possible *observed* slopes using the parameters described above. We then assessed 1) the amount of overlap between these distributions using the Bhattacharyya coefficient (eq. 8) where *p(x)* and *q(x)* are the probability density functions of the observed slopes for, e.g., the maintainer and the normal ager and 2) estimated the probability of incorrectly identifying (ordering) these *hypothetical* individuals. Note that the probability of incorrectly identifying these individuals at random is 50%.

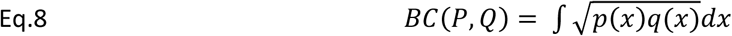

#### 2.3.5 Consequences of longitudinal reliability (III): Group membership based on trajectories

We used eq. 1 to estimate individual trajectories of brain change. In addition to measurement error and slope dispersion measures, we also used mean change (decline) and mean values. Using the *gamm* models described above (see *Parameter selection* section), we took the average values at age = 70 as the group mean, and the mean average derivatives between 60 and 80 as yearly change (*gratia r-package)* (Simpson, 2024). We simulated samples of 1,000 individuals for each cell (number of time points × number of observations). Based on the simulated samples, we estimated: a) the proportion of participants with no observed (measured) decline (observed brain maintainers); the proportion of those who show b) true (*latent)* decline (true brain maintainers), and c) true above-average decline (true brain decliners).

#### 2.3.6 Determinants of longitudinal reliability (I): Sample characteristics

Reliability is a property of the measurement and, thus, partially sample dependent. Hence, we reestimated the reliability of longitudinal brain change using slope variance parameters extracted from young and old adult subsamples (cut-off at 60 years, N = 70 and 569, respectively). A more refined approach to estimating age-dependent variability in brain change involves models such as generalized additive models for location, scale, and shape (GAMLSS). However, accurately capturing age-dependent dispersion with these models requires a significantly larger sample size than what was available in the present study.

#### 2.3.7 Determinants of longitudinal reliability (II): Preprocessing stream

In neuroimaging, reliability is not only a property of the measurement but also dependent on the preprocessing stream. To illustrate this, we re-estimated the reliability of longitudinal brain change using the cross-sectional FreeSurfer processing stream. The longitudinal stream is generally recommended for longitudinal analyses; however, it is computationally more demanding and can be susceptible to biases when observations are acquired at uneven time intervals or when major brain events occur. As a result, the cross-sectional stream continues to be widely used in longitudinal designs. The differential reliability between longitudinal and cross-sectional FreeSurfer processing streams was assessed using a three-way ANOVA with modality, total follow-up time, and number of observations.

#### 2.3.8. Determinants of longitudinal reliability (III): Global versus regional features

We repeated the same analyses using global summary features – based on the *aseg* and *aparc* parcellations: namely total cortical area and volume, mean cortical thickness, and subcortical and supratentorial volume (without the ventricles). When extracted bilaterally, values were combined. The reliabilities of the global features were compared to the regional estimates of the same modality based on the percentiles.

#### 2.3.9 Reliability of longitudinal brain change: Estimations based on empirical data

We used the multicohort described in the slope variance section (**Supplementary Table 1, Supplementary Figure 5**) to empirically estimate reliability, serving as a validation for the primary analytically derived estimates. This approach ensures reliability is estimated from the same set of individuals, rather than relying on parameter estimates derived from only partially overlapping datasets. Otherwise, analytical derivation is generally preferred as i) provides stronger theoretical justification, ii) is readily applicable to broader research questions and datasets; iii) follows a standardized approach and iv) yields exact estimates. In this analysis, slope variance was considered fixed and estimated as described above while the error variance of the slopes was assessed using the standard error of the slope for each individual and feature. This measure estimates the degree of uncertainty with which an individual slope is estimated. For each feature, the standard error of the slopes were fitted by total follow-up time, the number of observations, and its interaction with cohort as random intercept using generalized linear mixed effects models with a logarithmic link (*glmer*, *lme4* R-package) (Bates et al., 2015). The predictions were corrected by the number of observations as they slightly underestimate the error variance of the slope. See **Supplementary Information** for a detailed description and **Supplementary Figure 6** for visualization.

### 2.4. Supporting app

We provide an interactive tool, powered by *shiny app* (Chang et al., 2022) accompanying this paper to enable visualization and sharing of statistics associated with the manuscript and as an interactive tool for users to explore reliability and sample size estimates of choice for individual differences research using longitudinal data. Along this line, *Longpower* (Iddi and Donohue, 2022)*, LIFESPAN* (Brandmaier et al., 2015), and *ReX* (Xu et al., 2023) are other tools to estimate power and reliability in the context of neuroimaging and longitudinal designs.

## 3. Results

### 3.1 Reliability of longitudinal brain change

See **Figure 2a** for mean effects - across features - of follow-up time and number of observations on longitudinal reliability of MRI change. See the **supporting app** for all the reliability estimates based on feature, follow-up time, and number of observations. A three-way ANOVA with modality, follow-up time, and number of observations showed that ICC was dependent on the main effects of the three parameters (F = 592.7 [η² = 0.26], F = 15565.3 [η² = 0.94], and F = 663.0 [η² = 0.28], respectively; all p < .001). Longer follow-up times led to notable increases in longitudinal reliability, whereas increasing the number of observations led to a more modest increase. Mean reliability across features (i.e., subcortical volume, cortical area, thickness, and volume) was low (ICC = .24 [.10]) with 2 years of follow-up, showing a rapid increase at 4 (ICC = .54 [11]) and 6 (ICC = .72 [.09]) years of follow-up and gradually reaching a plateau with longer follow-up times (ICC = .82 [.07], .87 [.05], .91 [.04] with 8, 10, and 12 years). Increasing the number of time points also increased the reliability, albeit to a minor degree (ΔICC per additional observation = .016). Across all explored features, follow-ups and time-points, subcortical volumetric features (ICC = .78) showed higher reliability than the cortical modalities, while cortical area (ICC = .71) showed slightly higher reliability than both cortical thickness (ICC = .65) and volume (ICC = .67). In addition, ICC was also dependent on the number of observations × follow-up time (F = 14.7, η² = 0.04, p < .001) and the modality × follow-up time two-way interactions (F = 20.9, η² = 0.06, p < .001). The number of observations × follow-up time showed that including more observations with shorter follow-up times led to higher increments in reliability, while increasing follow-up time led to comparatively minor increases in reliability for subcortical features, likely reflecting higher mean values of subcortical features and the consequent plateau effect. See **Supplementary Figure 7** and **Supplementary Tables 7, 8** for the ANOVA visualization and statistics. Significant differences were found across features within modality, especially for subcortical structures (**Figure 2b**). Ventricular and caudate volume showed the highest reliability among subcortical structures, while the pallidum and the amygdala showed the lowest. Cortical features shared a similar regional profile (*ρ* ≈ .34 - .76), with middle cingulate regions, medial and lateral parietal regions, and somatosensory regions showing the highest reliability, while temporal, visual, and orbitofrontal features showed the lowest. Overall, we found a key role of follow-up time on longitudinal reliability, with modest reliability estimates for short study durations, i.e., 2 years, and a comparatively minor impact of increasing the number of observations.

**Figure 2.**
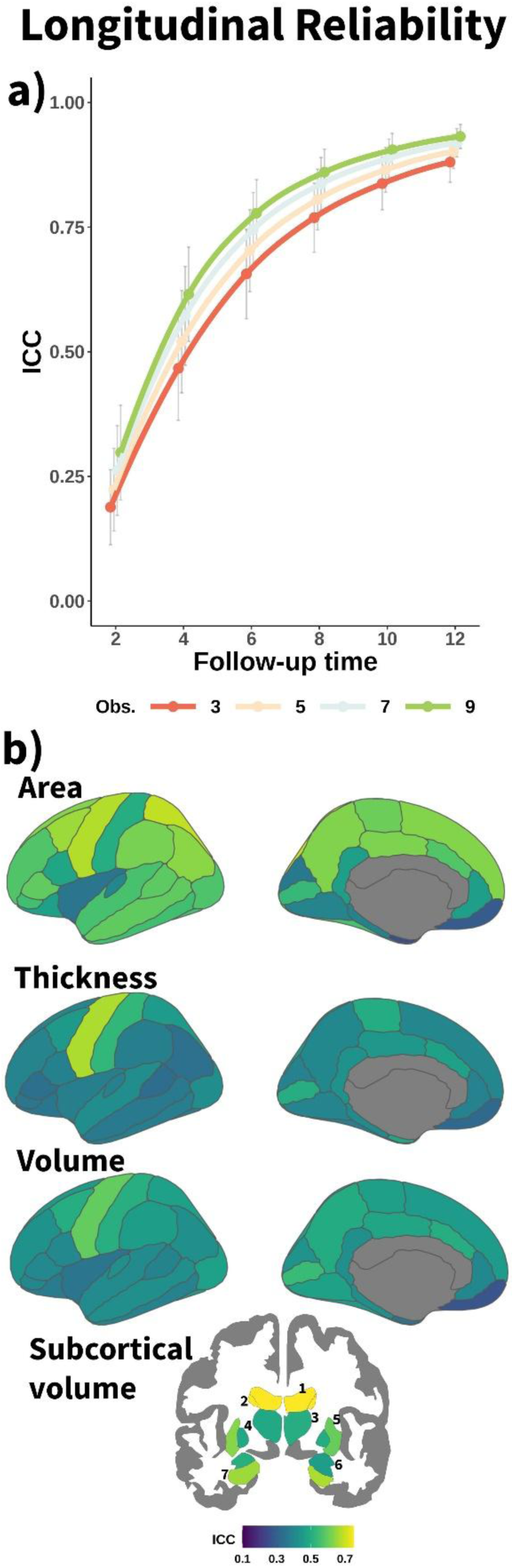
Longitudinal reliability of structural brain features. a) Mean reliability (ICC) of structural brain change across features as a function of total follow-up time and number of equispaced observations. Error bars represent ±1 SD. b) Longitudinal reliability (ICC) for individual structural features, grouped by modality, shown for follow-up time of 4 years and 3 observations. Subcortical features are numbered as follows: 1. Lateral Ventricle, 2. Caudate, 3. Thalamus, 4. Pallidum, 5. Putamen, 6. Amygdala, 7. Hippocampus. Obs. = Number of observations.

### 3.2. Consistency of parameters and reliability estimates across datasets

Next, we assessed the extent to which the parameters used, and the reported reliability estimates are generalizable of legacy MRI datasets and representative of a single of these. We explored whether measurement error (σ_ε_) and slope dispersion parameters (σ_s_) were comparable across datasets using single and mean reliability. See **Supplementary Tables 2** and **4** for information on datasets used for estimating slope dispersion and measurement error, respectively. Mean reliability reflects the degree to which the results are generalizable, i.e., whether they offer a valid characterization of legacy datasets. Single reliability indicates how well the results align with a single dataset, e.g. if one were to use the present results to a single dataset. Reliability of measurement error (σ_ε_) was ICC(2,1) = .65 (CI = .60 - .70) and ICC(2,k) = .92 (CI = .90 - .93) for single and mean measurements, while for slope dispersion (σ_s_) was ICC(2,1) = .82 (CI = .76 - .87) and ICC(2,k) = .97 (CI = .96 - .98). Within modality, cortical area, and subcortical volume showed better reliability estimates of measurement error and slope dispersion. See statistics and visualization in **Supplementary Table 9** and **Supplementary Figure 8**. We also explored the overall and local agreement of the longitudinal reliability estimates using different pairs of error and slope dispersion parameters. The single and mean overall agreements were ICC(2,1) = .84 (.06) and ICC(2,k) = .97 (.01). Within a given follow-up time and number of observations, the single and mean overall agreements were ICC(2,1) = .23 (.14) and ICC(2,k) = .62 (.08). See statistics and visualization in **Supplementary Table 10** and **Supplementary Figure 9**. Overall, measurement error (σ_ε_) and slope dispersion (σ_s_) parameters were comparable across datasets, and the overall reliability pattern was consistent regardless of the cohorts from which the parameters were selected. In contrast, the regional patterns of longitudinal reliability – given a fixed number of observations and study duration - were less stable.

### 3.3 Consequences of longitudinal reliability (I): Sample Size Estimates

Reliability places an upper limit on the maximum detectable effect size, and hence, suboptimal reliability requires larger sample sizes. For each feature, we estimated the impact of follow-up time and number of observations on the sample sizes required to achieve a desirable level of statistical power (80%) at p < 0.05, given a *real* effect size. See **Figure 3** for an illustration with correlation analysis and **Supplementary Table 11** for the accompanying summary statistics. On average, increasing the follow-up time to 4 (from 2) or 6 (from 4) years leads to substantial reductions in required sample size, while a higher number of observations leads also to reductions in required sample size in shorter follow-up designs. Across features, the mean sample size to achieve 80% of power for a real effect size of r = .5 is 154, 60, and 43 individuals following a longitudinal design with 3 observations and 2, 4, or 6 years of follow-up, while the mean sample sizes required for a real effect size of r = .3 and r = .1 are 565, 205, and 139 and 5095, 1860, and 1261 individuals, respectively (**Figure 3a**). In the three cases, a 2-year follow-up requires ≈2.7 and ≈4.0 times more individuals than designs with 4-year and 6-year follow-ups for achieving similar statistical power. Note that the observed correlations will be lower for samples with shorter follow-up design and number of observations due to error-related attenuation. See also estimated sample sizes for Left Hippocampus (**Figure 3b**) and Left Entorhinal thickness (**Figure 3c**). See accompanying statistics in **Supplementary Table 12**. Left Hippocampus, a feature with high longitudinal reliability, shows large reductions of required sample size up to 4 years of follow-up, while the Left Entorhinal thickness, a feature with low reliability, shows how extending the follow-up up to 6 years leads to notable reductions in required sample size. That is, the lower the reliability of longitudinal change, the more benefit one gets, in terms of sample size reduction with longer follow-ups. Extending the duration of short follow-up studies enhances longitudinal reliability and is crucial for reducing sample size estimates. The patterns described above are generally robust across the different features and parameters.

**Figure 3.**
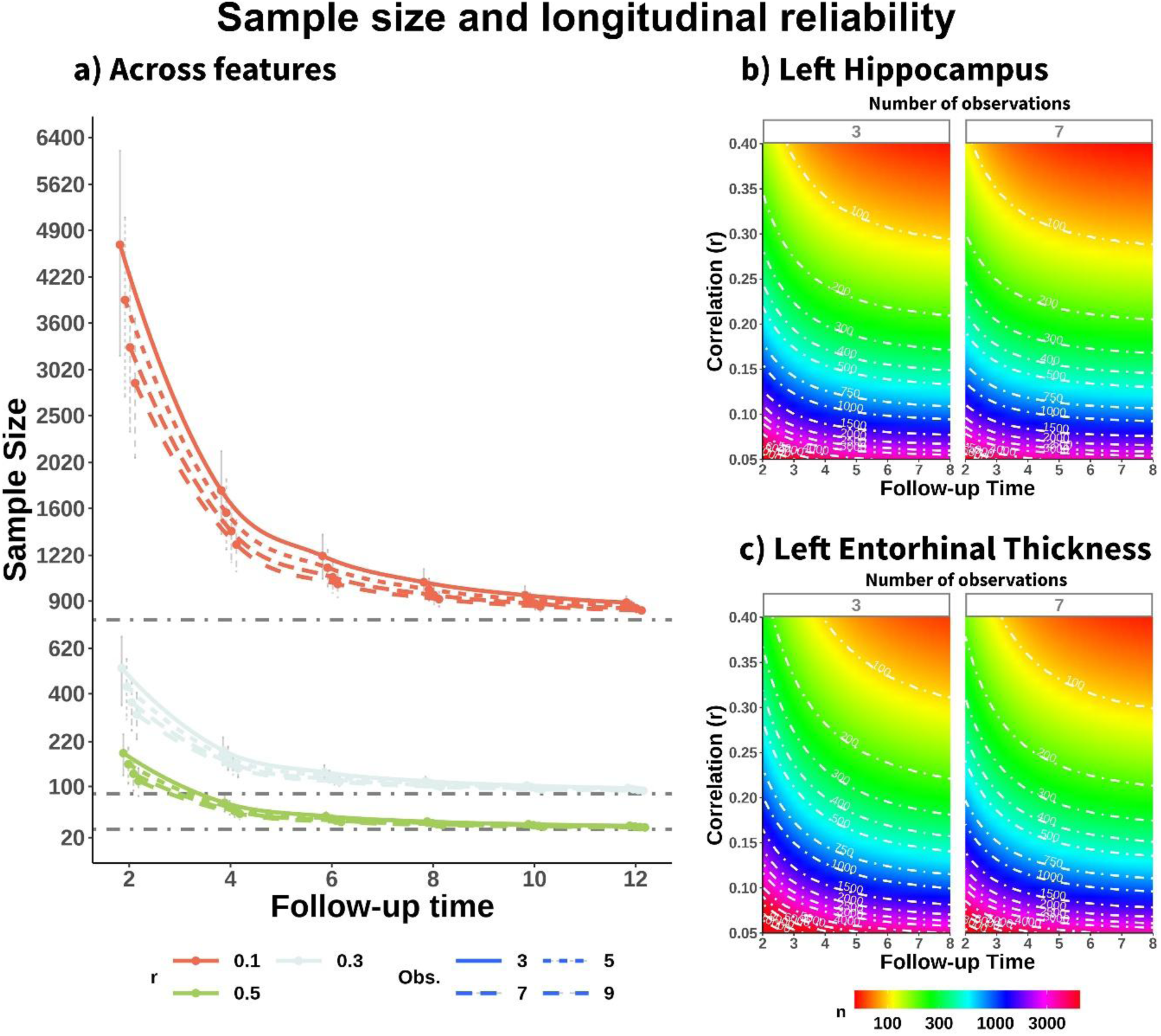
Power analysis for detecting correlations with longitudinal brain change. a) Mean required sample size across structural features (with power = 80%, p < .05) for detecting correlations with longitudinal brain change of small, medium, and large effect sizes (based on conventional guidelines) across different follow-up times and number of observations. Grey horizontal lines are the estimated effect sizes given reliability ICC = 1. Estimated sample size required to detect significant correlations (p < .05, 80% power) between b) the left hippocampus, c) left entorhinal thickness and phenotypes with real correlations ranging from r = 0.05 to 0.4, across different follow-up times and number of observations. For visualization purposes in b) and c), follow-up time is capped at 8 years and the number of observations shown is 3 and 7. r = Pearson’s Correlation. Obs. = Number of observations. Cth = Cortical thickness.

### 3.4. Consequences of longitudinal reliability (II): Misclassification of individual trajectories

Next, we illustrated the effects on the degree of overlap between different trajectories of brain aging given suboptimal longitudinal reliability, by considering a *hypothetical* normal ager, maintainer, and decliner whose brains change 0, −1, and 1 σ_s_ faster relative to the population average and its range of possible *observed* (measured) slopes. We estimated the degree of overlap between distributions using the Bhattacharyya coefficient (BC) and the probability of misclassification (i.e., observing steeper slopes for a normal ager than for a decliner). On average – across features - the distribution of observed values between the normal ager and the decliner (or maintainer) are highly overlapping at short follow-up times, e.g., BC = .97 [0.02] and .89 [0.06] with designs of 2 and 4 years of follow-up, and 3 observations. Increasing the follow-up time sharply decreased the amount of overlap between samples, while increasing the number of observations led to additional reductions in the distribution overlap (**Figure 4a**). This implies a probability of misclassification of p = 0.32 (0.04) and p = .018 (0.05) with designs of 2 and 4 years of follow-up and 3 observations (random classification is p = 0.5). Both the overlap between distributions and the probability of misclassification are much smaller – albeit significant – when comparing a decliner with a maintainer (e.g., BC = 0.89 [0.06], p = 0.18 (0.05) and BC = 0.63 [0.13), p = 0.04 [0.02] with designs of 2 and 4 years of follow-up, 3 observations). See full statistics in **Supplementary Table 13**. See an example of distribution overlap for specific features in **Figure 4b-c**; see corresponding statistics in **Supplementary Table 14**. See visualization and statistics for all features in the **supporting app**. Suboptimal reliability reduces the ability to detect differences in rates of change across individuals, significantly limiting the usefulness of most available longitudinal structural data for making accurate individual-level predictions.

**Figure 4.**
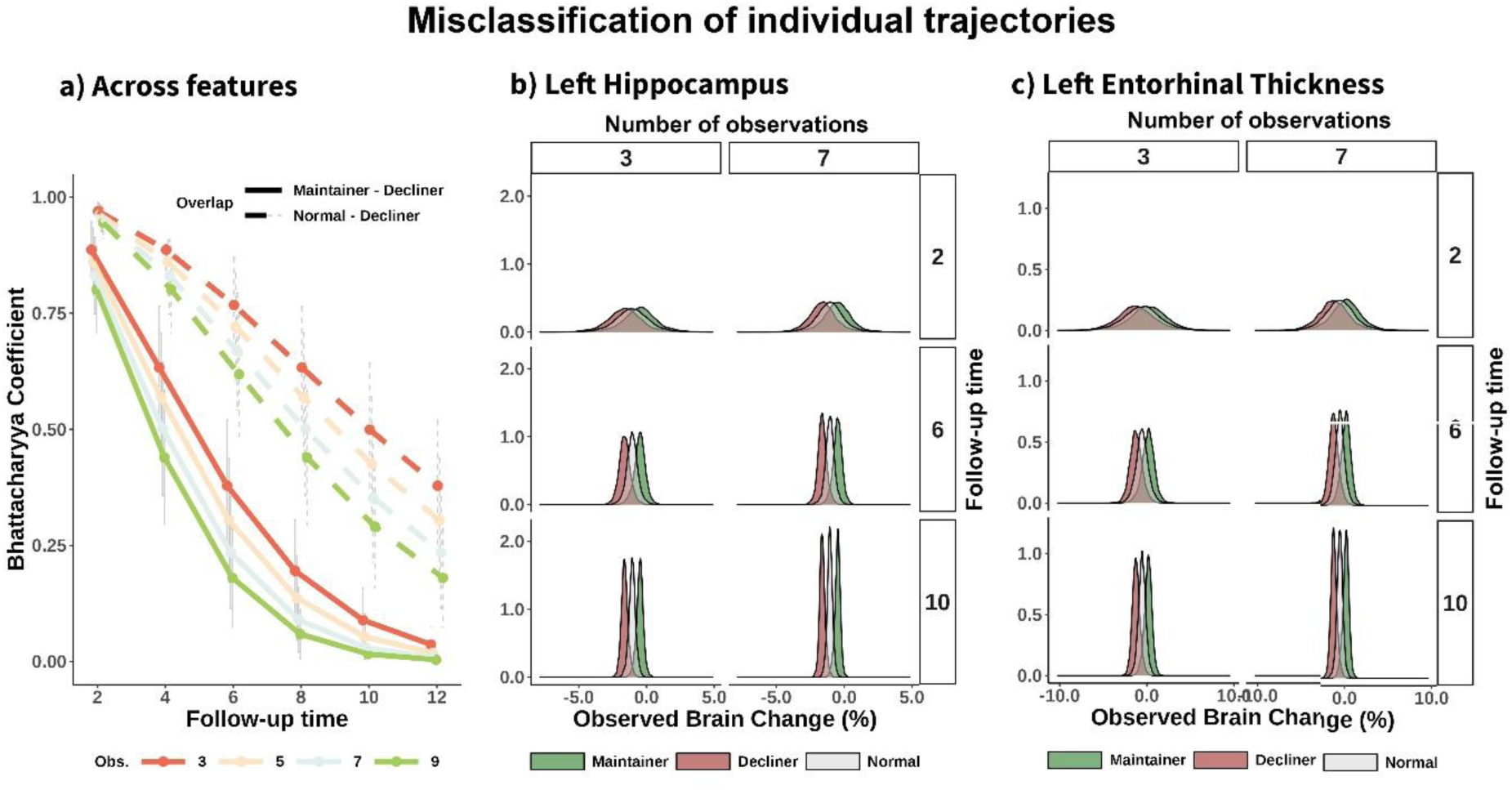
Overlapping between observed estimates of change. a) Mean Bhattacharyya coefficient (BC) across features, quantifying the degree of overlap between two samples as a function of follow-up time and number of observations. The distributions represent possible observed estimates of brain change for three individuals: a normal ager, a maintainer, and a decliner who show decline at an average rate, 1 SD slower, and 1 SD faster, respectively. Overlap in observed brain change distributions for the b) Left Hippocampus, and c) Left Entorhinal Thickness. For b) and c), distributions are shown for 3 and 7 observations and 2, 6, and 10 years of study duration.

### 3.5. Consequences of longitudinal reliability (III): Group membership based on trajectories

Suboptimal longitudinal reliability also has a substantial impact on subgroup classification based on individual trajectories, particularly when combined with observable criteria such as the absence of observed decline. Note that *observed* refers to the measured data while *true* represents the latent, error-free measure. To illustrate this, we simulated brain aging trajectories for individuals, identified those with no *observable* decline over time (*observed brain maintainers*), and estimated the proportion of a) those without *true (latent)* decline over time (*true brain maintainers*) and b) those that had above-average *true* decline (*true brain decliners*). The results showed the following trends: a) the proportion of participants classified as *observed brain maintenance* decreases with longer study duration. b) The proportion of *observed brain maintainers* that are *true brain maintainers* increases with longer study duration, and c) the proportion of *observed brain maintainers* that are *true brain decliners* decreases with longer follow-up times. Increasing the number of observations produced the same trends to a lesser degree. See **Figure 5** for examples of specific features. See statistics in **Supplementary Table 15** and information for the remaining features in the **supporting app**. At short follow-up times, the majority of individuals in which brain maintenance is observed present *true* brain decline. For features with lower reliability, the proportion of individuals with above-average decline showing no decline is also significant (e.g. 15.4% and 9.2% for entorhinal thinning with follow-ups of 2 years and 3 time points). The results showed suboptimal reliability limits the ability to identify biologically meaningful subgroups based on rates of brain aging and increases the risk of misinterpreting data when compared to objective criteria. For instance, many individuals classified as brain maintainers do, in fact, experience true brain decline to varying degrees.

**Figure 5.**
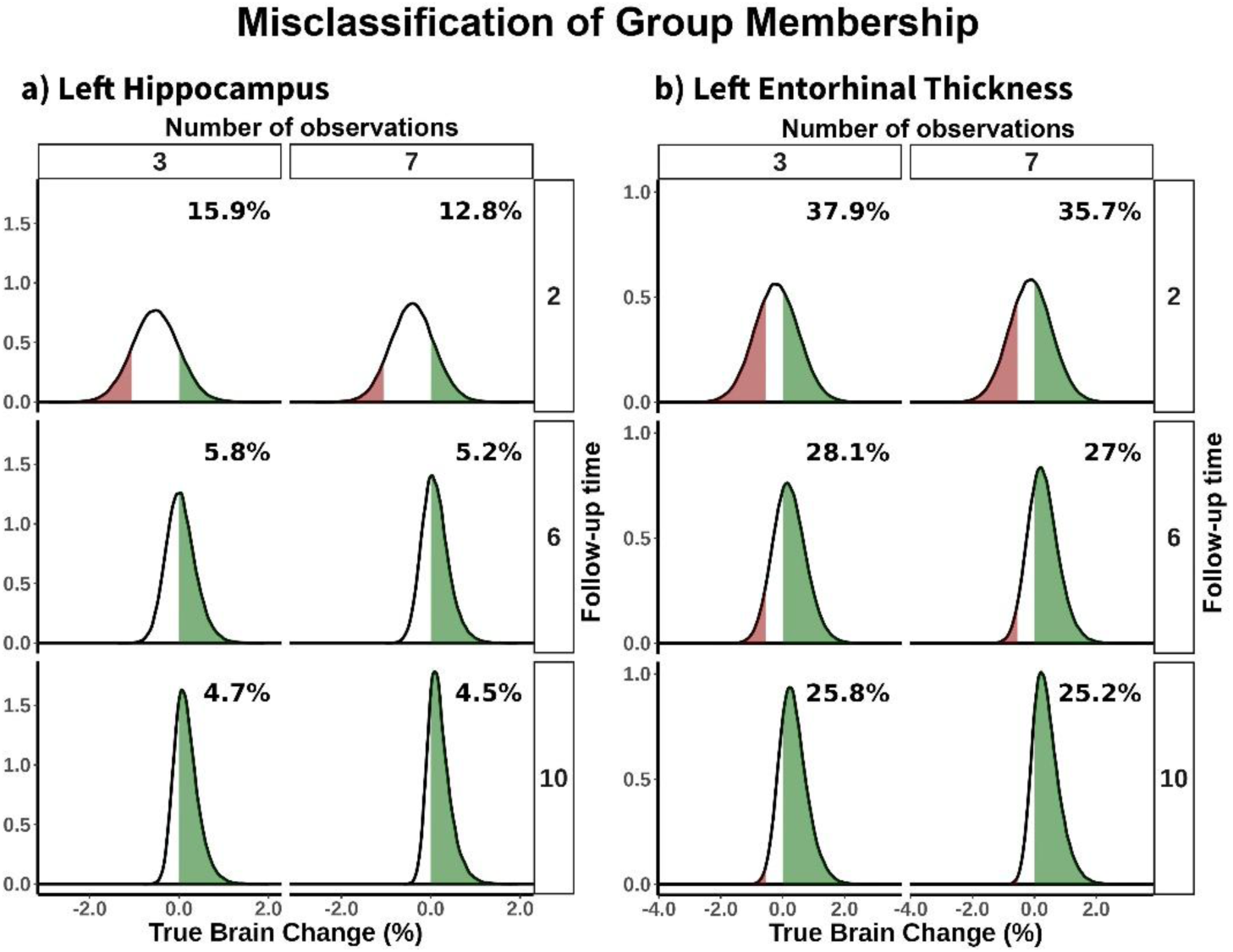
Misclassification based on external criteria. Misclassification of individuals based on an external criterion, i.e., whether they exhibit no brain decline over the duration of the study. The density plots show the distribution of real trajectories of those subjects for whom we would observe no brain decline over time. Green and red fillings represent the proportion of real brain maintenance and real brain decliners, respectively. The text represents the proportion of participants showing no observed brain decline. Shown for the a) Left Hippocampus and b) Left Entorhinal Thickness. Distributions displayed at 3 and 7 observations and 2, 6, and 10 years of study duration.

### 3.6. Determinants of longitudinal reliability (I): Sample characteristics affect longitudinal reliability

Reliability is partially influenced by the variance of the slopes, which itself depends on the sample composition. Younger and healthier samples tend to exhibit more uniform rates of decline compared to samples consisting of older individuals or individuals with pathological load. To illustrate this, we re-estimated the longitudinal reliability using slope variance extracted from a younger and older subsample (cut-off at 60 years). See **Figure 6** for the differences in longitudinal reliability when slope dispersion parameters are extracted either from middle-aged or old adults. Middle-aged adults present less variance of the slopes, and thus worse longitudinal reliability estimates in the (left) hippocampus, and entorhinal cortex than older adults. See statistics in **Supplementary Table 16** and the remaining features in the **supporting app.** Younger, healthier samples require longer follow-up times or a higher number of observations to reach a desired level of longitudinal reliability. Likewise, the consequences of suboptimal reliability, e.g., misclassification, will be more acute in younger datasets.

**Figure 6.**
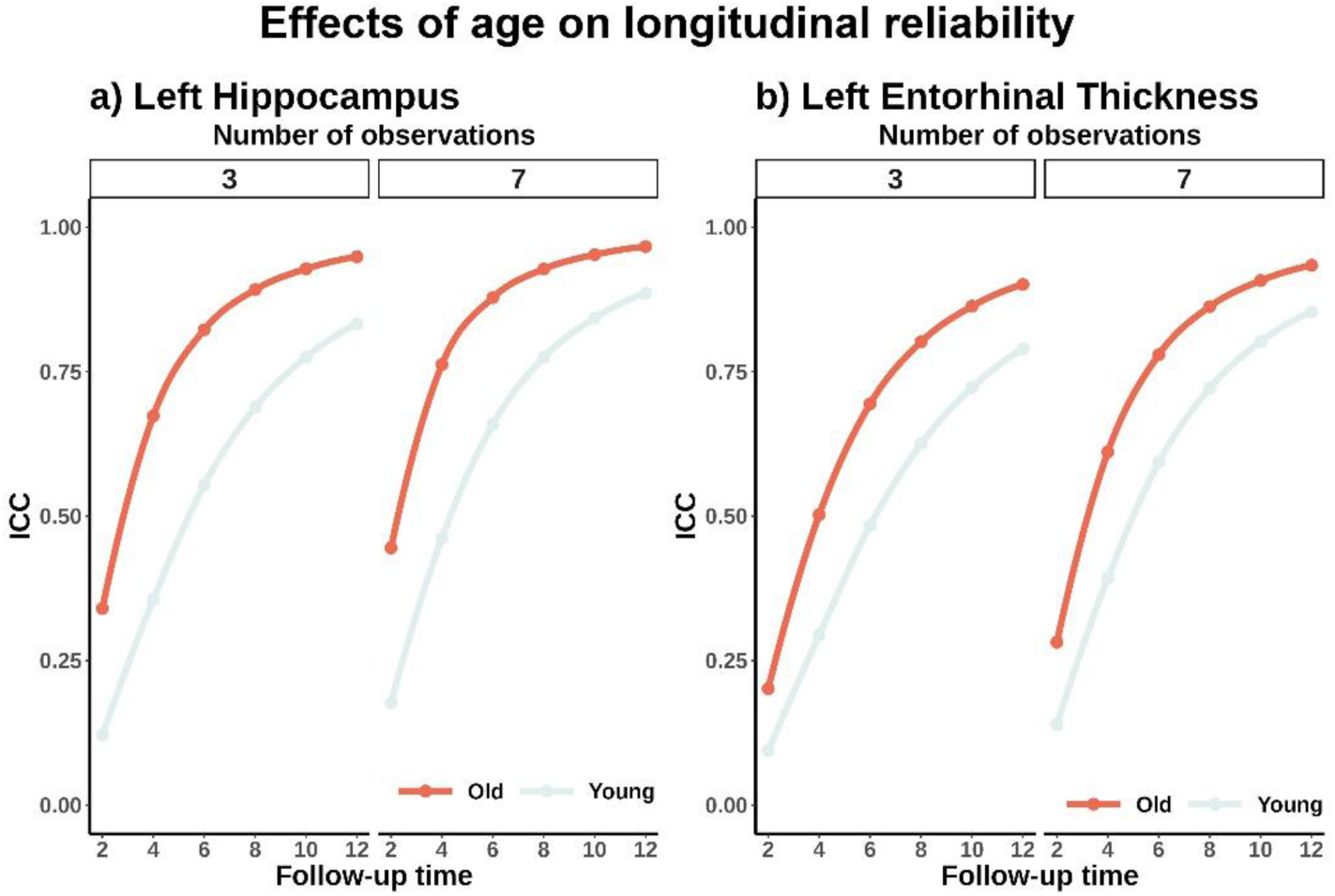
Longitudinal reliability and sample characteristics. Effect of cohort’s age on longitudinal reliability. Older individuals exhibit higher reliability than younger individuals, due to greater variability in slope estimates. Longitudinal reliability as a function of follow-up time, age, and number of observations for the a) Left Hippocampus, and b) Left Entorhinal Thickness. Only distributions at 3 and 7 observations are shown.

### 3.7. Determinants of longitudinal reliability (II): Preprocessing stream

Reliability is partially determined by cross-sectional measurement error. For neuroimaging data, measurement error is in part dependent on the acquisition sequence and the preprocessing pipelines. See **Figure 7a** for mean effects - across features - of follow-up time and number of observations on the reliability of brain change when processed with the cross-sectional FreeSurfer stream. See the **supporting app** for statistics. The overall pattern of longitudinal reliability was similar as shown with data processed using the longitudinal stream (see **Figure 2a**) with main effects of modality, follow-up time, and number of observations (**Supplementary Table 17**). However, longitudinal reliability was notably lower when using data processed with the cross-sectional stream. For the cross-sectional stream, mean reliability across features was ICC = .08 (.06) with 2 years of follow-up increasing to ICC = .24 (.12) and ICC = .41 (.14) after 4 and 6 years of following up and reaching mean reliability values of ICC = .54 [.15], .63 [.14], .71 [.13] with 8, 10, and 12 years). On average, longitudinal reliability was ΔICC = .25 (.12) lower when compared to those derived from the longitudinal FreeSurfer stream (t = 150.9, p < .001). Area and follow-up durations of 4 – 8 showed more pronounced decrements in reliability compared to the longitudinal FreeSurfer stream (**Figure 7b, c**). See **Supplementary Figure 10** for visualization**, Supplementary Table 18** for statistics. Using suboptimal pipelines increase measurement error and thus negatively affects the longitudinal reliability estimates.

**Figure 7.**
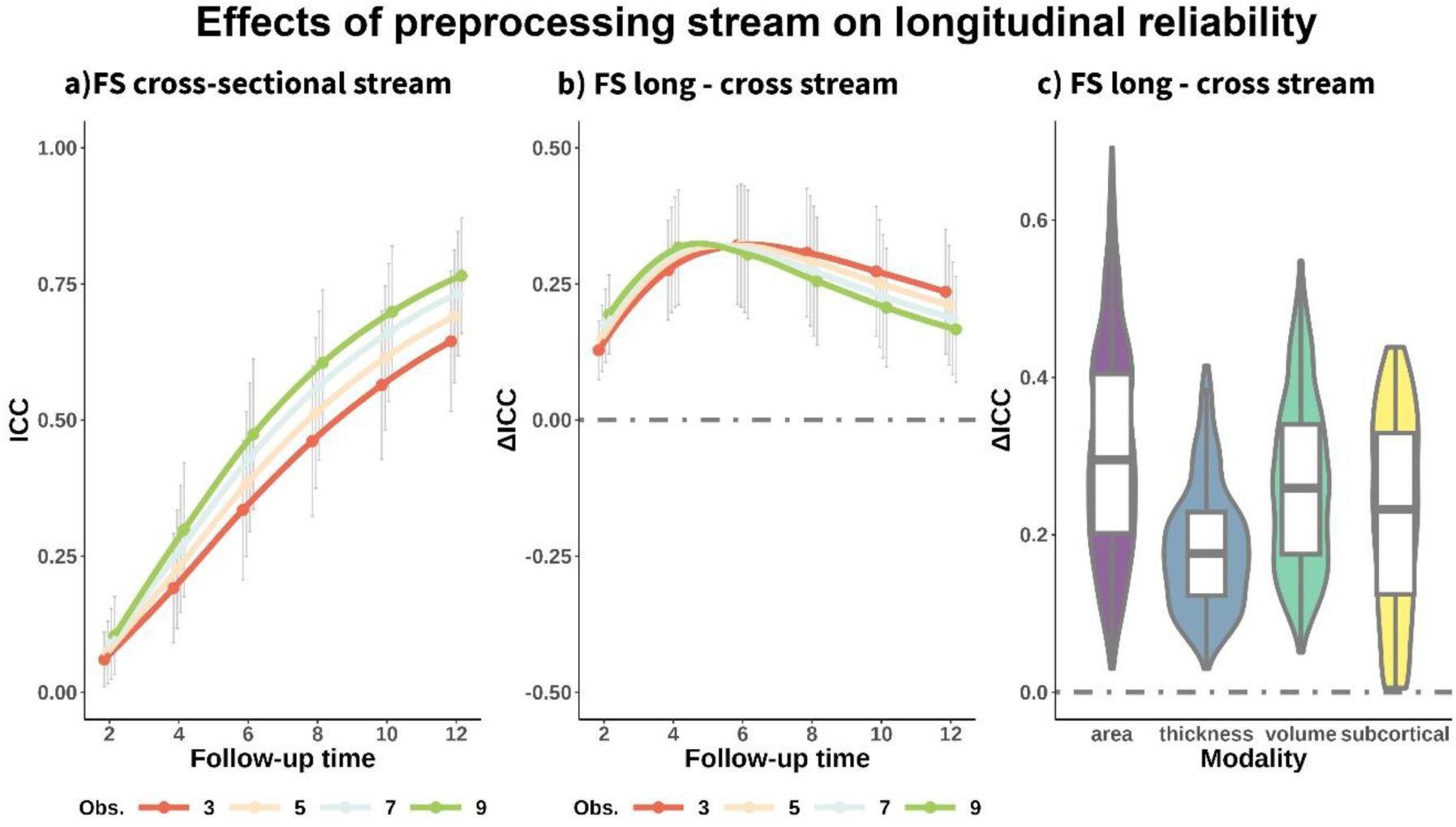
Longitudinal reliability using FreeSurfer cross-sectional stream. Impact of preprocessing stream on longitudinal reliability. a) Mean reliability (ICC) of structural brain change across features as a function of total follow-up time and number of equispaced observations, estimated using the FreeSurfer cross-sectional stream. Mean differences in longitudinal reliability, by b) follow-up time and number of observations and c) modality, between data processed with the longitudinal versus cross-sectional FreeSurfer stream. Positive ΔICC indicates improved reliability estimates when using the longitudinal FreeSurfer Stream. Error bars represent ±1 SD. FS = FreeSurfer. Obs. = Observations.

### 3.8. Determinants of longitudinal reliability (III): Global versus regional features

We computed the longitudinal reliability of global summary variables and compared their reliability with that of regional features within the same modality (**Supplementary Figure 11**). See full longitudinal reliability estimates in the **supporting app**. Mean cortical area and Supratentorial volume showed markedly better longitudinal reliability than most regional estimates being in the 12th and 15th percentiles of their modality. Mean subcortical volume, mean cortical thickness, and mean cortical volume showed average or slightly above-average reliabilities being in the 32nd, 35th, and 51st percentile of their modality. Overall, using common, global variables of brain change did not lead to meaningful improvements in longitudinal reliability.

### 3.9 Reliability of longitudinal brain change: Estimations based on empirical data

See **Figure 8a** for mean effects - across features - of follow-up time and number of observations on longitudinal reliability of MRI change estimated empirically. Note that we assumed a fixed variance of the slope. See the **supporting app** for statistics. The overall pattern of longitudinal reliability was comparable to the main (analytically derived) estimates (see **Figure 2a**) with main effects of modality, follow-up time, and number of observations (**Supplementary Table 19**) as well as a similar mean reliability (ΔICC = −0.03 [.10]). Longitudinal reliability estimated empirically was somewhat higher for study durations of 2 years and lower for study durations of 4 and 6 years; similarly for reliability estimates of cortical area (**Figure 8b, c**). See **Supplementary Figure 12** for visualization and **Supplementary Table 20** for statistics. This analysis provides validation to the main results as the estimation of *true* and *error* variance of the slope are derived from the same dataset.

**Figure 8.**
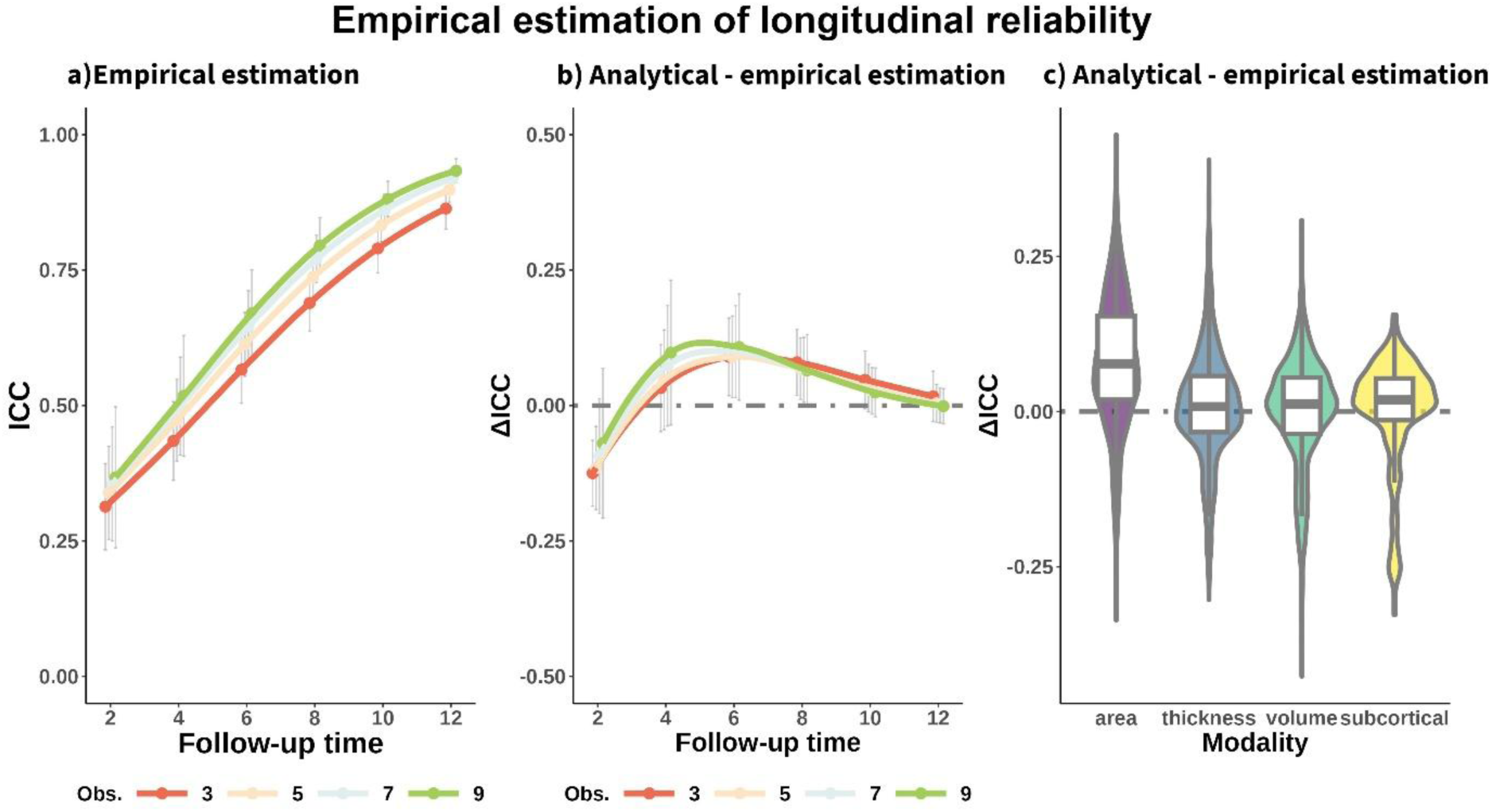
Longitudinal reliability estimated empirically. Error variance of the slopes was estimated from a multi-cohort dataset rather than being analytically-derived from the GRR index. a) Mean reliability (ICC) of structural brain change across features as a function of total follow-up time and number of equispaced observations. Mean differences in longitudinal reliability by b) follow-up time and number of observations and c) modality, between the analytically-derived and the empirical estimations of reliability. Positive ΔICC indicates higher estimates for the analytical derivation of reliability. Error bars represent ±1 SD. Obs. = Observations.

## 4. Discussion

Here, we estimated the reliability of structural brain change in the context of cognitively healthy aging. The results highlighted total follow-up time and measurement error as two crucial factors for detecting individual differences in longitudinal slopes, while the number of observations had, comparatively, a minor effect. These differences in reliability have substantial implications for sample size requirements and the ability to identify the individual trajectories in the context of brain imaging. Subcortical volumes showed higher reliability while the global features did not. The reliability of brain change estimates is also dependent on sample characteristics and the image preprocessing stream. The implications of these results are discussed next.

The main study finding is the notable impact of follow-up time on the reliability of individual differences in brain change. For most features, longitudinal reliability was poor with a measurement interval of 2 years and reached good-to-excellent levels of reliability when the follow-up time was at least 6 or 8 years. Longer study durations reduce error variance and enable a more precise estimation of the individual (linear) slopes. The impact of study duration has been highlighted previously (Brandmaier et al., 2018a, 2015; Fitzmaurice et al., 2012; Markus et al., 2024; Rast and Hofer, 2014). Fitzmaurice and colleagues stated that – for equispaced observations – doubling the length of the measurement interval decreases within-subject variability by a factor of 4, compared to a 29% decrement obtained by increasing a design from 2 to 6 observations. Rast and Hofer (2014) also emphasize the impact of study duration, noting that in short follow-ups, most longitudinal studies lack an adequate foundation for analysis due to the constrains that suboptimal reliability imposes on statistical power to detect associations (see also von Oertzen and Brandmaier, 2013). (Note that longer study durations reduce error variance when one uses yearly estimates of change; rather, if using cumulative change study duration increases interindividual differences in true change). Our results, using structural neuroimaging features are, by and large, in agreement with these landmark studies. To our knowledge, the reliability of brain change has been assessed by a single group by leveraging double-session data (Takao et al., 2022, 2021). These studies showed, on average, low (ICC = .33) and high (ICC = .88) test-retest reliability of brain change for cortical thickness and volumetric change (based on voxel-based morphometry) in cognitively healthy older adults. Takao’s reliability of cortical thinning is consistent with our estimations at 2 years of follow-up, but not those of volumetric data. This approach, however, assumes that the measurements acquired within the same session have independent measurement errors, an assumption that is likely greatly violated when employing signal intensity as a core measure (Elliott et al., 2023b). Although the extent of measurement error dependence is poorly understood, it inflates the reliability estimates, making them an upper bound rather than an accurate reflection of reliability. Using analytical derivation to estimate reliability not only avoids this limitation but also ensures broader applicability to various research questions and datasets. Additionally, it follows a standardized and widely accepted approach that provides a strong theoretical justification and interpretability, aligning reliability estimates with statistical models of measurement.

Higher longitudinal reliability of the subcortical volumes was driven by higher slope variances rather than measurement error, which is in agreement with the existing literature (Hedges et al., 2022; Sele et al., 2020). Slope variances explained regional differences in longitudinal reliability for subcortical volumes, while measurement error was key for cortical area, volume, and across all features when considered together. These results agree with prior evidence of both high variability in the aging trajectories of subcortical structures and the homogenous patterns of cortical decline and longitudinal stability of both cortical area and volume (Parsons et al., 2024; Sele et al., 2021, 2020). As such, cortical differences in longitudinal reliability are partially determined by region size and whether the region lies in the vicinity of air-filled cavities. In any case, caution is required when interpreting regional differences within cortical modalities as they seem dependent on the specific parameter source.

Precomputed global features (e.g., mean cortical thinning) did not present higher longitudinal reliabilities than regional estimates. Global features have both less measurement error and slope variance, which, to some extent, are canceled across brain regions. Increases in longitudinal reliability, will depend thus on how correlated the measurement error and slope variance are throughout the brain. Brain changes in cortical thickness, area, and volume are correlated throughout the brain (within modality) though the strength of these correlations is unclear (Cox et al., 2021; Sele et al., 2021), and the degree to which measurement error co-varies across the brain is unknown. Multivariate indices, e.g., BrainAge, may show better longitudinal reliabilities though performance will also depend on the specific features, samples, and algorithms. However, most research has used this measure cross-sectionally (c.f. Vidal-Pineiro et al., 2021) and efforts are often directed into reducing model error (More et al., 2023) – thus minimizing both true and error variance across individuals – rather than improving its validity and capacity to capture actual brain change. In conclusion, global features do not have better longitudinal reliability, but likely have reduced specificity, and hence lower *true* correlations (Smith et al., 2020). Using global features can be an advantage when modeling strategies account for measurement error throughout the brain.

Reliability has slightly different implications for experimental (e.g., drug trials) and individual differences research (e.g., brain – behavior correlations) (Cronbach, 1957). Experimental research benefits from precision, homogenous groups, and minimizing both within and between-subject variance (Hedge et al., 2018; Parsons et al., 2019; Zimmerman and Zumbo, 2015). Reliability, though, is directly related to statistical power in individual differences research as this aims to maximize the ratio of between vs. within-subject variance (Brandmaier et al., 2018b). In the latter case, reliability places an upper limit on the maximum detectable effect size, with low reliability leading to attenuated correlations and regression estimates or increased uncertainty (depending on if used as dependent or independent variable). Suboptimal reliability leads to either lower statistical power or a need for bigger sample sizes to achieve a desired power - as shown here - and thus results in an increased chance of false negatives (Zuo et al., 2019). Yet, studies with low statistical power also have a higher likelihood of false positives and inflated estimates of effect size, i.e., “*winner’s curse”*, when in the presence of other biases such as undisclosed analytical flexibility (Button et al., 2013; Loken and Gelman, 2017; Simmons et al., 2011). Note that statistical power to detect significant differences will also be affected by the non-brain variables, which often have imperfect reliability and/or validity, such as cognitive scores or cognitive reserve assessments (Fawns-Ritchie and Deary, 2020; Wilson et al., 2019). Suboptimal reliability also impacts the ability to identify a specific individual based on their trajectory and subgroups. Our results caution against making inferences and predictions at an individual level based on brain change for short follow-ups, and against treating model-based classifications as indicative of a *real* phenomenon, e.g., observation of no decline as evidence for brain maintenance. Altogether, suboptimal reliability of brain change hampers our ability to make decisions based on the evidence, to compare studies, and is likely to be an important factor behind the lack of converging evidence of associations between brain change and genetic, environmental, and cognitive factors (Oschwald et al., 2019; Walhovd et al., 2023).

Most studies on longitudinal reliability have focused on study design (Brandmaier et al., 2018a, 2015; Hertzog et al., 2008; Rast and Hofer, 2014; Willett, 1989) and thus differentiate between modifiable (duration, observations, and spacing) and non-modifiable (slope variance and measurement error) features. In neuroimaging, significant attention has also been given to measurement error, as it is partially mitigated through the implementation of advanced imaging sequences and processing pipelines. Yet, most researchers in longitudinal neuroimaging are not involved in the study design, i.e., they carry out secondary data analyses and thus need to consider longitudinal reliability from a different perspective. First, when possible, researchers should estimate reliability and statistical power to minimize the risk of performing underpowered studies, with reliability estimates ideally obtained from the same population (Vacha-Haase et al., 2000). See elsewhere for power analyses and sample size justification (Lakens, 2022). Alternatively, one can use, with caution, estimates presented elsewhere. Second, researchers may select datasets of higher quality. Datasets with longer follow-ups, more observations, and state-of-the-art sequences and scanners are preferable. Selecting participants who have been followed for extended periods may be a possibility, yet it can also reduce the variance of the slope and limit the generalizability of the study, especially in the context of aging. Variance in the slope is generally non-modifiable and constrained by the research question, yet occasionally, it is possible to select populations with high variance, for example, by including older samples or even participants undergoing pathological changes (Nelson and Dannefer, 1992). In any case, slope variance is ultimately limited by the fact that brain trajectories tend to go in the same direction, that is, everybody declines over time, resulting in a small degree of variability in change across the population (Rouder and Haaf, 2018).

Third, one should optimize processing pipelines and control for factors that explain within-subject variation. Here, we showed that FreeSurfer’s longitudinal stream leads to higher estimates of longitudinal reliability compared to the cross-sectional pipeline. This superior performance is attributed to reductions in measurement error (Hedges et al., 2022; Reuter et al., 2012a). In the same vein, *Samseg* (Puonti et al., 2016) may show superior performance to *aseg* subcortical processing (Sederevičius et al., 2021). Further testing is required for other neuroimaging modalities and suites. The only factor commonly considered when accounting for within-subject variation is scanner change. Yet, other factors such as head position, scanner upgrades, and time of the day have been repeatedly suggested to explain both between and within-subject variability (Alfaro-Almagro et al., 2021; Hedges et al., 2022; Karch et al., 2019; Medawar et al., 2021) and are often available - or can be easily estimated - in legacy data. Fourth, one can account for measurement error in the statistical models. Structural equation models (SEMs) are designed to examine relationships between variables (and latent constructs) while accounting for measurement imprecision in observed data. While increasingly popular, SEMs are not the tool of choice for most neuroimaging researchers (cf. Cooper et al., 2019). Alternatively, several methods can account for the effect of measurement error in a range of regression and correlation analyses (e.g. SIMEX, Regression Calibration, attenuation correction) (see overview in Buonaccorsi, 2010) and are implemented in open-source, statistical programming languages such as R (Lederer et al., 2019; Moss, 2019; Nab et al., 2021).

### Considerations

It can be problematic to blindly assume the estimates reported here generalize to other samples, processing pipelines, and acquisition parameters (i.e., reliability induction; Vacha-Haase et al., 2000) as both measurement error and slope variance are likely to differ across datasets. Parsons and colleagues (Parsons et al., 2024) have shown that measurement errors differ by site (scanner- and vendor-specific) while notable pipeline and version-specific measurement errors have been reported here and elsewhere (Hedges et al., 2022; Reuter et al., 2012b; Sederevičius et al., 2021). At the same time, variances of the slopes are also dependent on the population and differ as a function of age – as shown here – or patient inclusion (Jack et al., 2000). Thus, this study is most relevant to others using similar approaches and samples, and caution is required when generalizing the present longitudinal reliability estimates. In any case, we showed that most results are invariant to the specific datasets from which the slope and error parameters were derived (section 3.2), at least when constrained to an image preprocessing pipeline and samples of cognitively healthy adults. As such, we are confident these variations do not affect the key findings of the study: namely, the key effect of follow-up time and the poor reliability of brain change when assessed at short intervals (< 4 years). On the contrary, our conclusions regarding the general pattern of how ICC is affected by the different design choices will likely generalize across a wide variety of measurements (Brandmaier et al., 2024; Rast and Hofer, 2014).

The models used here rely on several assumptions that need to be considered in more detail. I) Older adults present long-term changes in brain structure that can be well captured by linear trajectories. This assumption is presumed – either explicitly or implicitly - in most aging neuroimaging research. While false, as structural brain decline accelerates later in life (Bethlehem et al., 2022; Vidal-Pineiro et al., 2020), nonlinearities are likely to have only a minor effect except in very long follow-up studies. This assumption may be more severely violated in other samples, such as in child and adolescent cohorts. Capturing (individual) non-linear trajectories with accuracy demands markedly more sampling (Ghisletta et al., 2020). Also, it assumes that the researcher is interested in long-term, protracted *aging* changes in brain structure, while state-dependent and short-term variations are considered noise. Within-person variance captures short-term and long-term change and measurement error and, ultimately, what constitutes error and true variance depends on the research question (Karch et al., 2019; Nesselroade, 1991). II) Both slope variance and cross-sectional measurement error are independent of study duration and number of observations. Slope variance can both increase and decrease as a function of study duration. In the context of aging, increasing study duration may lead to reductions in slope variance – and thus of longitudinal reliability – due to sample selectivity, as only an increasingly healthy and motivated subsample of the original participants is retained. Death, disease, and motivational factors affect attrition rates, with missingness – at best – at random. At the same time, some features, particularly ventricular volumes, present higher slope variance with age and consequently, also with longer follow-ups. Higher variance in the follow-ups should lead to better longitudinal reliability (Zorowitz and Niv, 2023). Measurement error, on the other hand, might be higher in older datasets – with longer follow-ups - due to older sequences and software and hardware upgrades.

III) Change and baseline levels are unrelated. Intercept-slope associations can be modeled leading to increments in longitudinal reliability (Brandmaier et al., 2018a). Yet, the relationship between brain structure intercept and change in cognitively healthy aging is often weak to insignificant (Vidal-Pineiro et al., 2021). In our data, a strong association between cross-sectional and longitudinal estimates, was observed only in the left and right lateral ventricles (r = 0.44, 0.41, respectively) (**Supplementary Figure 13**). IV) Brain decline and measurement error are unrelated. While a plausible assumption, there is some evidence that head motion is associated with both brain decline and measurement error (Geerligs et al., 2017; Kemenczky et al., 2022). In our data, we found a significant association between measurement error and age in only one region, reducing, to some extent, this concern (**Supplementary Figure 14**. V) The slopes of brain decline are approximately normally distributed. The scarcely available research shows brain change in normal aging is roughly normally distributed (Fujita et al., 2023), but it ultimately depends on the specific population and feature of interest. In our data, the distribution of brain change is, on average, mildly negatively skewed (**Supplementary Figure 15**). While skewness decreases reliability, the degree of skewness observed for most features is unlikely to have a major impact. Note also, that mean decline, despite in increases with age **(Supplementary Figure 16),** does not influence reliability estimates. Assuming fixed variance of the slopes, we replicated the main findings using empirical data, suggesting some of these concerns have a minor influence on longitudinal reliability.

Note that this study is not designed to optimize longitudinal reliability, rather it intends to be representative of the type of legacy data and study samples frequently analyzed in the aging neuroimaging field. See elsewhere for a priori assessments of longitudinal reliability (Brandmaier et al., 2018a, 2015; Hertzog et al., 2008; Rast and Hofer, 2014; von Oertzen, 2010) were spacing between observations - and measurement error - can be adjusted to maintain the levels of reliability while potentially shortening study duration. Spacing between observations (i.e., the temporal division) is the third factor – together with study duration and number of observations - that influences longitudinal reliability, though it has not formally been studied here because most datasets available have *roughly* equispaced observations. Assuming measurement error is independent between close measurements (see Elliott et al., 2023b; Maclaren et al., 2014), the closer measurements are taken towards the beginning and the end of the study period, the better in terms of longitudinal reliability (Rast and Hofer, 2014; Willett, 1989) (SST can then be estimated using **eq. 3** instead of **eq. 4**). Future datasets can leverage cluster-like acquisitions of rapid MRI scans to boost reliability and power to detect differences (Elliott et al., 2023a). Such approaches hold promise for estimating brain change with relatively short study durations. However, the longitudinal reliability of such datasets is beyond the scope of this study as they rely on considerably different sequences - shorter, noisier -, and introduce correlated errors across measurements acquired within a cluster.

Since slope variance was estimated from *observed* rather than *true* variation, it is overestimated with about 10%. However, it is also underestimated due to attrition bias as the variance is derived from individuals who remain in the study for extended periods. This attrition-related underestimation is more challenging to quantify. This signals higher uncertainty of the reliability estimates, and a likely attenuation of the benefits from extending study duration, as the sample becomes more homogeneous over time. The parameters for variance of the slopes and measurement error were obtained from different datasets, which could be problematic if these parameters were not consistent across datasets. We showed in **section 3.2** these parameters are highly consistent. Further, the empirical reliability estimation, which uses the same datasets to estimate *true* and *error* variance of the slopes shows similar estimations of longitudinal reliability compared to the main, analytically derived, results. Finally, longitudinal reliability refers to the measurement, e.g., *apparent* cortical thickness. Ultimately, validity will not be only constrained by measurement reliability but also by the relationship between our measures and the underlying biological basis (Natu et al., 2019), an association that can be age, and sample-dependent (Vidal-Pineiro et al., 2020).

Measurement error and slope variance are two key parameters of longitudinal reliability, yet are seldom reported in the literature (Hedges et al., 2022; Parsons et al., 2024; Sele et al., 2021). Cross-sectional reliability of brain structure depends on *real* (between-subject) and *error* (within-subjects) variance, yet only the latter is relevant for longitudinal reliability. Thus, cross-sectional and longitudinal reliability are not necessarily related. For example, cortical area shows markedly better cross-sectional reliability than cortical thickness (Hedges et al., 2022; Liem et al., 2015) but similar longitudinal reliability estimates, given that changes in cortical area are – relatively – more homogenous between individuals (Parsons et al., 2024). Previous research has considered *mean* annual change for estimating sample size in the context of longitudinal MRI (Ard and Edland, 2011). Estimation of means requires less data and is widely available in the literature (e.g., Fjell et al., 2009; Fujita et al., 2023; Sele et al., 2021). However, this approach is problematic (Holland et al., 2012), as it sets a specific range of variance based solely on the mean (mean decline and variance are not related in our sample across regions [see **supporting app**]). We encourage further studies to provide measurement error and variance of the slopes, and consequently, we provide these estimates in the **supporting app** for the different subsamples.

In addition to measurement error, slope variance, and corresponding longitudinal reliability estimates, the **supporting app** includes interactive tools for enabling researchers to estimate reliability of their measurements. This tool can be of help to researchers analyzing longitudinal data in other fields or with other populations. This paper is not intended as a critique of previous research; rather it aims to raise awareness of the suboptimal reliabilities met when using longitudinal neuroimaging data and serve as a tool for future research. We hope to draw attention to the assessment and optimization of reliability in longitudinal neuroimaging by providing suggestions and an interactive **supporting app**.

## 5. Conclusions

The results highlight the critical importance of follow-up time for longitudinal reliability, the need for long follow-ups to capture individual brain change in adulthood with precision, and the importance of minimizing measurement error of brain features. These findings call for considering reliability in longitudinal neuroimaging studies and are of relevance not only for the aging neuroimaging community but to researchers and financing bodies invested in understanding the determinants of brain and cognition change over time.

## 6. Data and Code Availability

The raw data were gathered from 21 different datasets. Different agreements are required for each dataset. Most dataset are openly available with prespecified data usage agreements. For some datasets, such as UKB, fees may apply. Requests for Lifebrain cohorts (LCBC, Umeå, UB), COGNORM, and S2C should be submitted to the corresponding principal investigator. See data availability and contact details for all datasets in **Supplementary Table 5** Statistical analyses in this manuscript are available alongside the manuscript and will be made available at https://github.com/LCBC-UiO/Long_Brain_Reliability. All analyses were performed in R 4.2.1. The scripts were run on the Colossus processing cluster, University of Oslo. MRI preprocessing and feature generation scripts were performed with the freely available FreeSurfer software (https://surfer.nmr.mgh.harvard.edu/).

## 7. Author Contributions

**D.V.P.** Conceptualization, Methodology, Formal analysis, Writing - Original Draft; **Ø.S.** Methodology, Software, Formal analysis, Writing - Review & Editing; **M.S.** Resources, Data Curation, Writing - Review & Editing; **I.K.A.** Software, Writing - Review & Editing; **M.A.** Resources, Data Curation, Writing - Review & Editing; **W.B.** Writing - Review & Editing; **D.B-F.** Resources, Writing - Review & Editing; **A.B.** Writing Review & Editing; **A-C.B.** Resources, Writing - Review & Editing; **P.Ga.** Formal analysis; Writing - Review & Editing; **P.Gh.** Writing - Review & Editing; **H.G.** Formal analysis, Writing - Review & Editing; **R.N.H.** Writing - Review & Editing; **R.A.K.** Writing - Review & Editing; **M.K.** Writing - Review & Editing; **S.K.** Writing - Review & Editing; **U.L.** Resources, Writing - Review & Editing; **A.M.M.** Software, Writing Review & Editing; **L.N.** Conceptualization, Resources, Writing - Review & Editing; **J.R.** Data Curation, Writing - Review & Editing; **M.H.S.** Software, Writing - Review & Editing; **C.S-P.** Resources, Writing - Review & Editing; **L-O.W.** Resources, Writing - Review & Editing; **K.B.W.** Conceptualization, Resources, Writing - Review & Editing; **A.M.F.** Conceptualization, Resources, Writing - Review & Editing.

## 8. Funding and Acknowledgements

This work was supported by the Department of Psychology, University of Oslo (to K.B.W., A.M.F.), the Norwegian Research Council (to K.B.W., A.M.F., D.V.P [ES694407]) and the project has received funding from the European Research Council’s Starting Grant scheme under grant agreements 283634, 725025 (to A.M.F.) and 313440 (to K.B.W.). R.N.H. was supported by the UK Medical Research Council [SUAG/046/G101400].

The different sub-studies are supported by different sources. **LCBC**: the Norwegian Research Council (to A.M.F., K.B.W.), and the National Association for Public Health’s dementia research program (A.M.F.). **Umeå (Betula):** a scholar grant from the Knut and Alice Wallenberg (KAW) foundation to L.N. **UB (Barcelona)**: D.B.F. was funded by an ICREA Academia Award. D.B.F, acknowledge the CERCA Programme/Generalitat de Catalunya and is supported by María de Maeztu Unit of Excellence (Institute of Neurosciences, University of Barcelona) MDM-2017-0729, Ministry of Science, Innovation and Universities. L.O.W. and data collection in **COGNORM** is funded by the South-Eastern Norway Regional Health Authorities (#2017095) The Norwegian Health Association (#19536) and by Wellcome Leap’s Dynamic Resilience Program (jointly funded by Temasek Trust) #104617). The funding sources had no role in the study design. Data used in preparation of this article were obtained from the **Alzheimer’s Disease Neuroimaging Initiative (ADNI) database** (adni.loni.usc.edu). The ADNI was launched in 2003 as a public-private partnership, led by Principal Investigator Michael W. Weiner, MD. The primary goal of ADNI has been to test whether serial magnetic resonance imaging (MRI), positron emission tomography (PET), other biological markers, and clinical and neuropsychological assessment can be combined to measure the progression of mild cognitive impairment (MCI) and early Alzheimer’s disease (AD). For up-to-date information, see https://adni.loni.usc.edu/. As such, the investigators within the ADNI contributed to the design and implementation of ADNI and/or provided data but did not participate in analysis or writing of this report. A complete listing of ADNI investigators can be found at: http://adni.loni.usc.edu/wp-content/uploads/how_to_apply/ADNI_Acknowledgement_List.pdf. Data collection and sharing for this project were funded by the ADNI (NIH Grant U01 AG024904). ADNI is funded by the National Institute on Aging, the National Institute of Biomedical Imaging and Bioengineering, and through generous contributions from the following: AbbVie, Alzheimer’s Association; Alzheimer’s Drug Discovery Foundation; Araclon Biotech; BioClinica, Inc.; Biogen; Bristol-Myers Squibb Company; CereSpir, Inc.; Cogstate Eisai Inc.; Elan Pharmaceuticals, Inc.; Eli Lilly and Company; EuroImmun; F. Hoffmann-La Roche Ltd and its affiliated company Genentech, Inc.; Fujirebio; GE Healthcare; IXICO Ltd.; Janssen Alzheimer Immunotherapy Research & Development, LLC.; Johnson & Johnson Pharmaceutical Research & Development LLC.; Lumosity; Lundbeck; Merck & Co., Inc.; Meso Scale Diagnostics, LLC.; NeuroRx Research; Neurotrack Technologies; Novartis Pharmaceuticals Corporation; Pfizer Inc.; Piramal Imaging; Servier; Takeda Pharmaceutical Company; and Transition Therapeutics. The Canadian Institutes of Health Research is providing funds to support ADNI clinical sites in Canada. Private sector contributions are facilitated by the Foundation for the National Institutes of Health (http://www.fnih.org). The grantee organization is the Northern California Institute for Research and Education, and the study is coordinated by the Alzheimer’s Therapeutic Research Institute at the University of Southern California. ADNI data are disseminated by the Laboratory for Neuro Imaging at the University of Southern California. Data used in the preparation of this article was obtained from the **Australian Imaging Biomarkers and Lifestyle flagship study of ageing (AIBL)** funded by the Commonwealth Scientific and Industrial Research Organisation (CSIRO) which was made available at the ADNI database (www.loni.usc.edu/ADNI). The AIBL researchers contributed data but did not participate in analysis or writing of this report. AIBL researchers are listed at www.aibl.csiro.au. Data used in the preparation of this article were obtained from the **Harvard Aging Brain Study** (**HABS** - P01AG036694; https://habs.mgh.harvard.edu). The HABS study was launched in 2010, funded by the National Institute on Aging. and is led by principal investigators Reisa A. Sperling MD and Keith A. Johnson MD at Massachusetts General Hospital/Harvard Medical School in Boston, MA.” **OASIS** data were provided [in part] by **OASIS 3**: Longitudinal Multimodal Neuroimaging: Principal Investigators: T. Benzinger, D. Marcus, J. Morris; NIH P30 AG066444, P50 AG00561, P30 NS09857781, P01 AG026276, P01 AG003991, R01 AG043434, UL1 TR000448, R01 EB009352. AV-45 doses were provided by Avid Radiopharmaceuticals, a wholly owned subsidiary of Eli Lilly and **OASIS-I (test-retest reliability dataset)**: Cross-Sectional: Principal Investigators: D. Marcus, R, Buckner, J, Csernansky J. Morris; P50 AG05681, P01 AG03991, P01 AG026276, R01 AG021910, P20 MH071616, U24 RR021382. **PREVENT-AD** was funded by the Canadian Institutes of Health Research, McGill University, the Fonds de Recherche du Québec – Santé, Alzheimer’s Association, Brain Canada, the Government of Canada, the Canada Fund for Innovation, the Douglas Hospital Research Centre and Foundation, the Levesque Foundation, an unrestricted research grant from Pfizer Canada. Private sector contributions are facilitated by the Development Office of the McGill University Faculty of Medicine and by the Douglas Hospital Research Centre Foundation (http://www.douglas.qc.ca/). **UK Biobank** is generously supported by its founding funders the Wellcome Trust and UK Medical Research Council, as well as the Department of Health, Scottish Government, the Northwest Regional Development Agency, British Heart Foundation and Cancer Research UK. The organisation has over 150 dedicated members of staff, based in multiple locations across the UK. Data were provided [in part] by the **Brain Genomics Superstruct Project (GSP)** of Harvard University and the Massachusetts General Hospital, (Principal Investigators: Randy Buckner, Joshua Roffman, and Jordan Smoller), with support from the Center for Brain Science Neuroinformatics Research Group, the Athinoula A. Martinos Center for Biomedical Imaging, and the Center for Human Genetic Research. 20 individual investigators at Harvard and MGH generously contributed data to GSP Open Access Data Use Terms Version: 2014-Apr-22. **Hangzhou Normal University (HNU)** dataset was made possible by Bing Chen and Qiu Ge from Center for Cognition and Brain Disorders, Hangzhou Normal University, China and supported by the National Natural Science Foundation of China (31070905, 31371134) National Social Science Foundation of China (11AZD119). **Maclaren dataset** was obtained from the OpenNeuro database with accession number is ds000239’; we are are grateful to both. Data used for the **S2C** were in part obtained through the Medical Birth Registry (MBRN) of Norway, the Norwegian Twin Registry (NTR) and the Norwegian Mother, Father and Child Cohort Study (MoBa). MoBa is supported by the Norwegian Ministry of Health and Care Services and the Ministry of Education and Research. We are grateful to all the participating twins and families in Norway who take part in this on-going cohort study. We are grateful to the **Consortium for Reliability and Reproducibility (CoRR)** that supports most test-retest datasets used here, supported by the National Institute on Drug Abuse (NIDA) and the National Natural Science Foundation of China (NSFC) and the work of the Child Mind Institute, the Institute of Psychology, Chinese Academy of Sciences and the Nathan Kline Institute. Part of the computation was performed on the Norwegian high-performance computation resources, sigma2, through the project no. NN9767K.

## 10. References

Alfaro-Almagro, F., McCarthy, P., Afyouni, S., Andersson, J.L.R., Bastiani, M., Miller, K.L., Nichols, T.E., Smith, S.M., 2021. Confound modelling in UK Biobank brain imaging. Neuroimage 224, 117002. 10.1016/j.neuroimage.2020.117002

Allen, M.J., Yen, W.M., 2001. Introduction to measurement theory. Waveland Press.

Appelbaum, M., Cooper, H., Kline, R.B., Mayo-Wilson, E., Nezu, A.M., Rao, S.M., 2018. Journal article reporting standards for quantitative research in psychology: The APA Publications and Communications Board task force report. Am Psychol 73, 3–25. 10.1037/amp0000191

Ard, M.C., Edland, S.D., 2011. Power Calculations for Clinical Trials in Alzheimer’s Disease. J Alzheimers Dis 26, 369–377. 10.3233/JAD-2011-0062

Bates, D., Mächler, M., Bolker, B., Walker, S., 2015. Fitting Linear Mixed-Effects Models Using lme4. Journal of Statistical Software 67, 1–48. 10.18637/jss.v067.i01

Bethlehem, R. a. I., Seidlitz, J., White, S.R., Vogel, J.W., Anderson, K.M., Adamson, C., Adler, S., Alexopoulos, G.S., Anagnostou, E., Areces-Gonzalez, A., Astle, D.E., Auyeung, B., Ayub, M., Bae, J., Ball, G., Baron-Cohen, S., Beare, R., Bedford, S.A., Benegal, V., Beyer, F., Blangero, J., Blesa Cábez, M., Boardman, J.P., Borzage, M., Bosch-Bayard, J.F., Bourke, N., Calhoun, V.D., Chakravarty, M.M., Chen, C., Chertavian, C., Chetelat, G., Chong, Y.S., Cole, J.H., Corvin, A., Costantino, M., Courchesne, E., Crivello, F., Cropley, V.L., Crosbie, J., Crossley, N., Delarue, M., Delorme, R., Desrivieres, S., Devenyi, G.A., Di Biase, M.A., Dolan, R., Donald, K.A., Donohoe, G., Dunlop, K., Edwards, A.D., Elison, J.T., Ellis, C.T., Elman, J.A., Eyler, L., Fair, D.A., Feczko, E., Fletcher, P.C., Fonagy, P., Franz, C.E., Galan-Garcia, L., Gholipour, A., Giedd, J., Gilmore, J.H., Glahn, D.C., Goodyer, I.M., Grant, P.E., Groenewold, N.A., Gunning, F.M., Gur, R.E., Gur, R.C., Hammill, C.F., Hansson, O., Hedden, T., Heinz, A., Henson, R.N., Heuer, K., Hoare, J., Holla, B., Holmes, A.J., Holt, R., Huang, H., Im, K., Ipser, J., Jack, C.R., Jackowski, A.P., Jia, T., Johnson, K.A., Jones, P.B., Jones, D.T., Kahn, R.S., Karlsson, H., Karlsson, L., Kawashima, R., Kelley, E.A., Kern, S., Kim, K.W., Kitzbichler, M.G., Kremen, W.S., Lalonde, F., Landeau, B., Lee, S., Lerch, J., Lewis, J.D., Li, J., Liao, W., Liston, C., Lombardo, M.V., Lv, J., Lynch, C., Mallard, T.T., Marcelis, M., Markello, R.D., Mathias, S.R., Mazoyer, B., McGuire, P., Meaney, M.J., Mechelli, A., Medic, N., Misic, B., Morgan, S.E., Mothersill, D., Nigg, J., Ong, M.Q.W., Ortinau, C., Ossenkoppele, R., Ouyang, M., Palaniyappan, L., Paly, L., Pan, P.M., Pantelis, C., Park, M.M., Paus, T., Pausova, Z., Paz-Linares, D., Pichet Binette, A., Pierce, K., Qian, X., Qiu, J., Qiu, A., Raznahan, A., Rittman, T., Rodrigue, A., Rollins, C.K., Romero-Garcia, R., Ronan, L., Rosenberg, M.D., Rowitch, D.H., Salum, G.A., Satterthwaite, T.D., Schaare, H.L., Schachar, R.J., Schultz, A.P., Schumann, G., Schöll, M., Sharp, D., Shinohara, R.T., Skoog, I., Smyser, C.D., Sperling, R.A., Stein, D.J., Stolicyn, A., Suckling, J., Sullivan, G., Taki, Y., Thyreau, B., Toro, R., Traut, N., Tsvetanov, K.A., Turk-Browne, N.B., Tuulari, J.J., Tzourio, C., Vachon-Presseau, É., Valdes-Sosa, M.J., Valdes-Sosa, P.A., Valk, S.L., van Amelsvoort, T., Vandekar, S.N., Vasung, L., Victoria, L.W., Villeneuve, S., Villringer, A., Vértes, P.E., Wagstyl, K., Wang, Y.S., Warfield, S.K., Warrier, V., Westman, E., Westwater, M.L., Whalley, H.C., Witte, A.V., Yang, N., Yeo, B., Yun, H., Zalesky, A., Zar, H.J., Zettergren, A., Zhou, J.H., Ziauddeen, H., Zugman, A., Zuo, X.N., Bullmore, E.T., Alexander-Bloch, A.F., 2022. Brain charts for the human lifespan. Nature 604, 525–533. 10.1038/s41586-022-04554-y

Brandmaier, A.M., Lindenberger, U., McCormick, E.M., 2024.Optimal Two-Time Point Longitudinal Models for Estimating Individual-Level Change: Asymptotic Insights and Practical Implications.

Brandmaier, A.M., von Oertzen, T., Ghisletta, P., Hertzog, C., Lindenberger, U., 2015. LIFESPAN: A tool for the computer-aided design of longitudinal studies. Frontiers in Psychology 6.

Brandmaier, A.M., von Oertzen, T., Ghisletta, P., Lindenberger, U., Hertzog, C., 2018a. Precision, Reliability, and Effect Size of Slope Variance in Latent Growth Curve Models: Implications for Statistical Power Analysis. Frontiers in Psychology 9.

Brandmaier, A.M., Wenger, E., Bodammer, N.C., Kühn, S., Raz, N., Lindenberger, U., 2018b. Assessing reliability in neuroimaging research through intra-class effect decomposition (ICED). eLife 7, e35718. 10.7554/eLife.35718

Breitner, J.C.S., Poirier, J., Etienne, P.E., Leoutsakos, J.M., 2016. Rationale and Structure for a New Center for Studies on Prevention of Alzheimer’s Disease (StoP-AD). J Prev Alzheimers Dis 3, 236–242. 10.14283/jpad.2016.121

Buonaccorsi, J.P., 2010. Measurement Error, 1st edition. ed. Chapman and Hall/CRC, Boca Raton.

Button, K.S., Ioannidis, J.P.A., Mokrysz, C., Nosek, B.A., Flint, J., Robinson, E.S.J., Munafò, M.R., 2013. Power failure: why small sample size undermines the reliability of neuroscience. Nat Rev Neurosci 14, 365–376. 10.1038/nrn3475

Chang, W., Cheng, J., Allaire, J.J., Sievert, C., Schloerke, B., Xie, Y., Allen, J., McPherson, J., Dipert, A., Borges, B., 2022.shiny: Web Application Framework for R.

Chen, B., Xu, T., Zhou, C., Wang, L., Yang, N., Wang, Z., Dong, H.-M., Yang, Z., Zang, Y.-F., Zuo, X.-N., Weng, X.-C., 2015. Individual Variability and Test-Retest Reliability Revealed by Ten Repeated Resting-State Brain Scans over One Month. PLOS ONE 10, e0144963. 10.1371/journal.pone.0144963

Cooper, S.R., Jackson, J.J., Barch, D.M., Braver, T.S., 2019. Neuroimaging of Individual Differences: A Latent Variable Modeling Perspective. Neurosci Biobehav Rev 98, 29–46. 10.1016/j.neubiorev.2018.12.022

Cox, S.R., Harris, M.A., Ritchie, S.J., Buchanan, C.R., Valdés Hernández, M.C., Corley, J., Taylor, A.M., Madole, J.W., Harris, S.E., Whalley, H.C., McIntosh, A.M., Russ, T.C., Bastin, M.E., Wardlaw, J.M., Deary, I.J., Tucker-Drob, E.M., 2021. Three major dimensions of human brain cortical ageing in relation to cognitive decline across the eighth decade of life. Mol Psychiatry 26, 2651–2662. 10.1038/s41380-020-00975-1

Cronbach, L.J., 1957. The two disciplines of scientific psychology. American Psychologist 12, 671– 684.

Dagley, A., LaPoint, M., Huijbers, W., Hedden, T., McLaren, D.G., Chatwal, J.P., Papp, K.V., Amariglio, R.E., Blacker, D., Rentz, D.M., Johnson, K.A., Sperling, R.A., Schultz, A.P., 2017. Harvard Aging Brain Study: dataset and accessibility. Neuroimage 144, 255–258. 10.1016/j.neuroimage.2015.03.069

Dale, A.M., Fischl, B., Sereno, M.I., 1999. Cortical surface-based analysis. I. Segmentation and surface reconstruction. Neuroimage 9, 179–194. 10.1006/nimg.1998.0395

Daugherty, A.M., Raz, N., 2016. Accumulation of iron in the putamen predicts its shrinkage in healthy older adults: A multi-occasion longitudinal study. Neuroimage 128, 11–20. 10.1016/j.neuroimage.2015.12.045

Desikan, R.S., Ségonne, F., Fischl, B., Quinn, B.T., Dickerson, B.C., Blacker, D., Buckner, R.L., Dale, A.M., Maguire, R.P., Hyman, B.T., Albert, M.S., Killiany, R.J., 2006. An automated labeling system for subdividing the human cerebral cortex on MRI scans into gyral based regions of interest. Neuroimage 31, 968–980. 10.1016/j.neuroimage.2006.01.021

Elliott, M.L., Hanford, L.C., Hamadeh, A., Hilbert, T., Kober, T., Dickerson, B.C., Mair, R.W., Eldaief, M.C., Buckner, R.L., 2023a. Brain morphometry in older adults with and without dementia using extremely rapid structural scans. NeuroImage 276, 120173. 10.1016/j.neuroimage.2023.120173

Elliott, M.L., Nielsen, J.A., Hanford, L.C., Hamadeh, A., Hilbert, T., Kober, T., Dickerson, B.C., Hyman, B.T., Mair, R.W., Eldaief, M.C., Buckner, R.L., 2023b. Precision Brain Morphometry Using Cluster Scanning. 10.1101/2023.12.23.23300492

Ellis, K.A., Bush, A.I., Darby, D., De Fazio, D., Foster, J., Hudson, P., Lautenschlager, N.T., Lenzo, N., Martins, R.N., Maruff, P., Masters, C., Milner, A., Pike, K., Rowe, C., Savage, G., Szoeke, C., Taddei, K., Villemagne, V., Woodward, M., Ames, D., AIBL Research Group, 2009. The Australian Imaging, Biomarkers and Lifestyle (AIBL) study of aging: methodology and baseline characteristics of 1112 individuals recruited for a longitudinal study of Alzheimer’s disease. Int Psychogeriatr 21, 672–687. 10.1017/S1041610209009405

Fawns-Ritchie, C., Deary, I.J., 2020. Reliability and validity of the UK Biobank cognitive tests. PLoS One 15, e0231627. 10.1371/journal.pone.0231627

Fischl, B., Sereno, M.I., Dale, A.M., 1999. Cortical surface-based analysis. II: Inflation, flattening, and a surface-based coordinate system. Neuroimage 9, 195–207. 10.1006/nimg.1998.0396

Fitzmaurice, G.M., Laird, N.M., Ware, J.H., 2012. Applied Longitudinal Analysis. John Wiley & Sons.

Fjell, A.M., Walhovd, K.B., Fennema-Notestine, C., McEvoy, L.K., Hagler, D.J., Holland, D., Brewer, J.B., Dale, A.M., 2009. One-year brain atrophy evident in healthy aging. J. Neurosci. 29, 15223– 15231. 10.1523/JNEUROSCI.3252-09.2009

Fujita, S., Mori, S., Onda, K., Hanaoka, S., Nomura, Y., Nakao, T., Yoshikawa, T., Takao, H., Hayashi, N., Abe, O., 2023. Characterization of Brain Volume Changes in Aging Individuals With Normal Cognition Using Serial Magnetic Resonance Imaging. JAMA Netw Open 6, e2318153. 10.1001/jamanetworkopen.2023.18153

Geerligs, L., Tsvetanov, K.A., Cam-Can, null, Henson, R.N., 2017. Challenges in measuring individual differences in functional connectivity using fMRI: The case of healthy aging. Hum Brain Mapp 38, 4125–4156. 10.1002/hbm.23653

Ghisletta, P., Mason, F., von Oertzen, T., Hertzog, C., Nilsson, L.-G., Lindenberger, U., 2020. On the use of growth models to study normal cognitive aging. International Journal of Behavioral Development 44, 88–96. 10.1177/0165025419851576

Hedge, C., Powell, G., Sumner, P., 2018. The reliability paradox: Why robust cognitive tasks do not produce reliable individual differences. Behav Res Methods 50, 1166–1186. 10.3758/s13428-017-0935-1

Hedges, E.P., Dimitrov, M., Zahid, U., Brito Vega, B., Si, S., Dickson, H., McGuire, P., Williams, S., Barker, G.J., Kempton, M.J., 2022. Reliability of structural MRI measurements: The effects of scan session, head tilt, inter-scan interval, acquisition sequence, FreeSurfer version and processing stream. Neuroimage 246, 118751. 10.1016/j.neuroimage.2021.118751

Hertzog, C., Kramer, A.F., Wilson, R.S., Lindenberger, U., 2008. Enrichment Effects on Adult Cognitive Development: Can the Functional Capacity of Older Adults Be Preserved and Enhanced? Psychol Sci Public Interest 9, 1–65. 10.1111/j.1539-6053.2009.01034.x

Holland, D., McEvoy, L.K., Dale, A.M., Alzheimer’s Disease Neuroimaging Initiative, 2012. Unbiased comparison of sample size estimates from longitudinal structural measures in ADNI. Hum Brain Mapp 33, 2586–2602. 10.1002/hbm.21386

Holmes, A.J., Hollinshead, M.O., O’Keefe, T.M., Petrov, V.I., Fariello, G.R., Wald, L.L., Fischl, B., Rosen, B.R., Mair, R.W., Roffman, J.L., Smoller, J.W., Buckner, R.L., 2015. Brain Genomics Superstruct Project initial data release with structural, functional, and behavioral measures. Sci Data 2, 150031. 10.1038/sdata.2015.31

Iddi, S., Donohue, M.C., 2022. Power and Sample Size for Longitudinal Models in R – The longpower Package and Shiny App. The R Journal 14, 264–282. 10.32614/RJ-2022-022

Idland, A.-V., Sala-Llonch, R., Watne, L.O., Brækhus, A., Hansson, O., Blennow, K., Zetterberg, H., Sørensen, Ø., Walhovd, K.B., Wyller, T.B., Fjell, A.M., 2020. Biomarker profiling beyond amyloid and tau: cerebrospinal fluid markers, hippocampal atrophy, and memory change in cognitively unimpaired older adults. Neurobiol. Aging 93, 1–15. 10.1016/j.neurobiolaging.2020.04.002

Iscan, Z., Jin, T.B., Kendrick, A., Szeglin, B., Lu, H., Trivedi, M., Fava, M., McGrath, P.J., Weissman, M., Kurian, B.T., Adams, P., Weyandt, S., Toups, M., Carmody, T., McInnis, M., Cusin, C., Cooper, C., Oquendo, M.A., Parsey, R.V., DeLorenzo, C., 2015. Test-retest reliability of freesurfer measurements within and between sites: Effects of visual approval process. Hum Brain Mapp 36, 3472–3485. 10.1002/hbm.22856

Jack, C.R., Petersen, R.C., Xu, Y., O’Brien, P.C., Smith, G.E., Ivnik, R.J., Boeve, B.F., Tangalos, E.G., Kokmen, E., 2000. Rates of hippocampal atrophy correlate with change in clinical status in aging and AD. Neurology 55, 484–490. 10.1212/WNL.55.4.484

Kanyongo, G., Brook, G., Kyei-Blankson, L., Gocmen, G., 2007. Reliability and Statistical Power: How Measurement Fallibility Affects Power and Required Sample Sizes for Several Parametric and Nonparametric Statistics. Journal of Modern Applied Statistical Methods 6. 10.22237/jmasm/1177992480

Karch, J.D., Filevich, E., Wenger, E., Lisofsky, N., Becker, M., Butler, O., Mårtensson, J., Lindenberger, U., Brandmaier, A.M., Kühn, S., 2019. Identifying predictors of within-person variance in MRI-based brain volume estimates. Neuroimage 200, 575–589. 10.1016/j.neuroimage.2019.05.030

Kemenczky, P., Vakli, P., Somogyi, E., Homolya, I., Hermann, P., Gál, V., Vidnyánszky, Z., 2022. Effect of head motion-induced artefacts on the reliability of deep learning-based whole-brain segmentation. Sci Rep 12, 1618. 10.1038/s41598-022-05583-3

Lakens, D., 2022. Sample Size Justification. Collabra: Psychology 8, 33267. 10.1525/collabra.33267

LaMontagne, P.J., Benzinger, T.L., Morris, J.C., Keefe, S., Hornbeck, R., Xiong, C., Grant, E., Hassenstab, J., Moulder, K., Vlassenko, A.G., Raichle, M.E., Cruchaga, C., Marcus, D., 2019. OASIS-3: Longitudinal Neuroimaging, Clinical, and Cognitive Dataset for Normal Aging and Alzheimer Disease. 10.1101/2019.12.13.19014902

Lavrakas, P.J., 2008. Encyclopedia of Survey Research Methods. Sage Publications, Inc. 10.4135/9781412963947

Lederer, W., Seibold, H., Küchenhoff, H., Lawrence, C., Brøndum, R.F., 2019. simex: SIMEX- And MCSIMEX-Algorithm for Measurement Error Models.

Liem, F., Mérillat, S., Bezzola, L., Hirsiger, S., Philipp, M., Madhyastha, T., Jäncke, L., 2015. Reliability and statistical power analysis of cortical and subcortical FreeSurfer metrics in a large sample of healthy elderly. Neuroimage 108, 95–109. 10.1016/j.neuroimage.2014.12.035

Loken, E., Gelman, A., 2017. Measurement error and the replication crisis. Science 355, 584–585. 10.1126/science.aal3618

Maclaren, J., Han, Z., Vos, S.B., Fischbein, N., Bammer, R., 2014. Reliability of brain volume measurements: A test-retest dataset. Sci Data 1, 140037. 10.1038/sdata.2014.37

Madan, C.R., Kensinger, E.A., 2017. Test–retest reliability of brain morphology estimates. Brain Inform 4, 107–121. 10.1007/s40708-016-0060-4

Marcus, D.S., Wang, T.H., Parker, J., Csernansky, J.G., Morris, J.C., Buckner, R.L., 2007. Open Access Series of Imaging Studies (OASIS): cross-sectional MRI data in young, middle aged, nondemented, and demented older adults. J Cogn Neurosci 19, 1498–1507. 10.1162/jocn.2007.19.9.1498

Markus, A.M., Lindenberger, U., McCormick, E.M., 2024. Optimal Two-Time Point Longitudinal Models for Estimating Individual-Level Change: Asymptotic Insights and Practical Implications. 10.31234/osf.io/6dc8e

McGraw, K.O., Wong, S.P., 1996. Forming inferences about some intraclass correlation coefficients. Psychological Methods 1, 30–46. 10.1037/1082-989X.1.1.30

Medawar, E., Thieleking, R., Manuilova, I., Paerisch, M., Villringer, A., Witte, A.V., Beyer, F., 2021. Estimating the effect of a scanner upgrade on measures of grey matter structure for longitudinal designs. PLOS ONE 16, e0239021. 10.1371/journal.pone.0239021

Milham, M.P., Craddock, R.C., Son, J.J., Fleischmann, M., Clucas, J., Xu, H., Koo, B., Krishnakumar, A., Biswal, B.B., Castellanos, F.X., Colcombe, S., Di Martino, A., Zuo, X.-N., Klein, A., 2018. Assessment of the impact of shared brain imaging data on the scientific literature. Nat Commun 9, 2818. 10.1038/s41467-018-04976-1

Miller, K.L., Alfaro-Almagro, F., Bangerter, N.K., Thomas, D.L., Yacoub, E., Xu, J., Bartsch, A.J., Jbabdi, S., Sotiropoulos, S.N., Andersson, J.L.R., Griffanti, L., Douaud, G., Okell, T.W., Weale, P., Dragonu, I., Garratt, S., Hudson, S., Collins, R., Jenkinson, M., Matthews, P.M., Smith, S.M., 2016. Multimodal population brain imaging in the UK Biobank prospective epidemiological study. Nature Neuroscience 19, 1523–1536. 10.1038/nn.4393

More, S., Antonopoulos, G., Hoffstaedter, F., Caspers, J., Eickhoff, S.B., Patil, K.R., 2023. Brain-age prediction: A systematic comparison of machine learning workflows. NeuroImage 270, 119947. 10.1016/j.neuroimage.2023.119947

Moss, J., 2019. Correcting for attenuation due to measurement error. 10.48550/arXiv.1911.01576

Mowinckel, A.M., Vidal-Piñeiro, D., 2020. Visualization of Brain Statistics With R Packages ggseg and ggseg3d. Advances in Methods and Practices in Psychological Science 3, 466–483. 10.1177/2515245920928009

Mueller, S.G., Weiner, M.W., Thal, L.J., Petersen, R.C., Jack, C., Jagust, W., Trojanowski, J.Q., Toga, A.W., Beckett, L., 2005. The Alzheimer’s disease neuroimaging initiative. Neuroimaging Clin. N. Am. 15, 869–877, xi–xii. 10.1016/j.nic.2005.09.008

Nab, L., Smeden, M. van, Keogh, R.H., Groenwold, R.H.H., 2021. Mecor: An R package for measurement error correction in linear regression models with a continuous outcome. Computer Methods and Programs in Biomedicine 208, 106238. 10.1016/j.cmpb.2021.106238

Natu, V.S., Gomez, J., Barnett, M., Jeska, B., Kirilina, E., Jaeger, C., Zhen, Z., Cox, S., Weiner, K.S., Weiskopf, N., Grill-Spector, K., 2019. Apparent thinning of human visual cortex during childhood is associated with myelination. Proceedings of the National Academy of Sciences 116, 20750–20759. 10.1073/pnas.1904931116

Nelson, E.A., Dannefer, D., 1992. Aged heterogeneity: fact or fiction? The fate of diversity in gerontological research. Gerontologist 32, 17–23. 10.1093/geront/32.1.17

Nesselroade, J.R., 1991. Interindividual differences in intraindividual change, in: Best Methods for the Analysis of Change: Recent Advances, Unanswered Questions, Future Directions. American Psychological Association, Washington, DC, US, pp. 92–105. 10.1037/10099-006

Nyberg, L., Salami, A., Andersson, M., Eriksson, J., Kalpouzos, G., Kauppi, K., Lind, J., Pudas, S., Persson, J., Nilsson, L.-G., 2010. Longitudinal evidence for diminished frontal cortex function in aging. Proc. Natl. Acad. Sci. U.S.A. 107, 22682–22686. 10.1073/pnas.1012651108

Orban, P., Madjar, C., Savard, M., Dansereau, C., Tam, A., Das, S., Evans, A.C., Rosa-Neto, P., Breitner, J.C.S., Bellec, P., 2015. Test-retest resting-state fMRI in healthy elderly persons with a family history of Alzheimer’s disease. Sci Data 2, 150043. 10.1038/sdata.2015.43

Oschwald, J., Guye, S., Liem, F., Rast, P., Willis, S., Röcke, C., Jäncke, L., Martin, M., Mérillat, S., 2019. Brain structure and cognitive ability in healthy aging: a review on longitudinal correlated change. Rev Neurosci 31, 1–57. 10.1515/revneuro-2018-0096

Parsons, S., Brandmaier, A.M., Lindenberger, U., Kievit, R., 2024. Longitudinal stability of cortical grey matter measures varies across brain regions, imaging metrics, and testing sites in the ABCD study. Imaging Neuroscience. 10.1162/imag_a_00086

Parsons, S., Kruijt, A.-W., Fox, E., 2019. Psychological Science Needs a Standard Practice of Reporting the Reliability of Cognitive-Behavioral Measurements. Advances in Methods and Practices in Psychological Science 2, 378–395. 10.1177/2515245919879695

Puonti, O., Iglesias, J.E., Van Leemput, K., 2016. Fast and sequence-adaptive whole-brain segmentation using parametric Bayesian modeling. NeuroImage 143, 235–249. 10.1016/j.neuroimage.2016.09.011

R Core Team, 2023. R: A Language and Environment for Statistical Computing. R Foundation for Statistical Computing, Vienna, Austria.

Rajaram, S., Valls-Pedret, C., Cofán, M., Sabaté, J., Serra-Mir, M., Pérez-Heras, A.M., Arechiga, A., Casaroli-Marano, R.P., Alforja, S., Sala-Vila, A., Doménech, M., Roth, I., Freitas-Simoes, T.M., Calvo, C., López-Illamola, A., Haddad, E., Bitok, E., Kazzi, N., Huey, L., Fan, J., Ros, E., 2017. The Walnuts and Healthy Aging Study (WAHA): Protocol for a Nutritional Intervention Trial with Walnuts on Brain Aging. Front Aging Neurosci 8. 10.3389/fnagi.2016.00333

Rast, P., Hofer, S.M., 2014. Longitudinal design considerations to optimize power to detect variances and covariances among rates of change: simulation results based on actual longitudinal studies. Psychol Methods 19, 133–154. 10.1037/a0034524

Raz, N., Lindenberger, U., 2011. Only time will tell: cross-sectional studies offer no solution to the age-brain-cognition triangle: comment on Salthouse (2011). Psychol Bull 137, 790–795. 10.1037/a0024503

Raz, N., Yang, Y., Dahle, C.L., Land, S., 2012. Volume of White Matter Hyperintensities in Healthy Adults: Contribution of Age, Vascular Risk Factors, and Inflammation-Related Genetic Variants. Biochim Biophys Acta 1822, 361–369. 10.1016/j.bbadis.2011.08.007

Reuter, M., Schmansky, N.J., Rosas, H.D., Fischl, B., 2012a. Within-subject template estimation for unbiased longitudinal image analysis. Neuroimage 61, 1402–1418. 10.1016/j.neuroimage.2012.02.084

Reuter, M., Schmansky, N.J., Rosas, H.D., Fischl, B., 2012b. Within-subject template estimation for unbiased longitudinal image analysis. NeuroImage 61, 1402–1418. 10.1016/j.neuroimage.2012.02.084

Rouder, J.N., Haaf, J.M., 2018. Power, Dominance, and Constraint: A Note on the Appeal of Different Design Traditions. Advances in Methods and Practices in Psychological Science 1, 19–26. 10.1177/2515245917745058

Sederevičius, D., Vidal-Piñeiro, D., Sørensen, Ø., van Leemput, K., Iglesias, J.E., Dalca, A.V., Greve, D.N., Fischl, B., Bjørnerud, A., Walhovd, K.B., Fjell, A.M., Alzheimers Disease Neuroimaging Initiative, 2021. Reliability and sensitivity of two whole-brain segmentation approaches included in FreeSurfer - ASEG and SAMSEG. Neuroimage 237, 118113. 10.1016/j.neuroimage.2021.118113

Sele, S., Liem, F., Mérillat, S., Jäncke, L., 2021. Age-related decline in the brain: a longitudinal study on inter-individual variability of cortical thickness, area, volume, and cognition. Neuroimage 240, 118370. 10.1016/j.neuroimage.2021.118370

Sele, S., Liem, F., Mérillat, S., Jäncke, L., 2020. Decline Variability of Cortical and Subcortical Regions in Aging: A Longitudinal Study. Front Hum Neurosci 14, 363. 10.3389/fnhum.2020.00363

Shrout, P.E., Fleiss, J.L., 1979. Intraclass correlations: Uses in assessing rater reliability. Psychological Bulletin 86, 420–428. 10.1037/0033-2909.86.2.420

Simmons, J.P., Nelson, L.D., Simonsohn, U., 2011. False-Positive Psychology: Undisclosed Flexibility in Data Collection and Analysis Allows Presenting Anything as Significant. Psychol Sci 22, 1359–1366. 10.1177/0956797611417632

Simpson, G.L., 2024.gratia: Graceful ggplot-Based Graphics and Other Functions for GAMs Fitted using mgcv.

Smith, S.M., Elliott, L.T., Alfaro-Almagro, F., McCarthy, P., Nichols, T.E., Douaud, G., Miller, K.L., 2020. Brain aging comprises many modes of structural and functional change with distinct genetic and biophysical associations. eLife 9, e52677. 10.7554/eLife.52677

Spearman, C., 1904. The proof and measurement of association between two things. The American Journal of Psychology 15, 72–101. 10.2307/1412159

Takao, H., Amemiya, S., Abe, O., Alzheimer’s Disease Neuroimaging Initiative, 2022. Reproducibility of Longitudinal Changes in Cortical Thickness Determined by Surface-Based Morphometry Between Non-Accelerated and Accelerated MR Imaging. J Magn Reson Imaging 55, 1151–1160. 10.1002/jmri.27929

Takao, H., Amemiya, S., Abe, O., Alzheimer’s Disease Neuroimaging Initiative, 2021. Reproducibility of Brain Volume Changes in Longitudinal Voxel-Based Morphometry Between Non-Accelerated and Accelerated Magnetic Resonance Imaging. J Alzheimers Dis 83, 281–290. 10.3233/JAD-210596

Tremblay-Mercier, J., Madjar, C., Das, S., Pichet Binette, A., Dyke, S.O.M., Étienne, P., Lafaille-Magnan, M.-E., Remz, J., Bellec, P., Louis Collins, D., Natasha Rajah, M., Bohbot, V., Leoutsakos, J.-M., Iturria-Medina, Y., Kat, J., Hoge, R.D., Gauthier, S., Tardif, C.L., Mallar Chakravarty, M., Poline, J.-B., Rosa-Neto, P., Evans, A.C., Villeneuve, S., Poirier, J., Breitner, J.C.S., PREVENT-AD Research Group, 2021. Open science datasets from PREVENT-AD, a longitudinal cohort of pre-symptomatic Alzheimer’s disease. Neuroimage Clin 31, 102733. 10.1016/j.nicl.2021.102733

Vacha-Haase, T., Kogan, L.R., Thompson, B., 2000. Sample compositions and variabilities in published studies versus those in test manuals: Validity of score reliability inductions. Educational and Psychological Measurement 60, 509–522. 10.1177/00131640021970682

Vidal-Piñeiro, D., Martin-Trias, P., Arenaza-Urquijo, E.M., Sala-Llonch, R., Clemente, I.C., Mena-Sánchez, I., Bargalló, N., Falcón, C., Pascual-Leone, Á., Bartrés-Faz, D., 2014. Task-dependent activity and connectivity predict episodic memory network-based responses to brain stimulation in healthy aging. Brain Stimul 7, 287–296. 10.1016/j.brs.2013.12.016

Vidal-Pineiro, D., Parker, N., Shin, J., French, L., Grydeland, H., Jackowski, A.P., Mowinckel, A.M., Patel, Y., Pausova, Z., Salum, G., Sørensen, Ø., Walhovd, K.B., Paus, T., Fjell, A.M., 2020. Cellular correlates of cortical thinning throughout the lifespan. Scientific Reports 10, 21803. 10.1038/s41598-020-78471-3

Vidal-Pineiro, D., Wang, Y., Krogsrud, S.K., Amlien, I.K., Baaré, W.F., Bartres-Faz, D., Bertram, L., Brandmaier, A.M., Drevon, C.A., Düzel, S., Ebmeier, K., Henson, R.N., Junqué, C., Kievit, R.A., Kühn, S., Leonardsen, E., Lindenberger, U., Madsen, K.S., Magnussen, F., Mowinckel, A.M., Nyberg, L., Roe, J.M., Segura, B., Smith, S.M., Sørensen, Ø., Suri, S., Westerhausen, R., Zalesky, A., Zsoldos, E., Walhovd, K.B., Fjell, A., 2021. Individual variations in ‘brain age’ relate to early-life factors more than to longitudinal brain change. eLife 10, e69995. 10.7554/eLife.69995

von Oertzen, T., 2010. Power equivalence in structural equation modelling. Br J Math Stat Psychol 63, 257–272. 10.1348/000711009X441021

von Oertzen, T., Brandmaier, A.M., 2013. Optimal study design with identical power: an application of power equivalence to latent growth curve models. Psychol Aging 28, 414–428. 10.1037/a0031844

Walhovd, K.B., Fjell, A.M., Westerhausen, R., Nyberg, L., Ebmeier, K.P., Lindenberger, U., Bartrés-Faz, D., Baaré, W.F.C., Siebner, H.R., Henson, R., Drevon, C.A., Strømstad Knudsen, G.P., Ljøsne, I.B., Penninx, B.W.J.H., Ghisletta, P., Rogeberg, O., Tyler, L., Bertram, L., Lifebrain Consortium, 2018. Healthy minds 0-100 years: Optimising the use of European brain imaging cohorts (“Lifebrain”). Eur. Psychiatry 50, 47–56. 10.1016/j.eurpsy.2017.12.006

Walhovd, K.B., Krogsrud, S.K., Amlien, I.K., Bartsch, H., Bjørnerud, A., Due-Tønnessen, P., Grydeland, H., Hagler, D.J., Håberg, A.K., Kremen, W.S., Ferschmann, L., Nyberg, L., Panizzon, M.S., Rohani, D.A., Skranes, J., Storsve, A.B., Sølsnes, A.E., Tamnes, C.K., Thompson, W.K., Reuter, C., Dale, A.M., Fjell, A.M., 2016. Neurodevelopmental origins of lifespan changes in brain and cognition. Proc. Natl. Acad. Sci. U.S.A. 113, 9357–9362. 10.1073/pnas.1524259113

Walhovd, K.B., Krogsrud, S.K., Amlien, I.K., Sørensen, Ø., Wang, Y., Bråthen, A.C.S., Overbye, K., Kransberg, J., Mowinckel, A.M., Magnussen, F., Herud, M., Håberg, A.K., Fjell, A.M., Vidal-Piñeiro, D., 2024. Back to the future: omnipresence of fetal influence on the human brain through the lifespan. eLife 12. 10.7554/eLife.86812.2

Walhovd, K.B., Lövden, M., Fjell, A.M., 2023. Timing of lifespan influences on brain and cognition. Trends in Cognitive Sciences 27, 901–915. 10.1016/j.tics.2023.07.001

Wickham, H., 2016. ggplot2: Elegant Graphics for Data Analysis, 2nd ed, Use R! Springer International Publishing. 10.1007/978-3-319-24277-4

Willett, J.B., 1989. Some Results on Reliability for the Longitudinal Measurement of Change: Implications for the Design of Studies of Individual Growth. Educational and Psychological Measurement 49, 587–602. 10.1177/001316448904900309

Wilson, R.S., Yu, L., Lamar, M., Schneider, J.A., Boyle, P.A., Bennett, D.A., 2019. Education and cognitive reserve in old age. Neurology 92, e1041–e1050. 10.1212/WNL.0000000000007036

Wood, S.N., 2017. Generalized Additive Models: An Introduction with R, 2nd ed. Chapman and Hall/CRC.

Xu, T., Kiar, G., Cho, J.W., Bridgeford, E.W., Nikolaidis, A., Vogelstein, J.T., Milham, M.P., 2023. ReX: an integrative tool for quantifying and optimizing measurement reliability for the study of individual differences. Nat Methods 20, 1025–1028. 10.1038/s41592-023-01901-3

Zimmerman, D.W., Zumbo, B.D., 2015. Resolving the Issue of How Reliability is Related to Statistical Power: Adhering to Mathematical Definitions. Journal of Modern Applied Statistical Methods 14, 9–26. 10.56801/10.56801/v14.i.770

Zorowitz, S., Niv, Y., 2023. Improving the Reliability of Cognitive Task Measures: A Narrative Review. Biol Psychiatry Cogn Neurosci Neuroimaging 8, 789–797. 10.1016/j.bpsc.2023.02.004

Zuo, X.-N., Xu, T., Milham, M.P., 2019. Harnessing reliability for neuroscience research. Nat Hum Behav 3, 768–771. 10.1038/s41562-019-0655-x

