## Supplementary Information for "Reliability of structural brain change in cognitively healthy adult samples"

**Corresponding author:**

**Didac Vidal Piñeiro**

Department of Psychology, Pb. 1094 Blindern

Oslo, Norway, 0317

Tel: (+47) -22845061

#### Supplementary Methods

### Parameter selection

*Relationship between observed and true (latent) variability of the slopes*

We conducted a simulation to explore the relationship between *observed* and *true* variance. First, we simulated individual trajectories of brain change using **eq. 1.** The repeated measures for each individual (i) measured at a given set of equispaced time points ($j_1$, …, $j_n$) were derived from the slopes of change ($\beta_{2i}$) which follow a normal distribution in the population with a mean δ and variance $\sigma_s^2$. Similarly, (cross-sectional) measurement errors ($\varepsilon_{i,j}$) were assumed to be normally distributed with a mean of 0 and variance $\sigma_\varepsilon^2$. Intercepts ($\beta_{1i}$) are also assumed to be normally distributed with a non-zero mean and variance. The outcome variable $Y_{i,j}$ represents the measurement of a brain feature for an individual *i* at time *j*.

(eq. 1) $$Y_{i,j} = \beta_{1i} + \beta_{2i}t_j + \varepsilon_{i,j}$$

We generated two datasets. The first dataset included 250 individuals in each condition, defined by the combination of the number of observations and total study duration (observations = seq(3,9, 1); years (2, 12, 1). The second dataset replicated the number of observations, and the study duration of the subset used to estimate slope variance (**Supplementary Table 2**). The model parameters for each feature were those described in the main text, **methods section**. Next, for each individual and brain feature, we estimated *observed* change using linear models where time predicted brain measurements while the *true change* was provided by $\beta_{2i}$. We then calculated the dispersion of *true and observed* change for each number of observations × study duration condition in the first dataset, and for the entire population for the second dataset. This was repeated n = 100. The results were expressed as a percentage of observed-to-true variability.

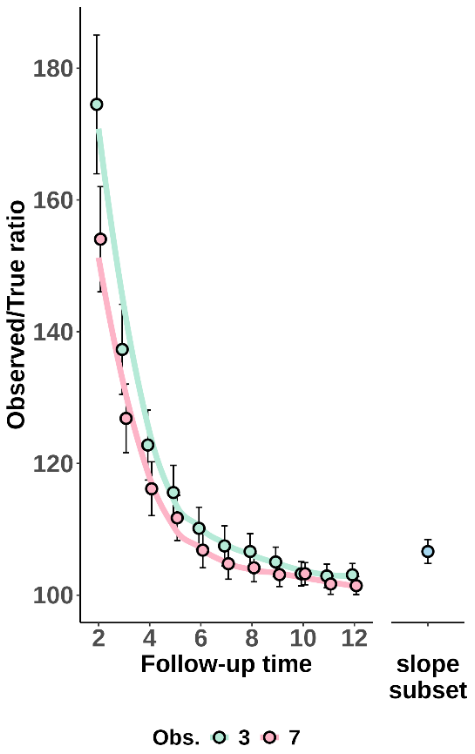


***Supplementary Figure 1. Ratio of observed to true variance of the slope.*** *Percentage overestimation of slope variance across all features, comparing observed variance to the true, latent slopes. The left panel displays the ratio as a function of follow-up duration and number of observations. The right panel presents the estimated ratio for the sample used to derive slope variance. On average, across all features, the overestimation was 10.4%, with slightly higher ratios observed in features with greater measurement error.*

*Slope variability to measurment error ratio* ($\sigma_s/\sigma_\varepsilon$).

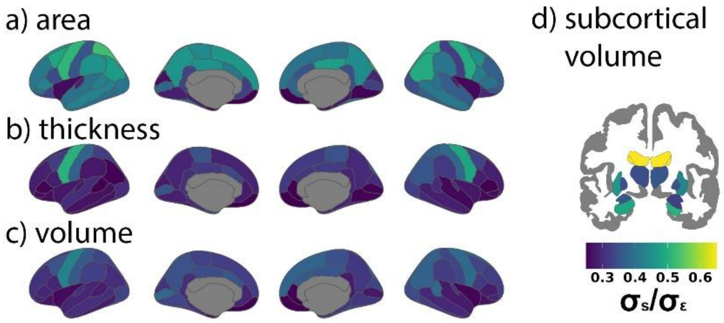


***Supplementary Figure 2. Ratio of slope dispersion to measurement error.*** *For each feature, the ratio of variability in brain change slopes* ($\sigma_s$) *- i.e. between-subject variability in brain change – to mean measurement error* ($\sigma_\varepsilon$)*. A greater ratio indicates higher longitudinal reliability for a given study duration and number of observations.*

#### Sample description (I). Longitudinal aging dataset.

A dataset composed of 11 different cohorts (n = 3611 unique individuals, n = 10964 observations) consisting of cognitively healthy adult participants with longitudinal MRI follow-ups was used for estimating slope variance and estimating *empirically* longitudinal reliability. The datasets include the *LCBC* (Walhovd et al., 2016), *Umeå* (Nyberg et al., 2010), and *UB* (Rajaram et al., 2017; Vidal-Piñeiro et al., 2014) datasets from the *Lifebrain* Consortium (Walhovd et al., 2018), the *COGNORM* (Idland et al., 2020), *ADNI* (Mueller et al., 2005), *AIBL* (Ellis et al., 2009), *HABS* (Dagley et al., 2017), *UKB* (https://www.ukbiobank.ac.uk/) (Miller et al., 2016), *preventAD* (Tremblay-Mercier et al., 2021), *OASIS3* (LaMontagne et al., 2019), and *Wayne* (Daugherty and Raz, 2016; Raz et al., 2012) datasets. See descriptives in **Supplementary Table 1** and visualization in **Supplementary Figure 3**. See a brief description of the datasets below. In addition to cohort-specific inclusion and exclusion criteria, individuals aged <18 years as indexed by mean age across observations and with severe neurological or psychiatric disorders were excluded. Further, observations with concurrent evidence of mild cognitive impairment, or Alzheimer's disease were excluded from the analyses while previous observations were kept. Observations from scanners with less than 15 measurements were also excluded for model fitting purposes. The initial dataset included individuals with 1 to 13 MRI acquisitions and longitudinal structural MRI scans were available for up to 15.8 years (mean = 4.1 [2.5] years). This dataset was fed into generalized additive mixed models (*gamm4 R-package*) (Wood, 2017) for harmonizing data across datasets and computing individual change over time. As harmonization generally benefits from a higher number of observations, individuals with either one or two observations were retained at this stage. Subsets of this dataset were used for estimating slope variance (**Supplementary Table 2**) and estimating longitudinal reliability *empirically* (**Supplementary Table 3**). All participants provided written informed consent and the studies were approved by the relevant ethical committees and conducted in accordance with the Declaration of Helsinki. See **Supplementary Table 5** for a summary on data availability, ethical standards, and contact information.

ADNI: The Alzheimer's Disease Neuroimaging Initiative (ADNI) (Mueller et al., 2005) is a multi-site project led by Doctor Michael W. Weiner to assess the progression of mild cognitive impairment (MCI) and early Alzheimer's Disease (AD), combining imaging, clinical and other biological markers, and neuropsychological and clinical assessments over time. For more information, visit https://adni.loni.usc.edu/about/. The age range for the participants is 55-90 years. The present study includes participants from ADNI 1, ADNIGO, ADNI2, and ADNI 3, who were cognitively healthy at baseline (*DX_bl* variable). Only observations in which participants were still cognitively healthy were included as determined by the ADNI team (*DX* variable). Amongst others, participants were required to have no evidence of ischemic stroke (Hachinski Ischemic Score ≤ 4), a Geriatric Depression scale score < 6, stable medications for 4 weeks before the screening, good auditory and visual acuity, good general health, no medical contraindications to MRI and at least 6 grades of education/work history. In-detailed general inclusion and exclusion criteria are described elsewhere (Petersen et al., 2010). All participants signed an informed consent form and the protocols were approved by the corresponding regional ethical committees in the US and Canada. Data was retrieved in April 2021.

<u>AIBL:</u> The Australian Imaging, Biomarker & Lifestyle Flagship Study of Ageing (AIBL) (Ellis et al., 2009) is a prospective study including cognitively normal participants, and patients with mild cognitive impairment, and AD aged 60 years or older. The study assesses the biomarkers, genetic factors, cognitive characteristics, and health and lifestyle factors that are associated with the development of AD combining techniques such as MRI, positron emission tomography (PET), blood tests, and fluid sample analysis, as well as neuropsychological and clinical assessments. Healthy participants must meet specific criteria, including being free of cognitive impairments and having test performance within 1.5 SD of age-adjusted norms. The test battery and sample description as well as in-detail general inclusion criteria have been described in detail previously (Ellis et al., 2009). All observations corresponding to cognitively normal individuals were included, as defined and evaluated by the AIBL team. All participants signed an informed consent form and the protocol was approved by the institutional human research committees of Austin Health, St Vincent's health, Hollywood Private Hospital, and Edit Cowan University (Australia). Data was retrieved in February 2023.

<u>COGNORM:</u> The COGNORM cohort (Idland et al., 2017) is an ongoing, prospective study coordinated by the Oslo University Hospital and Diakonhjemmet Hospital, Oslo, Norway. Patients (age ≥ 65 years) scheduled for elective gynecological, urological, or orthopedic surgery under spinal anesthesia were recruited. Participants were required to have no dementia, previous stroke with sequela, Parkinson's disease, or other neurodegenerative diseases that are likely to affect cognition. Patients with suspected undiagnosed dementia at any time within the first five years of follow-up (n = 15) (Sajjad et al., 2020), MMSE score <28 at baseline, and at least two abnormal cognitive test scores (-1.5 standard

deviation [SD] below the mean normal value for age, sex, and education) were excluded. All observations corresponding to cognitively normal individuals were included. All participants signed an informed consent form and the protocol was approved by the Norwegian Regional Committees for Medical and Health Research Ethics and the Data Protector Officer at Oslo University Hospital. Data was retrieved in March 2023.

HABS: The Harvard Aging Brain Study (HABS) (Dagley et al., 2017) is an ongoing, long-term observational study that aims to enhance our understanding of brain aging and the early stages of Alzheimer's disease. The study collects PET, MRI data, neuropsychological and clinical assessments. The age range was between 50 and 90 years at the time of baseline assessment and all patients were considered non-clinically impaired at the start of the study. Further participants had a CDR score of 0, MMSE score ≥ 25, < 11 on the Geriatric Depression Scale, and scores above age- and education-adjusted cutoffs on the 30-Minute Delayed Recall of the Logical Memory Story A to be included in the study. Participants with a history of alcoholism, drug abuse, head trauma, or current serious medical/psychiatric illness were excluded. Further details can be found elsewhere (Dagley et al., 2017). Observations with MCI or AD diagnostic (DX variable) were excluded. All participants signed an informed consent form and the protocol was approved by the Partners Healthcare Human Research Committee. Data was retrieved in August 2022.

LCBC: The Center for Lifespan Changes in Brain and Cognition cohort (LCBC, Oslo) (Fjell et al., 2023; Walhovd et al., 2016) consists of cognitively healthy, community-dwelling participants across the lifespan and is drawn from studies coordinated by the LCBC Research Group (LCBC www.oslobrains.no), approved by a Norwegian Regional Committee for Medical and Health Research Ethics. Written informed consent was obtained from all participants. The samples were recruited by a variety of methods such as newspapers and webpage ads. Most participants were recruited for observational studies, some currently ongoing, while a minority were recruited to enter into cognitive training. Written informed consent was obtained from all adult participants. All participants had to undergo a standardized health interview before being included in the study, and those with a history of neurological or psychiatric conditions or who reported concerns about their cognitive function were excluded. Additionally, all participants over the age of 40 years were required to score at least 25 on the Mini-Mental State Examination. The LCBC cohort was part of the Lifebrain obtained as part of the

Lifebrain consortium (Walhovd et al., 2018). MRI observations paired with MMSE ≤ 25 were excluded. Data was retrieved in November 2022.

OASIS3: The Open Access Series of Imaging Studies (OASIS3) (LaMontagne et al., 2019) is a retrospective collection of multimodal data that focuses on aging and AD and is openly accessible to the scientific community. OASIS-3 includes neuroimaging, clinical and neuropsychological data. Participants were recruited through the Washington University Knight Alzheimer Disease Research Center via flyers, word of mouth, and community engagements and were aged between 42 and 95 years. Only participants deemed cognitively normal at baseline were included in the observations. Exclusion criteria included medical conditions that precluded longitudinal participation or medical contraindications for the different study arms. See in-detail inclusion and exclusion criteria (LaMontagne et al., 2019). All participants consented to Knight ADRC-related projects following procedures approved by the Institutional Review Board of Washington University School of Medicine. Observations were included until the last observation in which a subject was deemed cognitively healthy as determined by the Clinical Dementia Rating Scale (CDR) (Morris, 1993).

PreventAD: The Pre-symptomatic Evaluation of Experimental or Novel Treatments for AD (PREVENT-AD) (Tremblay-Mercier et al., 2021) is a retrospective, long-term study that follows cognitively healthy older individuals with a familiar history of AD. It includes participants enrolled either from an observational cohort or the clinical trial of PREVENT-AD. This study comprises MRI images, blood and CSF samples, and clinical and neuropsychological assessments. Participants in the study had to be at least 60 years old, had ≥6 years of education, and they needed to be cognitively unimpaired at baseline. The Montreal Cognitive Assessment (MoCA) and CDR scales were used to assess cognitive abilities, and participants were considered cognitively intact if their MoCA scores were ≥ 26/30 or their CDR was = 0. Other exclusion criteria at baseline included medical conditions that prevented longitudinal participation or medical contraindications to MRI, use of acetylcholinesterase inhibitors, other approved prescription cognitive enhancers, hypertension, or substance abuse. The inclusion and exclusion criteria have been previously described in detail (Tremblay-Mercier et al., 2021). The protocols, consent forms, and study procedures were approved by the McGill Institutional Review Board and the Douglas Mental Health University Institute Research Ethics Board. Observations with RBANS > 1SD below the mean and probable MCI, as evaluated by a clinician, were excluded.

UB: The University of Barcelona cohort consisted of a series of retrospective substudies (Abellaneda-Pérez et al., 2019; Rajaram et al., 2017; Uribe et al., 2016; Vidal-Piñeiro et al., 2014). In all cases, samples consisted of cognitively healthy, community-dwelling participants with normal cognitive and visual function. Most participants were recruited for observational studies while a minority were recruited to enter into cognitive training. Inclusion criteria varied a bit across studies (see specific studies for details), but include severe neurologic and psychiatric disorders, recent head trauma or brain surgery, cognitive deterioration, or dementia with a score < 24 on the Mini-Mental State Examination and additional neuropsychological criteria, other neurodegenerative disorders like Parkinson's disease and chronic illness with a projected shortened lifespan. Observations with MMSE < 26 were excluded. All participants signed an informed consent and the protocols were approved by the ethical committees of the University of Barcelona and of the Hospital Clinic of Barcelona. The LCBC cohort was part of the Lifebrain obtained as part of the Lifebrain consortium (Walhovd et al., 2018).

UKB: The UK Biobank (UKB) (https://www.ukbiobank.ac.uk/about-biobank-uk/) is a major national and international health resource with the aim of improving the prevention, diagnosis and treatment of a wide range of illnesses. UK Biobank recruited ≈500,000 people aged between 40-69 years in 2006-2010 from across the country to take part in this project (Guggenheim et al., 2015). Potential participants were identified through National Health Service (NHS) registers according to being aged 40-69 and living within a reasonable traveling distance of an assessment center. Assessment centers are located in accessible and convenient locations with a large surrounding population. Participants have undergone measures and provided samples and detailed information about themselves and agreed to have their health followed. The study sample was drawn from the UK Biobank neuroimaging branch (Miller et al., 2016) and conducted under data application number 32048. Only individuals with longitudinal MRI data were used in this study. Participants signed an informed consent and the protocols were approved by the North West Multi-Center Research Ethics Committee [MREC]; see also https://www.ukbiobank.ac.uk/the-ethics-and-governance-council.

Umeå: The BETULA project (Umeå; Nilsson et al., 2010) is a prospective longitudinal study on aging, memory, and dementia, which used a population-based sampling of healthy middle-aged and older adults for recruitment. Detailed recruitment procedures are found elsewhere (Nilsson et al., 2004, 1997). For the current analyses, the MRI subsample of the study is used. Participation in the neuroimaging study was offered to all participants who had remained in the study and completed cognitive testing at the 5th Betula test wave onwards. Exclusion criteria were severe visual or auditory handicaps, intellectual or developmental disabilities, suspected dementia, having a mother tongue other than Swedish, MRI contraindications, severe neurological disorders, or visual/motor deficits that could interfere with fMRI data collection, MMSE <24, brain or head surgery, and substantial brain anatomical deviations. Some participants were later excluded due to discovered neurological conditions, severe depression, and MRI anatomical abnormalities. The LCBC cohort was part of the Lifebrain obtained as part of the Lifebrain consortium (Walhovd et al., 2018). All participants signed an informed consent and the protocols were approved by the Regional Ethical Vetting Board at Umeå University. Data was retrieved in August 2022.

Wayne: The Wayne cohort consists of two retrospective longitudinal datasets which are part of the Brain Aging in Detroit Longitudinal Study. The overarching aim of the study is to understand the mechanisms driving human brain changes over the adult lifespan, identify the risk factors and protective influences that modify the rate of change, and elucidate the relationships between changes in brain properties and cognitive performance. See a detailed description of the sample and inclusion criteria elsewhere (Burgmans et al., 2010; Kennedy and Raz, 2009). Briefly, all participants were screened at baseline via a health questionnaire for the following conditions: the presence of cardiovascular, neurological, or psychiatric disease, use of centrally acting medications, the habit of having three or more alcoholic drinks per day, as well as being minimally high school educated, native English speakers were inclusion/exclusion criteria. Healthy volunteers from the metropolitan Detroit area who responded to media advertisements and flyers participated in this study. Further exclusion criteria included MMSE < 26 and GDS >16. All participants provided written informed consent. The study protocols were approved by the (Wayne State) University Institutional Review Board.

##### *Sample description (I.I). Slope variance subset*

$\mathrm{Var}(\delta)$ was estimated using a subset of the individuals described in the *Longitudinal aging dataset.* Specifically, we selected individuals with follow-up > 4 years and with 4 or more observations from 8 of these datasets. See **Supplementary Figure 4** and **Supplementary Table 2** for the sample's descriptive statistics and visualization.

##### *Sample description (I.II). “Empirical” subset*

Longitudinal reliability was estimated directly from data. Specifically, we used all individuals described in the *Longitudinal aging dataset* with 3 or more observations. See **Supplementary Figure 5** and **Supplementary Table 3** for the sample’s descriptive statistics and visualization.

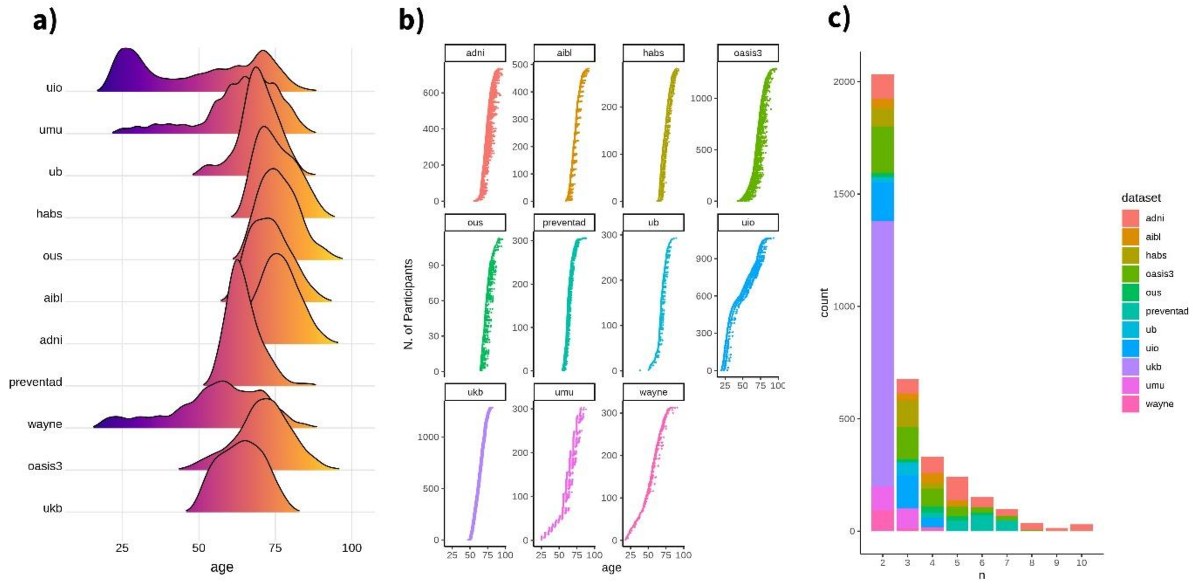


***Supplementary Figure 3. Sample descriptives for the longitudinal aging dataset.*** *a) Ridgeline plot showing the distribution of observations across age for each dataset. b) Spaghetti plots showing individual follow-ups across age for each dataset. c) Histogram showing the distribution of observations per participant, grouped also by dataset. Ten or more observations are collapsed. Note that this dataset was used only for harmonization purposes. Subsamples of this dataset are used for estimating reliability "empirically" and estimating the slope dispersion parameter. See* ***Supplementary Table 1*** *for associate descriptives.*

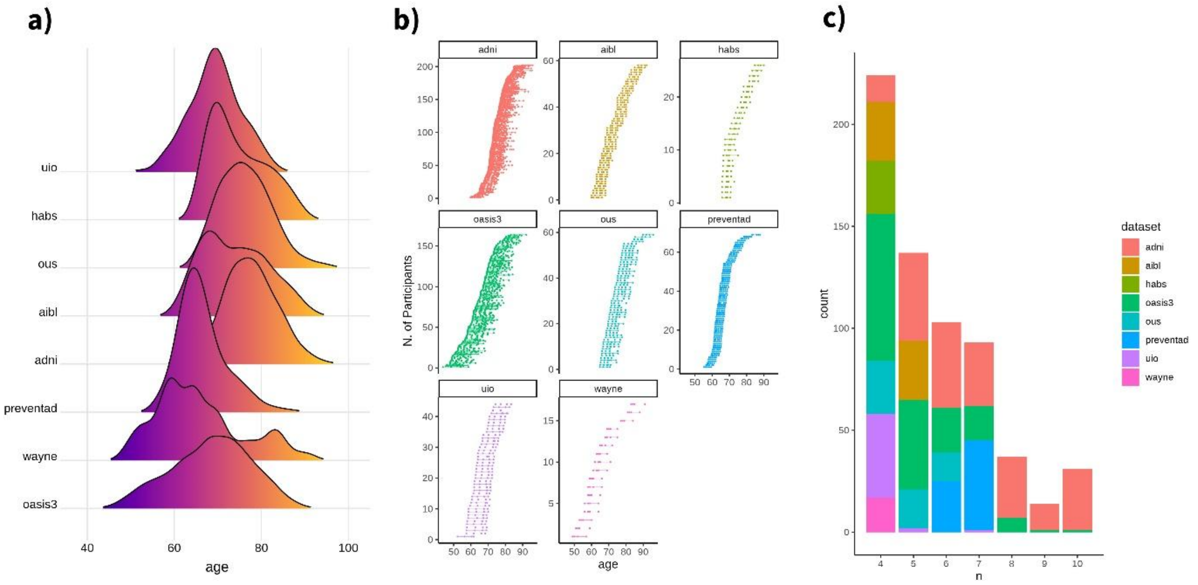


***Supplementary Figure 4. Sample descriptives for the slope dispersion dataset.*** *a) Ridgeline plot showing the distribution of observations across age for each dataset. b) Spaghetti plots showing individual follow-ups across age for each dataset. c) Histogram showing the distribution of observations per participant, grouped also by dataset. Ten or more observations are collapsed. This dataset is a subset of the longitudinal aging dataset (see **Supplementary Figure 3**, **Supplementary Table 1**) and was used to estimate the slope dispersion parameter. Only individuals with 4 or more observations and follow-up of at least 4 years were selected. See **Supplementary Table 2** for associate descriptives.*

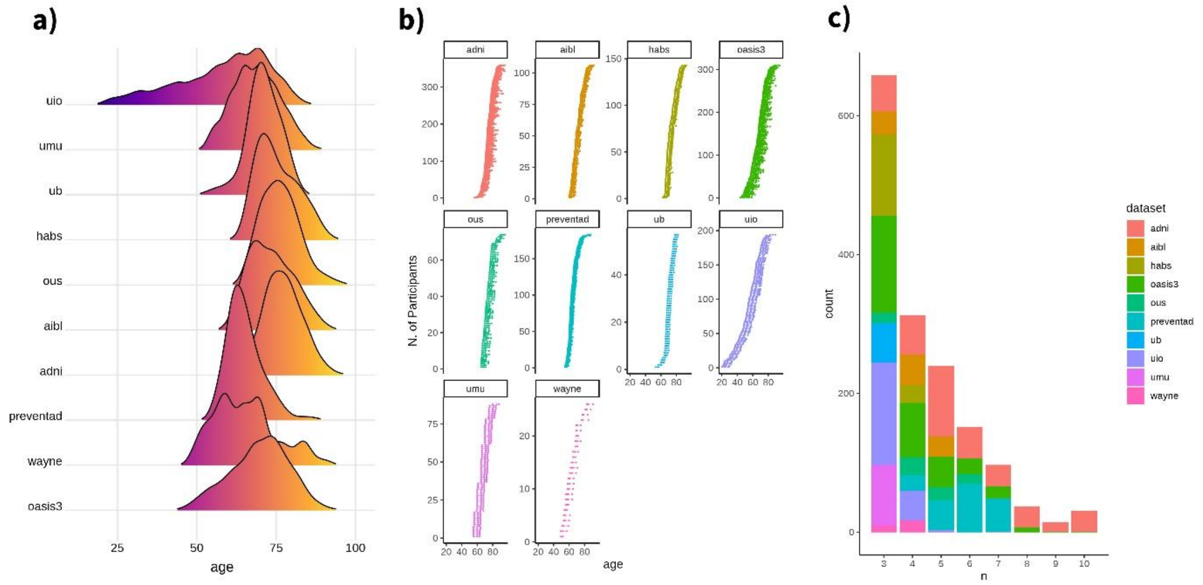


***Supplementary Figure 5. Sample descriptives for the "empirical" dataset.*** *a) Ridgeline plot showing the distribution of observations across age for each dataset. b) Spaghetti plots showing individual follow-ups across age for each dataset. c) Histogram showing the distribution of observations per participant, grouped also by dataset. Ten or more observations are collapsed. This dataset is a subset of the longitudinal aging dataset (see **Supplementary Figure 3**, **Supplementary Table 1**) and was used to estimate the reliability of brain change "empirically". Only individuals with 3 or more observations were selected. See **Supplementary Table 3** for associate descriptives.*

| | N | F:M | Obs. | Age | | | | Follow-up time | | | | N. Observations | | | | Scanners |
|---|---|---|---|---|---|---|---|---|---|---|---|---|---|---|---|---|
| | | | | mean | SD | Min | max | mean | SD | min | max | mean | SD | min | max | |
| *adni* | 494 | 270:224 | 2371 | 73.23 | 5.93 | 55.80 | 89.90 | 4.41 | 2.64 | 1.60 | 15.00 | 4.80 | 2.44 | 2 | 13 | 88 |
| *aibl* | 152 | 79:73 | 513 | 71.09 | 6.33 | 60.00 | 87.00 | 4.09 | 1.64 | 2.00 | 8.00 | 3.38 | 1.10 | 2 | 5 | 2 |
| *habs* | 223 | 129:94 | 615 | 73.34 | 6.08 | 62.50 | 89.25 | 4.41 | 1.17 | 2.00 | 8.50 | 2.76 | 0.65 | 2 | 4 | 1 |
| *oasis3* | 518 | 295:223 | 1697 | 67.84 | 9.28 | 43.52 | 93.01 | 6.12 | 3.35 | 1.50 | 15.84 | 3.28 | 1.49 | 2 | 10 | 5 |
| *ous* | 94 | 51:43 | 368 | 73.31 | 6.18 | 64.74 | 89.79 | 5.98 | 2.60 | 1.83 | 9.51 | 3.91 | 1.35 | 2 | 6 | 1 |
| *pre-ventad* | 187 | 133:54 | 1075 | 63.68 | 5.21 | 55.13 | 84.23 | 3.27 | 0.84 | 1.56 | 4.69 | 5.75 | 1.00 | 3 | 7 | 1 |
| *ub* | 79 | 50:29 | 215 | 68.44 | 5.02 | 51.70 | 78.10 | 3.58 | 1.02 | 1.70 | 5.00 | 2.72 | 0.45 | 2 | 3 | 1 |
| *uio* | 366 | 218:148 | 979 | 50.01 | 18.99 | 19.62 | 85.47 | 5.90 | 2.63 | 1.99 | 11.53 | 2.67 | 0.75 | 2 | 7 | 3 |
| *ukb* | 1179 | 593:586 | 2358 | 62.68 | 7.16 | 47.02 | 79.52 | 2.25 | 0.12 | 2.01 | 2.67 | 2.00 | 0.00 | 2 | 2 | 2 |
| *umu* | 196 | 88:108 | 480 | 59.52 | 12.46 | 25.00 | 80.00 | 6.06 | 1.83 | 3.00 | 9.00 | 2.45 | 0.50 | 2 | 3 | 1 |
| *wayne* | 123 | 78:45 | 293 | 56.86 | 12.97 | 19.00 | 83.00 | 3.21 | 2.16 | 2.00 | 9.00 | 2.38 | 0.72 | 2 | 4 | 2 |
| *all* | 3611 | 1984:1627 | 10964 | 64.67 | 11.75 | 19.00 | 93.01 | 4.10 | 2.54 | 1.50 | 15.84 | 3.04 | 1.62 | 2 | 13 | 107 |

***Supplementary Table 1. Descriptives of the longitudinal aging dataset.*** *Main sociodemographic and study descriptives of the longitudinal aging dataset. See* ***Supplementary Figure 3*** *for a visual representation. F:M = Female:Males. Obs. = Total number of observations. N. Observations = Observations per participant.*

| | *N* | *F:M* | *obs* | ***Age*** | | | | ***Follow-up time*** | | | | ***N. Observations*** | | | | *Scanners* |
|---|---|---|---|---|---|---|---|---|---|---|---|---|---|---|---|---|
| | | | | **mean** | **SD** | **Min** | **max** | **mean** | **SD** | **min** | **max** | **mean** | **SD** | **min** | **max** | |
| *adni* | 202 | 96:106 | 1415 | 73.80 | 5.55 | 59.80 | 89.70 | 7.01 | 2.20 | 4.10 | 15.00 | 7.00 | 2.06 | 4 | 13 | 66 |
| *aibl* | 58 | 35:23 | 261 | 70.81 | 7.18 | 60.00 | 87.00 | 5.76 | 0.68 | 5.00 | 8.00 | 4.50 | 0.50 | 4 | 5 | 2 |
| *habs* | 26 | 10:16 | 104 | 71.95 | 6.32 | 65.75 | 84.75 | 5.34 | 0.82 | 4.75 | 8.50 | 4.00 | 0.00 | 4 | 4 | 1 |
| *oasis3* | 164 | 105:59 | 834 | 63.65 | 8.80 | 43.80 | 83.19 | 9.60 | 2.77 | 4.01 | 15.84 | 5.09 | 1.27 | 4 | 10 | 5 |
| *ous* | 59 | 31:28 | 283 | 72.67 | 5.52 | 64.82 | 89.79 | 7.49 | 1.75 | 4.08 | 9.51 | 4.80 | 0.80 | 4 | 6 | 1 |
| *pre-ventad* | 69 | 49:20 | 458 | 64.90 | 5.34 | 55.30 | 83.25 | 4.17 | 0.14 | 4.01 | 4.69 | 6.64 | 0.48 | 6 | 7 | 1 |
| *uio* | 44 | 22:22 | 181 | 72.74 | 4.55 | 60.30 | 83.39 | 9.61 | 1.10 | 4.67 | 11.53 | 4.11 | 0.49 | 4 | 7 | 2 |
| *wayne* | 17 | 12:9 | 68 | 62.88 | 9.98 | 49.00 | 83.00 | 7.41 | 0.80 | 6.00 | 9.00 | 4.00 | 0.00 | 4 | 4 | 1 |
| *all* | 639 | 356:283 | 3604 | 69.42 | 8.09 | 43.80 | 89.79 | 7.42 | 2.67 | 4.01 | 15.84 | 5.64 | 1.78 | 4 | 13 | 79 |

***Supplementary Table 2. Descriptives of the slope dispersion dataset.*** *Main sociodemographic and study descriptives of the longitudinal aging dataset. See* ***Supplementary Figure 4*** *for a visual representation. F:M = Female:Males. Obs. = Total number of observations. N. Observations = Observations per participant.*

| | *N* | *F:M* | *obs* | ***Age*** | | | | ***Follow-up time*** | | | | ***N. Observations*** | | | | *Scanners* |
|---|---|---|---|---|---|---|---|---|---|---|---|---|---|---|---|---|
| | | | | **mean** | **SD** | **Min** | **max** | **mean** | **SD** | **min** | **max** | **mean** | **SD** | **min** | **max** | |
| *adni* | 387 | 196:191 | 2157 | 73.84 | 5.89 | 56.30 | 89.70 | 5.01 | 2.69 | 1.60 | 15.00 | 5.57 | 2.19 | 3 | 13 | 86 |
| *aibl* | 107 | 52:55 | 423 | 70.82 | 6.58 | 60.00 | 87.00 | 4.84 | 1.24 | 2.00 | 8.00 | 3.95 | 0.77 | 3 | 5 | 2 |
| *habs* | 143 | 86:57 | 455 | 72.63 | 6.08 | 62.50 | 87.75 | 5.12 | 0.65 | 2.75 | 8.50 | 3.18 | 0.39 | 3 | 4 | 1 |
| *oasis3* | 312 | 187:125 | 1285 | 66.27 | 9.17 | 43.80 | 89.30 | 7.79 | 3.14 | 1.55 | 15.84 | 4.12 | 1.38 | 3 | 10 | 5 |
| *ous* | 74 | 42:32 | 328 | 72.79 | 5.58 | 64.82 | 89.79 | 6.91 | 1.97 | 3.90 | 9.51 | 4.43 | 1.02 | 3 | 6 | 1 |
| *pre-ventad* | 187 | 133:54 | 1075 | 63.68 | 5.21 | 55.13 | 84.23 | 3.27 | 0.84 | 1.56 | 4.69 | 5.75 | 1.00 | 3 | 7 | 1 |
| *ub* | 57 | 37:20 | 171 | 68.95 | 4.96 | 53.10 | 78.10 | 4.18 | 0.38 | 3.30 | 5.00 | 3.00 | 0.00 | 3 | 3 | 1 |
| *uio* | 194 | 116:78 | 635 | 55.66 | 16.01 | 20.05 | 83.39 | 7.82 | 1.96 | 2.47 | 11.53 | 3.27 | 0.54 | 3 | 7 | 3 |
| *umu* | 88 | 39:49 | 264 | 64.46 | 6.40 | 55.00 | 80.00 | 7.95 | 0.55 | 7.00 | 9.00 | 3.00 | 0.00 | 3 | 3 | 1 |
| *wayne* | 30 | 16:14 | 107 | 62.53 | 9.36 | 49.00 | 83.00 | 5.83 | 2.45 | 2.00 | 9.00 | 3.57 | 0.50 | 3 | 4 | 1 |
| *all* | 1579 | 904:675 | 6900 | 67.63 | 10.33 | 20.05 | 89.79 | 5.94 | 2.72 | 1.55 | 15.84 | 4.37 | 1.70 | 3 | 13 | 102 |

***Supplementary Table 3. Descriptives of the empirical dataset.*** *Main sociodemographic and study descriptives of the "empirical" dataset. See **Supplementary Figure 5** for a visual representation. F:M = Female:Males. Obs. = Total number of observations. N. Observations = Observations per participant.*

#### Sample description (II). Test-retest dataset

The test-retest dataset was composed of 6 different test-retest cohorts consisting of cognitively healthy adult participants. This dataset was used to estimate cross-sectional measurement error $\sigma_e$ for the different cortical and subcortical features. This dataset was composed of the following studies: the S2C (Walhovd et al., 2024), the preventAD (Orban et al., 2015), OASIS (Marcus et al., 2007), and GSP (Holmes et al., 2015) reliability subsets, and the HNU1 (Chen et al., 2015) and Maclaren (Maclaren et al., 2014). See a brief description of the datasets below.

GSP reliability subset: This cohort is a test-retest subset of the Brain Genomics Superstruct Project (GSP) (Holmes et al., 2015). The goal of the GSP is to enable large-scale exploration of the links between brain function, behavior, and genetic variation. The authors made openly available a repository of structural and functional MRI, coupled with genetic information, and self-report behavioral and cognitive measures from a sample of cognitively healthy individuals (aged between 18 and 35 years). Most participants were aware of the study through local college recruitment efforts and studies connected to Harvard University and the Massachusetts General Hospital. Participants were only enrolled if they were participating in a study of normal brain function or serving as control participants in a case-control study of a clinical population. Participants were excluded from the present data release if their self-reported health information indicated current or past history of Axis I pathology or neurological disorder, current psychotropic medication usage, acute physical illness, or displayed atypical brain anatomy. The test-retest reliability datasets are included from 69 participants scanned within six months of their initial visit (77.2 [55.9] days). Data was made publicly available as part of the Consortium for Reliability and Reproducibility (CoRR) (Zuo et al., 2014). Participants provided written informed consent in accordance with guidelines established by the Partners Health Care Institutional Review Board and the Harvard University Committee on the Use of Human Subjects in Research.

HNU1: The Hangzhou Normal University (HNU1) test-retest dataset (Chen et al., 2015) is a set of scanners made openly available to study the functional and structural variability of the human brain. The dataset consists of 30 healthy adults (aged between 20 and 30 years) who were scanned within one month, every 2-3 days, resulting in ten repeated measurements for each participant. Both resting-state and structural MRI sequences were acquired. The test-retest has been made openly available (http://dx.doi.org/10.15387/fcp_indi.corr.hnu1) as part of the Consortium for Reliability and Reproducibility (CoRR) (Zuo et al., 2014), which shares the data via the Neuroimaging Informatics Tools and

Resources Clearinghouse (NITRC). None of the participants had a history of neurological or psychiatric disorders, substance abuse, or head injury with loss of consciousness. The ethics committee of the Center for Cognition and Brain Disorders (CCBD) at Hangzhou Normal University approved this study and written informed consent was obtained from all participants.

Maclaren: The Maclaren test-retest dataset (Maclaren et al., 2014) is an openly available dataset aiming to generate publicly available data to assist in the validation and reliability of current and future segmentation methods. The dataset consists of 120 structural MRIs from 3 cognitively healthy adult individuals (40 scanners each) which were acquired in 20 different sessions for 31 days. Each subject was scanned twice within each session, with repositioning between the two scans. Human data collection was performed with the approval of the Stanford University Institutional Review Board and all subjects gave their written informed consent. The data was obtained from the OpenNeuro database (Markiewicz et al., 2021) with accession number ds000239.

OASIS reliability subset: This cohort is a test-retest subset of the Open Access Series of Imaging Studies (OASIS) (Marcus et al., 2007). OASIS consists of a series of MRI datasets that are publicly available for study and analysis. The initial dataset, from which the reliability subset was derived, consists of a cross-sectional collection of individuals aged 18 to 96 years who were cognitively healthy or clinically diagnosed with MCI or AD. OASIS is an open science project aiming to freely share MRI datasets with the scientific community and facilitate future discoveries in basic and clinical neuroscience. See above (OASIS-3 sample description in the *Longitudinal Aging dataset s*ection) and elsewhere for detailed inclusion criteria (Marcus et al., 2007). The test-retest reliability subset consists of 20 cognitively healthy individuals (aged between 19 and 29 years) who underwent a second MRI session within 90 days of their initial visit (mean delay = 20.55 days). Protocols were approved by the Washington University Human Studies Committee and individuals gave informed consent. The cohort was coordinated by Washington University, but individuals included in the reliability subset are not part of the *OASIS* dataset described in the *Longitudinal Aging dataset* section.

PreventAD reliability subset: This cohort is a test-retest resting-state and structural MRI subset consisting of 80 cognitively normal elderly volunteers (65.4 [6.2] years) derived from the "Pre-symptomatic Evaluation of Novel or Experimental Treatments for Alzheimer's Disease" (PREVENT-AD) Cohort (Orban et al., 2015). The cohort consisted of individuals with a family history of Alzheimer's disease in first-degree relatives who were enrolled in a double-blind randomized clinical trial (Tremblay-Mercier et al., 2021). Two scanning sessions were acquired ≈3 months apart. Data was made publicly available as the sample *UM1* of the Consortium for Reliability and Reproducibility (CoRR) (Zuo et al., 2014). See above (*PreventAD* sample description in the *Longitudinal Aging dataset* section) and elsewhere (Orban et al., 2015) for detailed inclusion criteria. The cohort was assembled at the StoP-AD Centre, Montreal, Canada. All subjects had given informed consent and the study was approved by the Research, Ethics, and Compliance Committee of McGill University. Individuals included in the reliability subset are part of the preventAD dataset described in the *Longitudinal Aging dataset* section.

S2C reliability subset: The Set-2-Change cohort is a Norwegian twin cohort coordinated by the Research Group for Lifespan Changes in Brain and Cognition (LCBC www.oslobrains.no). The S2C sample was recruited through the Norwegian Mother, Father, and Child Cohort Study (MoBa) (Magnus et al., 2016), and the Norwegian Twin Registry (NTR) (Nilsen et al., 2019), that is, this study includes data from MoBa and NTR, and both studies are conducted by the Norwegian Institute of Public Health. The Set-2-Change project consists of a cognitive training design based on spatial navigation learning, virtual reality, and exercise. All participants had to undergo a standardized health interview before being included in the study, and those with a history of neurological or psychiatric conditions were excluded. The reliability subset consists of 139 participants (between 18.2 and 82.8 years) scanned twice, approximately 3 months apart. The participants signed an informed consent and the study was approved by a Norwegian Regional Committee for Medical and Health Research Ethics. The cohort was coordinated by LCBC, but individuals included in the reliability subset are not part of the *LCBC* dataset described in the *Longitudinal Aging dataset* section.

| | N | F:M | Obs. | Age | | | | Delay | | | |
|---|---|---|---|---|---|---|---|---|---|---|---|
| | | | | mean | SD | min | max | mean | SD | min | max |
| *gsp* | 69 | 34 | 2 | 21.12 | 2.59 | 19.00 | 29.00 | 77.19 | 56.13 | 2 | 175 |
| *hnu* | 30 | 15 | 10 | 24.37 | 2.41 | 20.00 | 30.00 | 33.07 | 2.88 | 31 | 40 |
| *maclaren* | 3 | 2 | 40 | 29.00 | 2.65 | 26.00 | 31.00 | 31.00 | 0.00 | 31 | 31 |
| *oasis* | 20 | 0 | 2 | 23.40 | 4.03 | 19.00 | 34.00 | 20.55 | 23.86 | 1 | 89 |
| *preventad* | 80 | 58 | 2 | 65.36 | 6.26 | 55.00 | 84.00 | 111.40 | 24.26 | 74 | 194 |
| *s2c* | 139 | 42 | 2 | 35.22 | 15.42 | 18.17 | 78.55 | 82.75 | 24.41 | 49 | 288 |

***Supplementary Table 4. Descriptives of the test-retest dataset.*** *Main sociodemographic and study descriptives of the test-retet, measurement error dataset. F:M = Female:Males. Obs. = Number of observations per participant. Delay = Time between the first and the last observations in days.*

| *Sample* | *Link* | *PI and/or Admin Contact* | *IRB* |
|---|---|---|---|
| | | *Longitudinal Aging Dataset* | |
| *ADNI* | https://adni.loni.usc.edu/ (O) | Weiner MW; (PI) (AC) | Approved by the Institutional Review Boards of all of the participating institutions |
| *AIBL* | https://aibl.csiro.au/research/ (O) | Christopher Rowe; (PI) | Institutional ethics committees of Austin Health, StVincent's Health, Hollywood |
| *COGNORM* | https://www.med.uio.no/klinmed/english/research/groups/delirium/index.html; http://www.oslobrains.no (R) | Leiv Otto Watne; (PI); Anders Martin Fjell; (PI) | Norwegian Regional Committees for Medical and Health Research Ethics and the Data Protector Officer at Oslo University Hospital |
| *HABS* | https://habs.mgh.harvard.edu | Reisa Sperling; (PI); (AC) | Partners Healthcare Human Research Committee |
| *LCBC* | http://www.oslobrains.no (R) | Kristine B. Walhovd; (PI) | Norwegian Regional Committee for Medical and Health Research Ethic; Regional Ethical Committee of South Norway |
| *OASIS3* | https://www.oasis-brains.org/ (O) | Pamela J. LaMontagne; (PI); Daniel Marcus; (PI); https://www.oasis-brains.org/#contact (AC) | Institutional Review Board of Washington University School of Medicine |
| *preventAD* | https://prevent-alzheimer.net; https://openpreventad.loris.ca/ (O) | Jennifer Tremblay; (PI); https://openpreventad.loris.ca/contact/ (AC) | The McGill Institutional Review Board and the Douglas Mental Health University Institute Research Ethics Board |
| *UB* | http://www.ub.edu/bbslab/bbslab/ (R) | David Bartrés-Faz; (PI) | Comisión de Bioética de la Universidad de Barcelona and Hospital Clinic |
| *UKB* | https://www.ukbiobank.ac.uk/ (O) | Rory Collins (PI); (AC) | North West Multi-Center Research Ethics Committee [MREC]; https://www.ukbiobank.ac.uk/the-ethics-and-governance-council |
| *Umeå* | http://www.ufbi.umu.se/english (R) | Lars Nyberg; (PI) | Regional Ethical Vetting Board at Umeå University |
| *Wayne* | http://fcon_1000.projects.nitrc.org/indi/retro/wayne_10. | Naftali Raz (PI) | The (Wayne State) University Institutional Review Board |

| | | | |
|---|---|---|---|
| | html; http://fcon_1000.projects.nitrc.org/indi/retro/wayne_11.html | | |
| | Test-retest (reliability) dataset | | |
| *GSP* | https://www.neuroinfo.org/gsp/ (O) | Randy L. Buckner; (PI); (AC) | Partners Health Care Institutional Review Board and the Harvard University Committee on the Use of Human Subjects in Research |
| *HNU1* | http://fcon_1000.projects.nitrc.org/indi/CoRR/html/hnu_1.html (O) | Xi-Nian Zuo;.c (PI); Michael P. Milham; (PI) | The ethics committee of the Center for Cognition and Brain Disorders at Hangzhou Normal Uni-versit |
| *Maclaren* | https://openneuro.org/datasets/ds000239/versions/00001 (O) | Julian Maclaren; (PI) | Stanford University Institutional Review Board |
| *preventAD* | https://fcon_1000.projects.nitrc.org/indi/CoRR/html/um_1.html (O) | Pierre Orban; (PI); Pierre Bellec; (PI) | The Research, Ethics, and Compliance Committee of McGill University |
| *OASIS* | https://www.oasis-brains.org/ (O) | Randy L. Buckner; (PI); Daniel Marcus; (PI); https://www.oasis-brains.org/#contact (AC) | Washington University Human Studies Committee |
| *S2C* | http://www.oslobrains.no (R) | Kristine B. Walhovd; (PI) | Norwegian Regional Committee for Medical and Health Research Ethic; Regional Ethical Committee of South Norway |

***Supplementary Table 5. Data availability.*** *Data availability, contact and principal investigator information, and ethical approval for the different datasets used. PI = Principal Investigator. AC = Administrative contact. IRB = Institutional Review Boards. O = Openly available. Automatic or semi-automatic data agreements. Fees may apply (e.g. UKB). R = Restricted. Ad-hoc permission is required. Contact PI or AC for specific details on securing access to data.*

#### MRI acquisition and processing

Structural T1-weighted (T1w) MPRAGE, FSPGR, and IR-SPGR scans were collected using 1.5, 3, and 4 T scanners. See information on scanner parameters and scanners per dataset in **Supplementary Table 6**. When not obtained in BIDS form, data was transformed into the Brain Imaging Data Structure (BIDS) format (Gorgolewski et al., 2016). ADNI, AIBL, and HABS transformation to BIDS were performed with Clinica software (Routier et al., 2021; Samper-González et al., 2018). We used the longitudinal FreeSurfer v.7.1.0 stream (Reuter et al., 2012) for cortical reconstruction of the structural T1w scans (Dale et al., 1999; Fischl et al., 1999). We averaged scanners when multiple scanners were available from the same session. Briefly, the images were processed using the cross-sectional stream, which includes the removal of nonbrain tissues, Talairach transformation, intensity correction, tissue and volumetric segmentation, cortical surface reconstruction, and cortical parcellation. Next, an unbiased within-subject template space based on all cross-sectional images was created for each participant, using robust, inverse-consistent registration (Reuter et al., 2010). The processing of each time point was then reinitialized with common information from the within-subject template, to increase reliability and statistical power. Data were summarized based on the *Desikan* atlas (Desikan et al., 2006) for cortical thickness and cortical area measures ($|N|$ = 34 features per hemisphere and modality) and the *aseg* atlas for subcortical volumetric data (Fischl et al., 2002) from which we selected the left and right Lateral Ventricle, Thalamus, Caudate, Putamen, Pallidum, Hippocampus, and Amygdala volumes. Except for the Umeå dataset, all data was preprocessed on the Colossus processing cluster, part of the Tjenester for Sensitive Data (TSD), University of Oslo. For the cross-sectional data, values were summarized from the output of the initial cross-sectional stream. The same datasets and individuals were used except for the Umeå dataset for which we only had longitudinal data available.

| *Sample* | *Scanner* | *Field* | *Sequence parameters* |
|---|---|---|---|
| | | | *Longitudinal Aging Dataset* |
| *ADNI* | Multisite (n > 50) | 1.5/ 3.0 | See https://adni.loni.usc.edu/methods/documents/mri-protocols/ |
| *AIBL* | Avanto Siemens | 1.5 | MPRAGE. TR: 2300 ms; TE: 2.98 ms, TI: 900 ms; flip angle 9°, slice thickness: 1.25 mm, FoV: 240 x 256, 160 slices. |
| | Verio Siemens | 3.0 | MPRAGE. TR: 2300 ms; TE: 2.98 ms, TI: 900 ms; flip angle 9°, slice thickness: 1.25 mm, FoV: 240 x 256, 160 slices. |
| | Tim Trio Siemens | 3.0 | MPRAGE. TR: 2300 ms; TE: 2.98 ms, TI: 900 ms; flip angle 9°, slice thickness: 1.25 mm, FoV 240 x 256, 160 slices. |
| *COGNORM* | Siemens Avanto | | MPRAGE. TR: 2400 ms; TE: 3.79 ms, TI: 1000 ms; flip angle 8°, slice thickness: 1.2 mm, FoV 240 x 240, 160 slices. |
| | Siemens Prisma | 3.0 | MPRAGE. TR: 2400 ms; TE: 2.22 ms, TI: 1000 ms; flip angle 8°, slice thickness: 0.8 mm, FoV 240 x 256, 208 slices, iPat = 2. |
| *HABS*[c] | Tim Trio Siemens | 3.0 | MPRAGE. TR: 2300 ms; TE: 2.98 ms, TI: 900 ms; flip angle 9°, slice thickness: 1.2 mm, FoV 240 x 256, 160 slices. MPRAGE. TR: 2200 ms; TE: 1.5/3.4/5.2/7.0 ms, TI: 1100 ms; flip angle 7°, slice thickness: 1.2 mm, FoV: 228 x 228, 144 slices, Multi-echo = x4. |
| *LCBC* | Avanto Siemens | 1.5 | MPRAGE. TR: 2400 ms; TE: 3.79 ms, TI: 1000 ms; flip angle 8°, slice thickness: 1.2 mm, FoV 240 x 240, 160 slices. |
| | Siemens Skyra | 3.0 | MPRAGE. TR: 2300 ms; TE: 2.98 ms, TI: 850 ms; flip angle 8°, slice thickness: 1 mm, FoV: 256 x 256, 176 slices. |
| | Siemens Prisma | 3.0 | MPRAGE. TR: 2400 ms; TE: 2.22 ms, TI: 1000 ms; flip angle 8°, slice thickness: 0.8 mm, FoV 240 x 256, 208 slices, iPat = 2. |
| *OASIS3* | Siemens Vision | 1.5 | MPRAGE. TR: 9,7 ms; TE: 4.0 ms, TI: 20 ms; flip angle 10°, slice thickness: 1.25 mm, FoV: 256 x 256, 160 slices. |
| | Siemens Sonata | 1.5 | MPRAGE. TR: 9,7 ms; TE: 3.9 ms, TI: 20 ms; flip angle 15°, slice thickness: 1 mm, FoV: 224 x 256, 160 slices. |
| | Tim Trio Siemens | 3.0 | MPRAGE. TR: 2400 ms; TE: 3.1 ms, TI: 1000 ms; flip angle 8°, slice thickness: 1 mm, FoV 256 x 256, 176 slices. |
| | Magnetom Vida Siemens | 3.0 | MPRAGE. TR: 2300 ms; TE: 2.3 ms, TI: 900 ms; flip angle 9°, slice thickness: 1.2 mm, FoV 240 x 256, 176 slices. |
| | Siemens BioGraph mMR | 3.0 | MPRAGE. TR: 2300 ms; TE: 2.3 ms, TI: 900 ms; flip angle 9°, slice thickness: 1.2 mm, FoV 240 x 256, 176 slices. |
| *preventAD* | Tim Trio Siemens | 3.0 | MPRAGE. TR: 2300 ms; TE: 2.98 ms, TI: 900 ms; flip angle 9°, slice thickness: 1 mm, FoV 240 x 256, 176 slices. |
| *UB* | Tim trio Siemens | 3.0 | MPRAGE. TR: 2400 ms; TE: 2.98 ms, TI: 900 ms; flip angle 9°, slice thickness: 1 mm, FoV: 256 x 256, 240 slices. |
| *UKB* | Siemens Skyra[a] | 3.0 | MPRAGE. TR: 2000 ms; TE: - ms, TI: 880 ms; flip angle -, slice thickness: 1 mm, FoV: 208 x 256, 256 slices. |
| *Umeå* | Discovery GE | 3.0 | 3D FSPGR. TR: 8.19 ms; TE: 3.2ms, TI: 450 ms; flip angle 12°, slice thickness: 1 mm, FoV: 250 x 250, 180 slices. |
| *Wayne* | Siemens Sonata Magnetom | 1.5 | MPRAGE. TR: 800 ms; TE: 3.93 ms, TI: 420 ms; flip angle 20°, slice thickness: 1.5 mm, FoV 192 x 192, 144 slices. |

| | | | |
|---|---|---|---|
| | 4T-System Bruker Bio-spin | 4.0 | MPRAGE. TR: 1600 ms; TE: 4.38 ms, TI: 800 ms; flip angle 8°, slice thickness: 1.34 mm, FoV 256 x 256, 176 slices. |
| | Test-retest (reliability) dataset | | |
| *GSP* | Tim Trio Siemens[b] | 3.0 | MPRAGE. TR: 2200 ms; TE: 1.5/3.4/5.2/7.0 ms, TI: 1100 ms; flip angle 7°, slice thickness: 1.2 mm, FoV: 228 x 228, 144 slices, Multi-echo = x4. |
| *HNU1* | Discovery MR750 GE | 3.0 | 3D FSPGR. TR: 8.1 ms; TE: 3.1 ms, TI: 450 ms; flip angle 8°, slice thickness: 1 mm, FoV: 256 x 256, 176 slices. |
| *Maclaren* | Discovery MR750 GE | 3.0 | 3D IR-SPGR. TR: 7.30 ms; TE: 3.0 ms, TI: 400 ms; flip angle 12°, slice thickness: 1.2 mm, FoV: 256 x 256, 225 slices. |
| *preventAD* | Tim Trio Siemens | 3.0 | MPRAGE. TR: 2300 ms; TE: 2.98 ms, TI: 900 ms; flip angle 9°, slice thickness: 1 mm, FoV 240 x 256, 176 slices. |
| *OASIS* | Siemens Vi-sion | 1.5 | MPRAGE. TR: 9,7 ms; TE: 4.0 ms, TI: 20 ms; flip angle 10°, slice thickness: 1.25 mm, FoV: 256 x 256, 160 slices. |
| *S2C* | Siemens Prisma | 3.0 | MPRAGE. TR: 2400 ms; TE: 2.22 ms, TI: 1000 ms; flip angle 8°, slice thickness: 0.8 mm, FoV 240 x 256, 208 slices, iPat = 2. |

***Supplementary Table 6**. **Scanner acquisition parameters.** TR = Repetition Time; TE = Echo Time; TI = inversion time; FoV = Field of View, iPat = in-plane acceleration. [a,c]Two matched scanners. [b]Several matched scanners.*

#### Reliability of longitudinal brain change: Estimations based on empirical data.

We used the multicohort described in the slope variance section to estimate reliability empirically. Slope variance was considered fixed and estimated as described above while error variance of the slopes was assessed using the standard error of the slope for each individual and feature. That is, for each individual (i) with 3 or more observations, a linear model fitting the feature (j) values with time (from baseline) will provide the βeta coefficient for the slope as well as an associated standard error which we will refer as SE($\Delta_{i,j}$)). This measure SE($\Delta_{i,j}$)) is an indicator of the degree of uncertainty in the fitting of the slope. For each feature, the standard error of the slopes SE($\Delta_{i,j}$)) were fitted by total follow-up time, the number of observations, and its interaction with cohort as random intercept using generalized linear mixed effects models with a logarithmic link (*glmer*, *lme4* R-package) (Bates et al., 2015).

The predictions were corrected by the number of observations as they slightly underestimated the error variance of the slope. The standard error of the slopes SE($\Delta_{i,j}$)) slightly underestimate the error variance as it reflects the uncertainty with respect to the *observed* rather than the *real* slope. To estimate the relationship between SE($\Delta_{i,j}$)) and error we estimated reliability using a synthetic dataset. The repeated measures for each individual (i) measured at a given set of occasions ($t_1$, …, $t_n$), can be expressed as follows (**eq.1 main text**) and change can be expressed in terms of linear trends and captured by slopes. We assume that the slopes of change ($\beta_{2i}$) are normally distributed in the population with mean δ and variance $\sigma_S^2$ and similarly for (cross-sectional) measurement errors ($\varepsilon_{i,j}$) with a mean equal to 0 and variance $\sigma_\varepsilon^2$.The intercepts ($\beta_{1i}$) are also assumed to be normally distributed with a non-zero mean and variance. Effectively, the reliability of brain atrophy from this dataset (by comparing the observed values, but sampling n error distributions is identical to the one obtained using the formula outlined in the main text (**eq.5 main text**) (**Supplementary Figure 6a**). This procedure enables estimation of both the standard error of the slopes SE($\Delta_{i,j}$)) and error as we know the *real* slopes for a given individual (**Supplementary Figure 6b**). The relationship between both is dependent on the number of observations (**Supplementary Figure 6c**). The predictions were corrected using linear models. Note that one could assume varying slope variance by estimating global variance as a function of study duration and the number of observations and correcting for error variance using, for example, Generalized additive models for location, scale, and shape (*gamlss*) (R. A. Rigby and D. M. Stasinopoulos, 2005). However, in our experience, sample requirements are higher to obtain robust estimates.

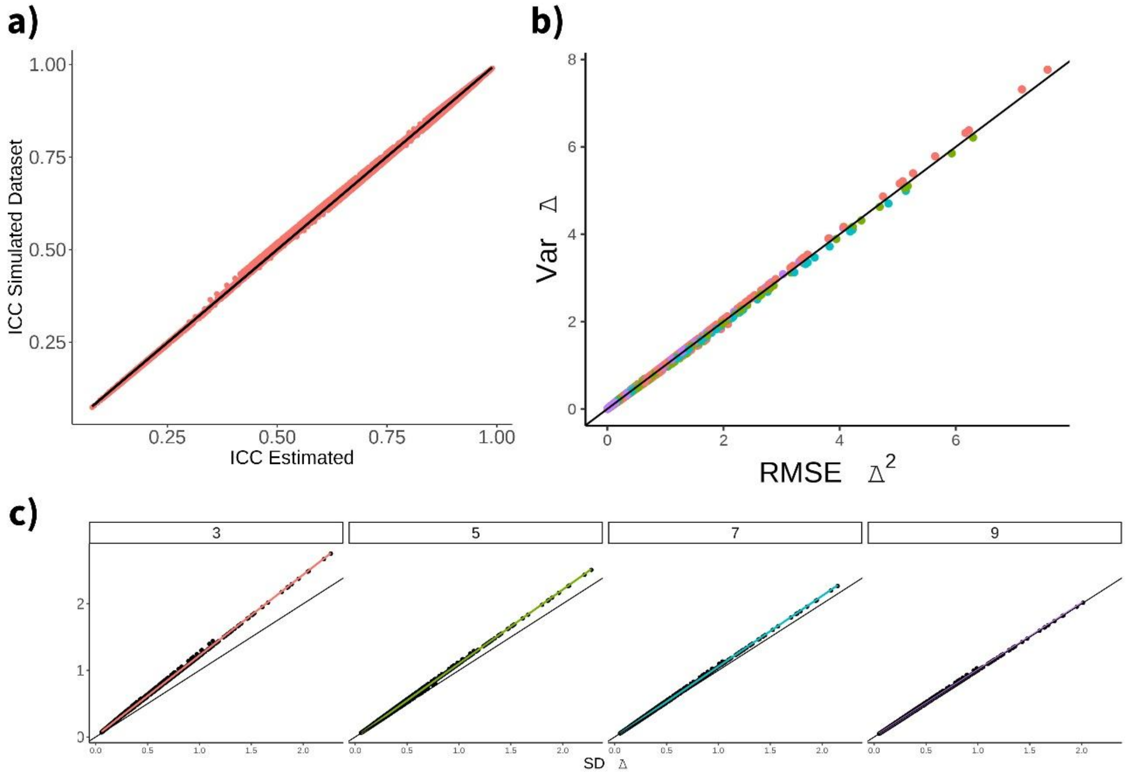


***Supplementary Figure 6. Relationship between SE of the slope and error.*** *A) Simulated datasets can be used to estimate the reliability of brain change producing nearly-equivalent estimates compared to the procedure outlined in the main text. The simulated datasets allow comparison between b) standard error of the slopes and within-subject error which c) is dependent on the number of observations.*

#### Supplementary Results

##### Reliability of longitudinal brain change.

| *term* | *Df* | *F-stat* | *p-value* | *partial Eta squared (η2)* |
|---:|---|---|---|---|
| *Mod* | 3 | 592.711 | 0.00 | 0.257 |
| *Obs* | 3 | 663.030 | 0.00 | 0.279 |
| *Follow-Up* | 5 | 15565.342 | 0.00 | 0.938 |
| *Mod×Obs* | 9 | 1.034 | 0.41 | 0.002 |
| *Mod×Follow-Up* | 15 | 20.887 | 0.00 | 0.057 |
| *Obs×Follow-Up* | 15 | 14.709 | 0.00 | 0.041 |
| *Mod×Obs×Follow-Up* | 45 | 0.312 | 1.00 | 0.003 |

***Supplementary Table 7**. **ANOVA summary**. Modality × Follow-up Time × Number of Observations on reliability of brain change. See also **Figure 2** and **Supplementary Figure 7** for visualization and **Supplementary Table 8** for additional statistics. Mod = Modality. Obs. = Number of Observations.*

| ***Modality*** | ***Follow-Up*** | ***Obs = 3*** | | ***Obs = 5*** | | ***Obs = 7*** | | ***Obs = 9*** | |
|---|---|---|---|---|---|---|---|---|---|
| | | *mean* | *SD* | *mean* | *SD* | *mean* | *SD* | *mean* | *SD* |
| *area* | 2 | 0.22 | 0.07 | 0.26 | 0.08 | 0.30 | 0.09 | 0.34 | 0.09 |
| | 4 | 0.51 | 0.11 | 0.57 | 0.11 | 0.62 | 0.11 | 0.66 | 0.10 |
| | 6 | 0.69 | 0.10 | 0.74 | 0.09 | 0.78 | 0.08 | 0.81 | 0.07 |
| | 8 | 0.80 | 0.08 | 0.83 | 0.07 | 0.86 | 0.06 | 0.88 | 0.05 |
| | 10 | 0.86 | 0.06 | 0.88 | 0.05 | 0.90 | 0.04 | 0.92 | 0.04 |
| | 12 | 0.90 | 0.05 | 0.92 | 0.04 | 0.93 | 0.03 | 0.94 | 0.03 |
| *subcortical* | 2 | 0.32 | 0.15 | 0.37 | 0.15 | 0.41 | 0.16 | 0.46 | 0.16 |
| | 4 | 0.63 | 0.14 | 0.67 | 0.13 | 0.72 | 0.12 | 0.75 | 0.11 |
| | 6 | 0.78 | 0.10 | 0.82 | 0.08 | 0.85 | 0.07 | 0.87 | 0.06 |
| | 8 | 0.86 | 0.07 | 0.89 | 0.06 | 0.91 | 0.05 | 0.92 | 0.04 |
| | 10 | 0.91 | 0.05 | 0.92 | 0.04 | 0.94 | 0.03 | 0.95 | 0.03 |
| | 12 | 0.93 | 0.03 | 0.94 | 0.03 | 0.95 | 0.02 | 0.96 | 0.02 |
| *thickness* | 2 | 0.15 | 0.04 | 0.18 | 0.05 | 0.22 | 0.05 | 0.25 | 0.06 |
| | 4 | 0.41 | 0.07 | 0.47 | 0.07 | 0.52 | 0.07 | 0.57 | 0.07 |
| | 6 | 0.61 | 0.06 | 0.66 | 0.06 | 0.71 | 0.05 | 0.74 | 0.05 |
| | 8 | 0.73 | 0.05 | 0.78 | 0.04 | 0.81 | 0.04 | 0.84 | 0.03 |
| | 10 | 0.81 | 0.04 | 0.84 | 0.03 | 0.87 | 0.03 | 0.89 | 0.02 |
| | 12 | 0.86 | 0.03 | 0.89 | 0.03 | 0.91 | 0.02 | 0.92 | 0.02 |
| *volume* | 2 | 0.17 | 0.03 | 0.20 | 0.04 | 0.24 | 0.05 | 0.27 | 0.05 |
| | 4 | 0.44 | 0.06 | 0.50 | 0.06 | 0.55 | 0.06 | 0.59 | 0.06 |
| | 6 | 0.64 | 0.06 | 0.69 | 0.06 | 0.73 | 0.05 | 0.77 | 0.05 |
| | 8 | 0.76 | 0.05 | 0.79 | 0.04 | 0.83 | 0.04 | 0.85 | 0.03 |
| | 10 | 0.83 | 0.04 | 0.86 | 0.03 | 0.88 | 0.03 | 0.90 | 0.02 |
| | 12 | 0.87 | 0.03 | 0.90 | 0.02 | 0.91 | 0.02 | 0.93 | 0.02 |

***Supplementary Table 8*. *Cell means***. *Average reliability of brain change in the different Modality × Follow-up Time × Number of Observations cells. See also* ***Figure 2*** *and* ***Supplementary Figure 7*** *for visualization and* ***Supplementary Table 7*** *for additional statistics. Obs. = Number of Observations.*

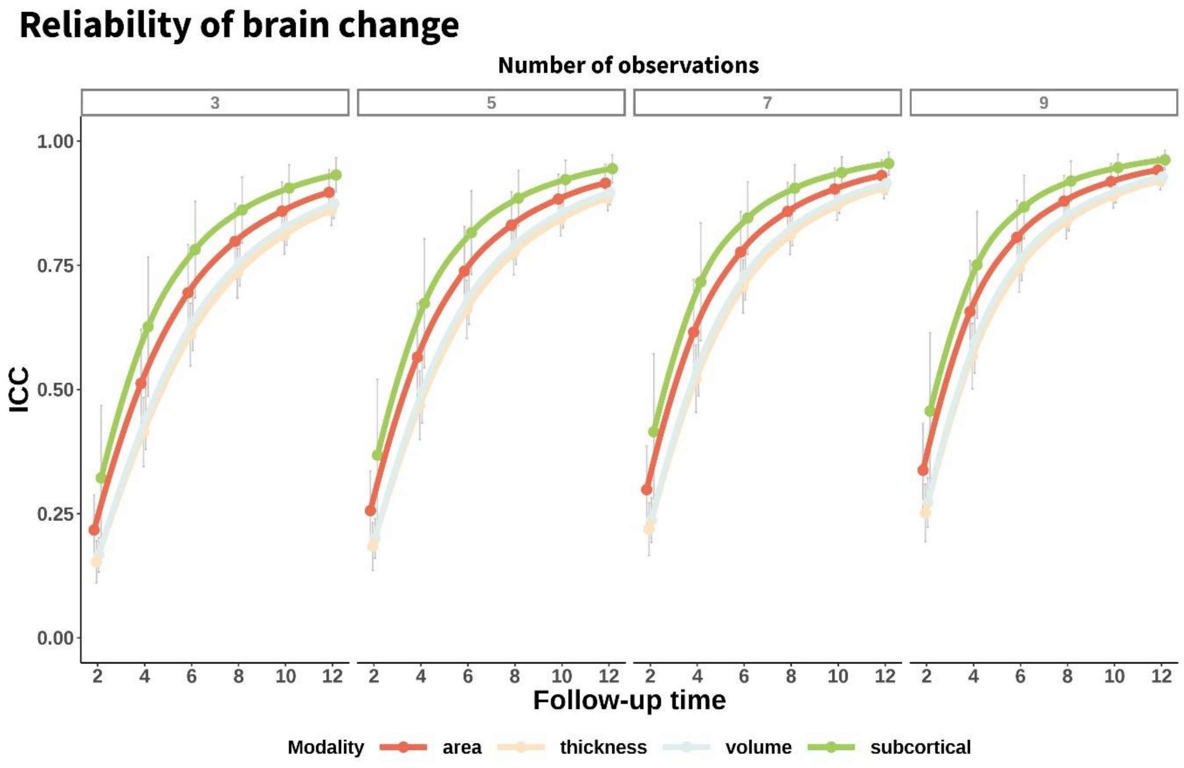


***Supplementary Figure 7. Reliability of brain change.*** *Mean – across features - reliability (ICC) of structural brain change as a function of total follow-up time, modality, and number of equispaced observations. Error bars represent 1 SD. See also* ***Figure 2*** *and* ***Supplementary Tables 7, 8*** *for additional statistics.*

#### Consistency of parameters and reliability estimates across datasets.

| Parameter | Modality | Type | ICC | lb | ub |
|---|---|---|---|---|---|
| *error (ε)* | volume | icc(2,1) | 0.54 | 0.44 | 0.64 |
| *error (ε)* | volume | icc(2,k) | 0.88 | 0.82 | 0.91 |
| *error (ε)* | area | icc(2,1) | 0.78 | 0.72 | 0.83 |
| *error (ε)* | area | icc(2,k) | 0.95 | 0.94 | 0.97 |
| *error (ε)* | thickness | icc(2,1) | 0.39 | 0.29 | 0.50 |
| *error (ε)* | thickness | icc(2,k) | 0.80 | 0.71 | 0.86 |
| *error (ε)* | subcortical | icc(2,1) | 0.63 | 0.44 | 0.80 |
| *error (ε)* | subcortical | icc(2,k) | 0.91 | 0.83 | 0.96 |
| *slope dis. (SD(δ))* | volume | icc(2,1) | 0.69 | 0.57 | 0.78 |
| *slope dis. (SD(δ))* | volume | icc(2,k) | 0.94 | 0.90 | 0.96 |
| *slope dis. (SD(δ))* | area | icc(2,1) | 0.81 | 0.72 | 0.87 |
| *slope dis. (SD(δ))* | area | icc(2,k) | 0.97 | 0.95 | 0.98 |
| *slope dis. (SD(δ))* | thickness | icc(2,1) | 0.52 | 0.36 | 0.65 |
| *slope dis. (SD(δ))* | thickness | icc(2,k) | 0.88 | 0.80 | 0.93 |
| *slope dis. (SD(δ))* | subcortical | icc(2,1) | 0.91 | 0.84 | 0.96 |
| *slope dis. (SD(δ))* | subcortical | icc(2,k) | 0.99 | 0.97 | 0.99 |
| *error (ε)* | all | icc(2,1) | 0.65 | 0.60 | 0.70 |
| *error (ε)* | all | icc(2,k) | 0.92 | 0.90 | 0.93 |
| *slope dis. (SD(δ))* | all | icc(2,1) | 0.82 | 0.76 | 0.87 |
| *slope dis. (SD(δ))* | all | icc(2,k) | 0.97 | 0.96 | 0.98 |

***Supplementary Table 9. Consistency of slope of parameters across datasets.*** *Consistency of the slope dispersion and cross-sectional measurement error parameters across datasets grouped within modality. Pairwise and global agreement. error (ε) = cross-sectional measurement error. slope dis. (SD(δ)) = slope dispersion; i.e. dispersion of the brain atrophy (standard deviation). lb = lower-bound; ub = upper bound (CI = 95%). See also* ***Supplementary Figure 8*** *for visualization.*

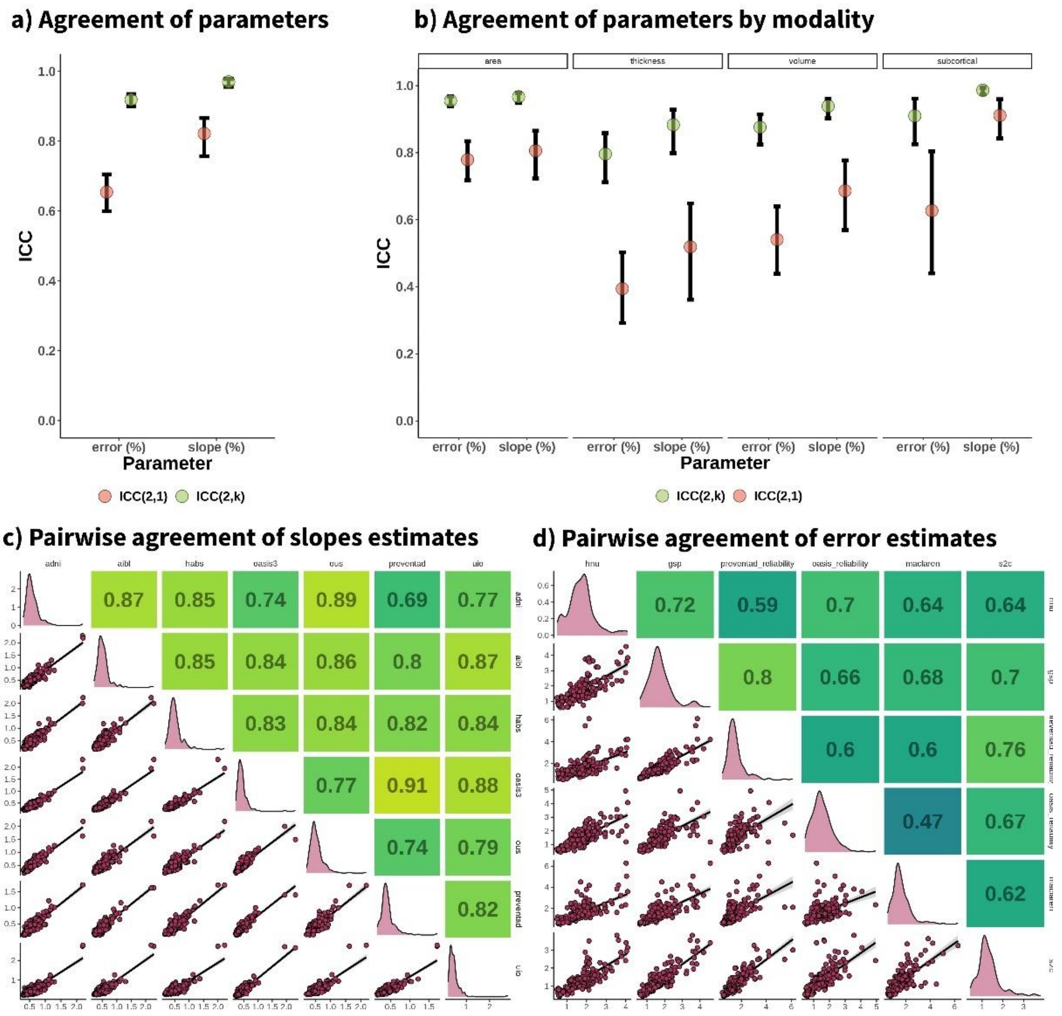


***Supplementary Figure 8. Consistency of slope of parameters across datasets.*** *Consistency of the slope dispersion and cross-sectional measurement error parameters across datasets a) across all features and b) grouped within modality. Pairwise and global agreement is shown. error(%) = cross-sectional measurement error. Slope(%) = slope dispersion; i.e. dispersion of the brain atrophy (standard deviation). Bars represent confident intervals (CI = 95%). Pairwise agreement of c) slope dispersion and d) cross-sectional measurement error. The lower triangle shows the correlation across features, the upper triangle provides the pairwise reliability for each dataset pair (ICC(2,1)). See also* ***Supplementary Table 9*** *for statistics.*

| *Follow-up* | *Type* | *Obs = 3* | | *Obs = 5* | | *Obs = 7* | | *Obs = 9* | |
|---|---|---|---|---|---|---|---|---|---|
| | | **mean** | **SD** | **mean** | **SD** | **mean** | **SD** | **mean** | **SD** |
| *2* | ICC(2,1) | 0.34 | 0.13 | 0.33 | 0.13 | 0.32 | 0.13 | 0.31 | 0.13 |
| *4* | ICC(2,1) | 0.27 | 0.14 | 0.26 | 0.13 | 0.25 | 0.13 | 0.24 | 0.13 |
| *6* | ICC(2,1) | 0.24 | 0.13 | 0.23 | 0.13 | 0.22 | 0.13 | 0.21 | 0.13 |
| *8* | ICC(2,1) | 0.21 | 0.13 | 0.21 | 0.13 | 0.20 | 0.13 | 0.20 | 0.13 |
| *10* | ICC(2,1) | 0.20 | 0.13 | 0.19 | 0.13 | 0.19 | 0.12 | 0.19 | 0.12 |
| *12* | ICC(2,1) | 0.19 | 0.13 | 0.19 | 0.12 | 0.18 | 0.12 | 0.18 | 0.12 |
| *2* | ICC(2,k) | 0.74 | 0.03 | 0.73 | 0.03 | 0.73 | 0.04 | 0.72 | 0.04 |
| *4* | ICC(2,k) | 0.68 | 0.04 | 0.67 | 0.04 | 0.65 | 0.04 | 0.64 | 0.04 |
| *6* | ICC(2,k) | 0.63 | 0.04 | 0.62 | 0.04 | 0.61 | 0.04 | 0.60 | 0.04 |
| *8* | ICC(2,k) | 0.60 | 0.04 | 0.59 | 0.04 | 0.58 | 0.04 | 0.57 | 0.04 |
| *10* | ICC(2,k) | 0.58 | 0.04 | 0.57 | 0.04 | 0.56 | 0.05 | 0.55 | 0.05 |
| *12* | ICC(2,k) | 0.56 | 0.05 | 0.55 | 0.05 | 0.54 | 0.05 | 0.53 | 0.05 |

***Supplementary Table 10. Regional consistency of reliability estimates across datasets.*** *Regional consistency of the reliability estimates across datasets. Different combinations of datasets for deriving slope dispersion and cross-sectional measurement error parameters were used and reliability was assessed within a given number of observations and follow-up time. Pairwise and global agreement. See also* ***Supplementary Figure 9*** *for visualization.*

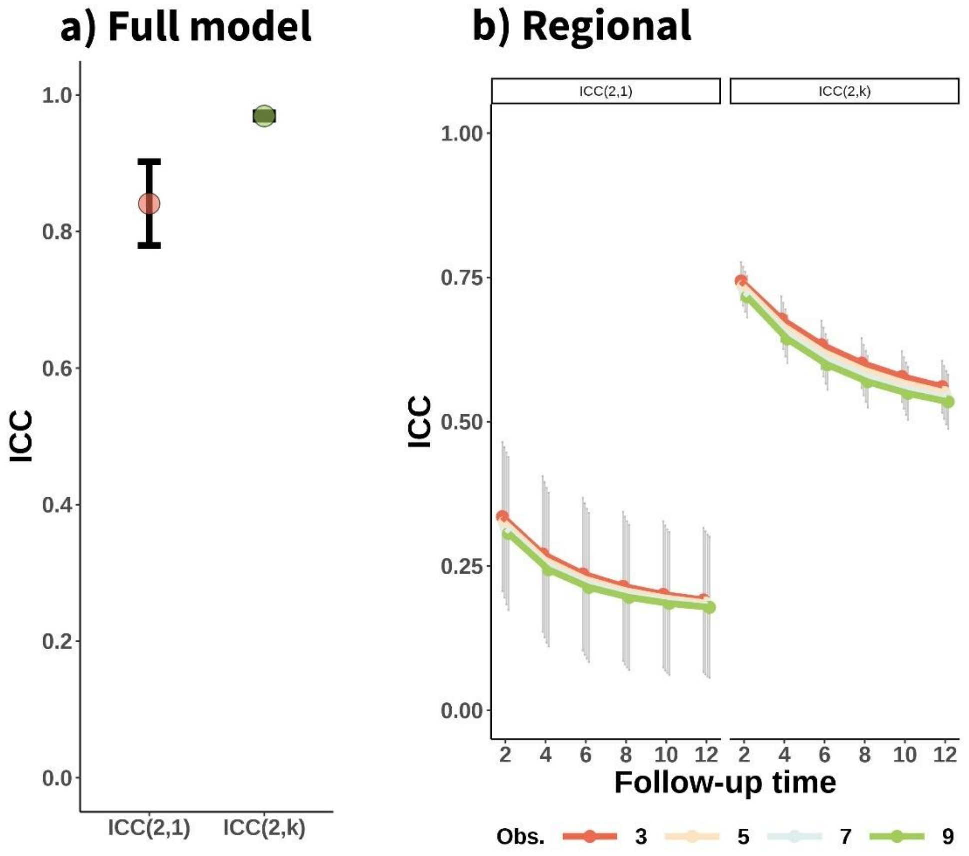


***Supplementary Figure 9. Regional consistency of reliability estimates across datasets.*** *Regional consistency of the reliability estimates across datasets. Different combinations of datasets for deriving slope dispersion and cross-sectional measurement error parameters were used and reliability was assessed a) across all number of observations and follow-up time and b) within a given number of observations and follow-up time. Pairwise and global agreement. See also **Supplementary Table 10** for visualization.*

#### Consequences of longitudinal reliability (I): Sample Size Estimates.

| ***Follow-up*** | ***Pearson's r*** | ***Obs = 3*** | | ***Obs = 5*** | | ***Obs = 7*** | | ***Obs = 9*** | |
|---|---|---|---|---|---|---|---|---|---|
| | | **mean** | **SD** | **mean** | **SD** | **mean** | **SD** | **mean** | **SD** |
| *2* | 0.1 | 4685 | 1497 | 3905 | 1198 | 3292 | 962 | 2864 | 799 |
| *4* | 0.1 | 1758 | 375 | 1563 | 300 | 1410 | 241 | 1303 | 200 |
| *6* | 0.1 | 1216 | 167 | 1130 | 133 | 1062 | 107 | 1014 | 89 |
| *8* | 0.1 | 1027 | 94 | 978 | 75 | 940 | 61 | 913 | 50 |
| *10* | 0.1 | 939 | 60 | 908 | 48 | 883 | 39 | 866 | 32 |
| *12* | 0.1 | 891 | 42 | 869 | 34 | 852 | 27 | 841 | 23 |
| *2* | 0.3 | 519 | 167 | 432 | 134 | 364 | 107 | 316 | 89 |
| *4* | 0.3 | 194 | 42 | 172 | 34 | 155 | 27 | 143 | 23 |
| *6* | 0.3 | 133 | 19 | 124 | 15 | 116 | 12 | 111 | 10 |
| *8* | 0.3 | 112 | 11 | 107 | 9 | 102 | 7 | 100 | 6 |
| *10* | 0.3 | 102 | 7 | 99 | 6 | 96 | 5 | 94 | 4 |
| *12* | 0.3 | 97 | 5 | 95 | 4 | 93 | 3 | 92 | 3 |
| *2* | 0.5 | 185 | 60 | 154 | 48 | 130 | 39 | 113 | 32 |
| *4* | 0.5 | 68 | 15 | 60 | 12 | 54 | 10 | 50 | 8 |
| *6* | 0.5 | 47 | 7 | 43 | 6 | 40 | 5 | 38 | 4 |
| *8* | 0.5 | 39 | 4 | 37 | 3 | 36 | 3 | 34 | 3 |
| *10* | 0.5 | 36 | 3 | 34 | 2 | 33 | 2 | 33 | 2 |
| *12* | 0.5 | 34 | 2 | 33 | 2 | 32 | 2 | 32 | 1 |

***Supplementary Table 11. Sample size estimates.*** *Mean sample size estimates for achieving 80% power $p < 0.05$, as a function of Pearson's correlations strength (r), follow-up time, and number of observations. Mean sample size across cortical volume, area, and thickness, and subcortical volume features. Obs. = Number of observations per participant. See also* ***Figure 3a*** *for visualization.*

| ***Follow-up*** | ***Pearson's r*** | ***Left Hippocampus*** | | | | ***Left Entorhinal Thickness*** | | | |
|---|---|---|---|---|---|---|---|---|---|
| | | **Obs = 3** | **Obs = 5** | **Obs = 7** | **Obs = 9** | **Obs = 3** | **Obs = 5** | **Obs = 7** | **Obs = 9** |
| *2* | 0.1 | 2427 | 2098 | 1840 | 1660 | 4097 | 3434 | 2913 | 2550 |
| *4* | 0.1 | 1193 | 1111 | 1047 | 1002 | 1611 | 1445 | 1315 | 1224 |
| *6* | 0.1 | 965 | 928 | 900 | 880 | 1151 | 1077 | 1019 | 979 |
| *8* | 0.1 | 885 | 864 | 848 | 837 | 989 | 948 | 915 | 893 |
| *10* | 0.1 | 848 | 835 | 825 | 817 | 915 | 888 | 867 | 853 |
| *12* | 0.1 | 828 | 819 | 812 | 807 | 874 | 856 | 841 | 831 |
| *2* | 0.3 | 267 | 231 | 202 | 182 | 453 | 379 | 321 | 281 |
| *4* | 0.3 | 130 | 121 | 114 | 109 | 177 | 158 | 144 | 134 |
| *6* | 0.3 | 105 | 101 | 98 | 95 | 125 | 117 | 111 | 106 |
| *8* | 0.3 | 96 | 94 | 92 | 91 | 108 | 103 | 99 | 97 |
| *10* | 0.3 | 92 | 90 | 89 | 88 | 99 | 96 | 94 | 92 |
| *12* | 0.3 | 90 | 89 | 88 | 87 | 95 | 93 | 91 | 90 |
| *2* | 0.5 | 95 | 81 | 71 | 64 | 161 | 135 | 114 | 99 |
| *4* | 0.5 | 45 | 42 | 39 | 38 | 62 | 55 | 50 | 46 |
| *6* | 0.5 | 36 | 35 | 33 | 33 | 43 | 41 | 38 | 37 |
| *8* | 0.5 | 33 | 32 | 31 | 31 | 37 | 35 | 34 | 33 |
| *10* | 0.5 | 31 | 31 | 30 | 30 | 34 | 33 | 32 | 32 |
| *12* | 0.5 | 31 | 30 | 30 | 30 | 32 | 32 | 31 | 31 |

***Supplementary Table 12. Sample size estimates for specific features.*** *Sample size estimates for achieving 80% power $p < 0.05$, as a function of Pearson's correlations strength (r), follow-up time, and number of observations in the Left Hippocampus and Entorhinal Thickness features. Obs. = Number of observations per participant. See also* ***Figure 3b, c*** *for visualization.*

#### Consequences of longitudinal reliability (II): Misclassification of individual trajectories.

| *Follow-Up* | *Obs.* | ***Bhattacharyya coefficient*** | | | | ***Probability of misclassification*** | | | |
|---|---|---|---|---|---|---|---|---|---|
| | | **Main. vs. Decliner** | | **Main./Decliner vs. Normal** | | **Maint. vs. decliner** | | **Main./Decliner vs. Normal** | |
| | | **Mean** | **SD** | **mean** | **SD** | **mean** | **SD** | **mean** | **SD** |
| *2* | 3 | 0.89 | 0.06 | 0.97 | 0.02 | 0.18 | 0.05 | 0.32 | 0.04 |
| *2* | 5 | 0.86 | 0.07 | 0.96 | 0.02 | 0.15 | 0.05 | 0.30 | 0.04 |
| *2* | 7 | 0.83 | 0.08 | 0.95 | 0.03 | 0.12 | 0.05 | 0.28 | 0.05 |
| *2* | 9 | 0.80 | 0.09 | 0.94 | 0.03 | 0.10 | 0.04 | 0.26 | 0.05 |
| *4* | 3 | 0.63 | 0.13 | 0.89 | 0.06 | 0.04 | 0.02 | 0.18 | 0.05 |
| *4* | 5 | 0.57 | 0.14 | 0.86 | 0.07 | 0.02 | 0.02 | 0.15 | 0.05 |
| *4* | 7 | 0.50 | 0.14 | 0.83 | 0.08 | 0.02 | 0.01 | 0.12 | 0.05 |
| *4* | 9 | 0.44 | 0.14 | 0.80 | 0.09 | 0.01 | 0.01 | 0.10 | 0.04 |
| *6* | 3 | 0.38 | 0.14 | 0.77 | 0.10 | 0.01 | 0.01 | 0.09 | 0.04 |
| *6* | 5 | 0.30 | 0.13 | 0.72 | 0.11 | 0.00 | 0.00 | 0.06 | 0.03 |
| *6* | 7 | 0.23 | 0.12 | 0.67 | 0.13 | 0.00 | 0.00 | 0.05 | 0.03 |
| *6* | 9 | 0.18 | 0.11 | 0.62 | 0.13 | 0.00 | 0.00 | 0.03 | 0.02 |
| *8* | 3 | 0.20 | 0.11 | 0.63 | 0.13 | 0.00 | 0.00 | 0.04 | 0.02 |
| *8* | 5 | 0.14 | 0.09 | 0.57 | 0.14 | 0.00 | 0.00 | 0.02 | 0.02 |
| *8* | 7 | 0.09 | 0.07 | 0.50 | 0.14 | 0.00 | 0.00 | 0.02 | 0.01 |
| *8* | 9 | 0.06 | 0.05 | 0.44 | 0.14 | 0.00 | 0.00 | 0.01 | 0.01 |
| *10* | 3 | 0.09 | 0.07 | 0.50 | 0.14 | 0.00 | 0.00 | 0.01 | 0.01 |
| *10* | 5 | 0.05 | 0.05 | 0.43 | 0.14 | 0.00 | 0.00 | 0.01 | 0.01 |
| *10* | 7 | 0.03 | 0.03 | 0.35 | 0.14 | 0.00 | 0.00 | 0.00 | 0.01 |
| *10* | 9 | 0.02 | 0.02 | 0.29 | 0.13 | 0.00 | 0.00 | 0.00 | 0.00 |
| *12* | 3 | 0.04 | 0.04 | 0.38 | 0.14 | 0.00 | 0.00 | 0.01 | 0.01 |
| *12* | 5 | 0.02 | 0.02 | 0.30 | 0.13 | 0.00 | 0.00 | 0.00 | 0.00 |
| *12* | 7 | 0.01 | 0.01 | 0.23 | 0.12 | 0.00 | 0.00 | 0.00 | 0.00 |
| *12* | 9 | 0.00 | 0.01 | 0.18 | 0.11 | 0.00 | 0.00 | 0.00 | 0.00 |

***Supplementary Table 13. Misclassification of individual trajectories.*** *Degree of overlap between distributions (Bhattacharyya coefficient) and the probability of misclassification (i.e., observing steeper slopes for a normal ager than for a decliner) normal, maintainer, and decliner participants whose brains change 0, -1, and 1 SD faster relative to the population average. Mean values across cortical volume, area, and thickness, and subcortical volume features. Obs. = Number of observations per participant. Main. = Maintainer. See also* ***Figure 4a*** *for visualization.*

| *Obs.* | *Follow-up* | *Left Hippocampus* | | | | *Left Entorhinal Thickness* | | | |
|---|---|---|---|---|---|---|---|---|---|
| | | **Bhattacharyya coefficient** | | **probability of misclassification** | | **Bhattacharyya coefficient** | | **probability of misclassification** | |
| | | **Maint. vs. Decl.** | **Maint./ Decl. vs. Normal** | **Maint. vs. Decl.** | **Maint./ Decl. vs. Normal** | **Maint. vs. Decl.** | **Maint./ Decl. vs. Normal** | **Maint. vs. Decl.** | **Maint./ Decl. vs. Normal** |
| *3* | 2 | 0.79 | 0.94 | 0.08 | 0.24 | 0.89 | 0.97 | 0.17 | 0.31 |
| *5* | 2 | 0.74 | 0.93 | 0.06 | 0.22 | 0.86 | 0.96 | 0.14 | 0.29 |
| *7* | 2 | 0.69 | 0.91 | 0.04 | 0.19 | 0.83 | 0.95 | 0.11 | 0.27 |
| *9* | 2 | 0.64 | 0.89 | 0.03 | 0.17 | 0.80 | 0.95 | 0.09 | 0.25 |
| *3* | 4 | 0.39 | 0.79 | 0.00 | 0.08 | 0.62 | 0.89 | 0.03 | 0.17 |
| *5* | 4 | 0.30 | 0.74 | 0.00 | 0.06 | 0.55 | 0.86 | 0.01 | 0.14 |
| *7* | 4 | 0.23 | 0.69 | 0.00 | 0.04 | 0.48 | 0.83 | 0.01 | 0.11 |
| *9* | 4 | 0.17 | 0.64 | 0.00 | 0.03 | 0.41 | 0.80 | 0.00 | 0.09 |
| *3* | 6 | 0.12 | 0.58 | 0.00 | 0.02 | 0.34 | 0.77 | 0.00 | 0.07 |
| *5* | 6 | 0.07 | 0.51 | 0.00 | 0.01 | 0.26 | 0.72 | 0.00 | 0.05 |
| *7* | 6 | 0.04 | 0.43 | 0.00 | 0.00 | 0.19 | 0.66 | 0.00 | 0.03 |
| *9* | 6 | 0.02 | 0.37 | 0.00 | 0.00 | 0.14 | 0.61 | 0.00 | 0.02 |
| *3* | 8 | 0.02 | 0.39 | 0.00 | 0.00 | 0.15 | 0.62 | 0.00 | 0.03 |
| *5* | 8 | 0.01 | 0.30 | 0.00 | 0.00 | 0.09 | 0.55 | 0.00 | 0.01 |
| *7* | 8 | 0.00 | 0.23 | 0.00 | 0.00 | 0.05 | 0.48 | 0.00 | 0.01 |
| *9* | 8 | 0.00 | 0.17 | 0.00 | 0.00 | 0.03 | 0.41 | 0.00 | 0.00 |
| *3* | 10 | 0.00 | 0.23 | 0.00 | 0.00 | 0.05 | 0.48 | 0.00 | 0.01 |
| *5* | 10 | 0.00 | 0.16 | 0.00 | 0.00 | 0.02 | 0.40 | 0.00 | 0.00 |
| *7* | 10 | 0.00 | 0.10 | 0.00 | 0.00 | 0.01 | 0.32 | 0.00 | 0.00 |
| *9* | 10 | 0.00 | 0.06 | 0.00 | 0.00 | 0.00 | 0.25 | 0.00 | 0.00 |
| *3* | 12 | 0.00 | 0.12 | 0.00 | 0.00 | 0.01 | 0.34 | 0.00 | 0.00 |
| *5* | 12 | 0.00 | 0.07 | 0.00 | 0.00 | 0.00 | 0.26 | 0.00 | 0.00 |
| *7* | 12 | 0.00 | 0.04 | 0.00 | 0.00 | 0.00 | 0.19 | 0.00 | 0.00 |
| *9* | 12 | 0.00 | 0.02 | 0.00 | 0.00 | 0.00 | 0.14 | 0.00 | 0.00 |

***Supplementary Table 14. Misclassification of individual trajectories for specific features.*** *Degree of overlap between distributions (Bhattacharyya coefficient) and the probability of misclassification (i.e., observing steeper slopes for a normal ager than for a decliner) normal, maintainer, and decliner participants whose brains change 0, -1, and 1 SD faster relative to the population average. Values for the Left Hippocampus and Entorhinal Thickness. Obs. = Number of observations per participant. Main. = Maintainer. Decl. = Decliner. See also* ***Figure 4b,c*** *for visualization.*

##### Consequences of longitudinal reliability (III): Group membership based on trajectories.

| ***Feature*** | ***Obs*** | ***Observed Maintainer (%)*** | | | | ***Maintainer \| Observed Maintainer (%)*** | | | | ***Decliner \| Observed Maintainer (%)*** | | | |
|---|---|---|---|---|---|---|---|---|---|---|---|---|---|
| *Obs.* | | 3 | 5 | 7 | 9 | 3 | 5 | 7 | 9 | 3 | 5 | 7 | 9 |
| *Left Hippocampus* | 2 | 0.16 | 0.14 | 0.13 | 0.11 | 0.15 | 0.18 | 0.20 | 0.23 | 0.15 | 0.12 | 0.09 | 0.06 |
| | 4 | 0.08 | 0.07 | 0.07 | 0.06 | 0.36 | 0.39 | 0.45 | 0.49 | 0.01 | 0.00 | 0.00 | 0.00 |
| | 6 | 0.06 | 0.05 | 0.05 | 0.05 | 0.53 | 0.56 | 0.61 | 0.65 | 0.00 | 0.00 | 0.00 | 0.00 |
| | 8 | 0.05 | 0.05 | 0.05 | 0.05 | 0.63 | 0.67 | 0.70 | 0.73 | 0.00 | 0.00 | 0.00 | 0.00 |
| | 10 | 0.05 | 0.05 | 0.04 | 0.04 | 0.71 | 0.75 | 0.76 | 0.79 | 0.00 | 0.00 | 0.00 | 0.00 |
| | 12 | 0.05 | 0.04 | 0.04 | 0.04 | 0.76 | 0.79 | 0.80 | 0.83 | 0.00 | 0.00 | 0.00 | 0.00 |
| *Left Entorhinal Thickness* | 2 | 0.38 | 0.37 | 0.36 | 0.35 | 0.39 | 0.41 | 0.43 | 0.45 | 0.32 | 0.30 | 0.27 | 0.25 |
| | 4 | 0.31 | 0.30 | 0.29 | 0.29 | 0.54 | 0.57 | 0.60 | 0.62 | 0.16 | 0.13 | 0.11 | 0.09 |
| | 6 | 0.28 | 0.28 | 0.27 | 0.27 | 0.65 | 0.67 | 0.71 | 0.73 | 0.07 | 0.06 | 0.04 | 0.03 |
| | 8 | 0.27 | 0.26 | 0.26 | 0.25 | 0.72 | 0.75 | 0.78 | 0.80 | 0.03 | 0.02 | 0.01 | 0.01 |
| | 10 | 0.26 | 0.25 | 0.25 | 0.25 | 0.78 | 0.80 | 0.82 | 0.83 | 0.01 | 0.01 | 0.00 | 0.00 |
| | 12 | 0.25 | 0.25 | 0.25 | 0.25 | 0.81 | 0.83 | 0.85 | 0.86 | 0.00 | 0.00 | 0.00 | 0.00 |

***Supplementary Table 15. Misclassification of group membership for specific features.*** *Probability of observing individuals without decline (i.e. to observe brain maintenance) and the conditional probability of them being "true" maintainers or individuals that decline faster than the population average. Probabilities are shown for the Left Hippocampus and Entorhinal Thickness. Obs. = Number of observations per participant. See also* ***Figure 5*** *for visualization.*

#### Determinants of longitudinal reliability (I): Sample characteristics affect longitudinal reliability.

| ***Age Group*** | ***Follow-Up*** | ***Left Entorhinal Thickness*** | | | | ***Left Hippocampus*** | | | |
|---|---|---|---|---|---|---|---|---|---|
| | | **Obs = 3** | **Obs = 5** | **Obs = 7** | **Obs = 9** | **Obs = 3** | **Obs = 5** | **Obs = 7** | **Obs = 9** |
| *Young (<60 yrs)* | 2 | 0.09 | 0.12 | 0.14 | 0.16 | 0.12 | 0.15 | 0.18 | 0.21 |
| | 4 | 0.29 | 0.34 | 0.39 | 0.44 | 0.36 | 0.41 | 0.46 | 0.51 |
| | 6 | 0.48 | 0.54 | 0.59 | 0.64 | 0.55 | 0.61 | 0.66 | 0.70 |
| | 8 | 0.62 | 0.68 | 0.72 | 0.76 | 0.69 | 0.73 | 0.77 | 0.81 |
| | 10 | 0.72 | 0.76 | 0.80 | 0.83 | 0.78 | 0.81 | 0.84 | 0.87 |
| | 12 | 0.79 | 0.82 | 0.85 | 0.88 | 0.83 | 0.86 | 0.89 | 0.90 |
| *Old (>60 yrs)* | 2 | 0.20 | 0.24 | 0.28 | 0.32 | 0.34 | 0.39 | 0.45 | 0.49 |
| | 4 | 0.50 | 0.56 | 0.61 | 0.65 | 0.67 | 0.72 | 0.76 | 0.79 |
| | 6 | 0.69 | 0.74 | 0.78 | 0.81 | 0.82 | 0.85 | 0.88 | 0.90 |
| | 8 | 0.80 | 0.83 | 0.86 | 0.88 | 0.89 | 0.91 | 0.93 | 0.94 |
| | 10 | 0.86 | 0.89 | 0.91 | 0.92 | 0.93 | 0.94 | 0.95 | 0.96 |
| | 12 | 0.90 | 0.92 | 0.93 | 0.94 | 0.95 | 0.96 | 0.97 | 0.97 |

***Supplementary Table 16*. *Longitudinal reliability across different age groups*.** *Reliability of left hippocampal atrophy and left entorhinal thinning as a function of follow-up time, number of observations, and age group. See* ***Figure 6*** *for visualization. Obs. = Number of Observations. Yrs. = Years.*

#### Determinants of longitudinal reliability (II): Preprocessing stream.

| term | df | F-stat | p-value | partial Eta squared (η2) |
|---|---|---|---|---|
| *Mod* | 3 | 252.468 | 0 | 0.129 |
| *Obs* | 3 | 263.614 | 0 | 0.133 |
| *Follow-Up* | 5 | 4087.557 | 0 | 0.799 |
| *Mod×Obs* | 9 | 0.064 | 1 | 0.000 |
| *Mod×Follow-Up* | 15 | 3.594 | 0 | 0.010 |
| *Obs×Follow-Up* | 15 | 4.666 | 0 | 0.013 |
| *Mod×Obs×Follow-Up* | 45 | 0.077 | 1 | 0.001 |

***Supplementary Table 17**. **ANOVA summary**. Modality × Follow-up Time × Number of Observations on the reliability of brain change estimated from data processed with the FreeSurfer Cross-Sectional Stream. See also **Figure 7** and **Supplementary Figure 10** for visualization and **Supplementary Table 18** for additional statistics. Mod = Modality. Obs. = Number of Observations.*

| *Modality* | *Follow-Up* | *Obs = 3* | | *Obs = 5* | | *Obs = 7* | | *Obs = 9* | |
|---|---|---|---|---|---|---|---|---|---|
| | | *mean* | *SD* | *mean* | *SD* | *mean* | *SD* | *mean* | *SD* |
| *area* | 2 | 0.05 | 0.03 | 0.06 | 0.04 | 0.08 | 0.05 | 0.09 | 0.05 |
| | 4 | 0.17 | 0.09 | 0.20 | 0.10 | 0.24 | 0.11 | 0.27 | 0.12 |
| | 6 | 0.31 | 0.13 | 0.35 | 0.14 | 0.40 | 0.14 | 0.44 | 0.15 |
| | 8 | 0.43 | 0.15 | 0.48 | 0.15 | 0.53 | 0.15 | 0.57 | 0.15 |
| | 10 | 0.53 | 0.15 | 0.58 | 0.15 | 0.63 | 0.15 | 0.66 | 0.14 |
| | 12 | 0.61 | 0.15 | 0.66 | 0.14 | 0.70 | 0.14 | 0.73 | 0.13 |
| *subcortical* | 2 | 0.14 | 0.15 | 0.16 | 0.16 | 0.19 | 0.18 | 0.21 | 0.19 |
| | 4 | 0.33 | 0.22 | 0.37 | 0.23 | 0.41 | 0.23 | 0.45 | 0.23 |
| | 6 | 0.48 | 0.22 | 0.53 | 0.22 | 0.57 | 0.21 | 0.61 | 0.20 |
| | 8 | 0.60 | 0.20 | 0.65 | 0.19 | 0.69 | 0.17 | 0.72 | 0.16 |
| | 10 | 0.69 | 0.17 | 0.73 | 0.16 | 0.77 | 0.14 | 0.79 | 0.13 |
| | 12 | 0.75 | 0.15 | 0.79 | 0.13 | 0.82 | 0.11 | 0.84 | 0.10 |
| *thickness* | 2 | 0.06 | 0.03 | 0.08 | 0.03 | 0.10 | 0.04 | 0.11 | 0.05 |
| | 4 | 0.21 | 0.07 | 0.25 | 0.08 | 0.29 | 0.09 | 0.33 | 0.09 |
| | 6 | 0.37 | 0.10 | 0.42 | 0.10 | 0.47 | 0.10 | 0.52 | 0.10 |
| | 8 | 0.50 | 0.10 | 0.56 | 0.10 | 0.61 | 0.09 | 0.65 | 0.09 |
| | 10 | 0.61 | 0.09 | 0.66 | 0.09 | 0.70 | 0.08 | 0.74 | 0.07 |
| | 12 | 0.69 | 0.08 | 0.73 | 0.08 | 0.77 | 0.07 | 0.80 | 0.06 |
| *volume* | 2 | 0.05 | 0.02 | 0.06 | 0.03 | 0.07 | 0.03 | 0.09 | 0.04 |
| | 4 | 0.16 | 0.07 | 0.20 | 0.08 | 0.23 | 0.09 | 0.27 | 0.10 |
| | 6 | 0.30 | 0.10 | 0.35 | 0.11 | 0.40 | 0.12 | 0.44 | 0.12 |
| | 8 | 0.43 | 0.12 | 0.48 | 0.12 | 0.53 | 0.12 | 0.57 | 0.12 |
| | 10 | 0.53 | 0.12 | 0.58 | 0.12 | 0.63 | 0.12 | 0.67 | 0.11 |
| | 12 | 0.61 | 0.12 | 0.66 | 0.11 | 0.71 | 0.11 | 0.74 | 0.10 |

***Supplementary Table 18*. *Cell means*.** *The average reliability of brain change from data obtained using the FreeSurfer cross-sectional stream in the different Modality × Follow-up Time × Number of Observations cells. See also* ***Figure 7*** *and* ***Supplementary Figure 10*** *for visualization and* ***Supplementary Table 17*** *for additional statistics. Obs. = Number of Observations.*

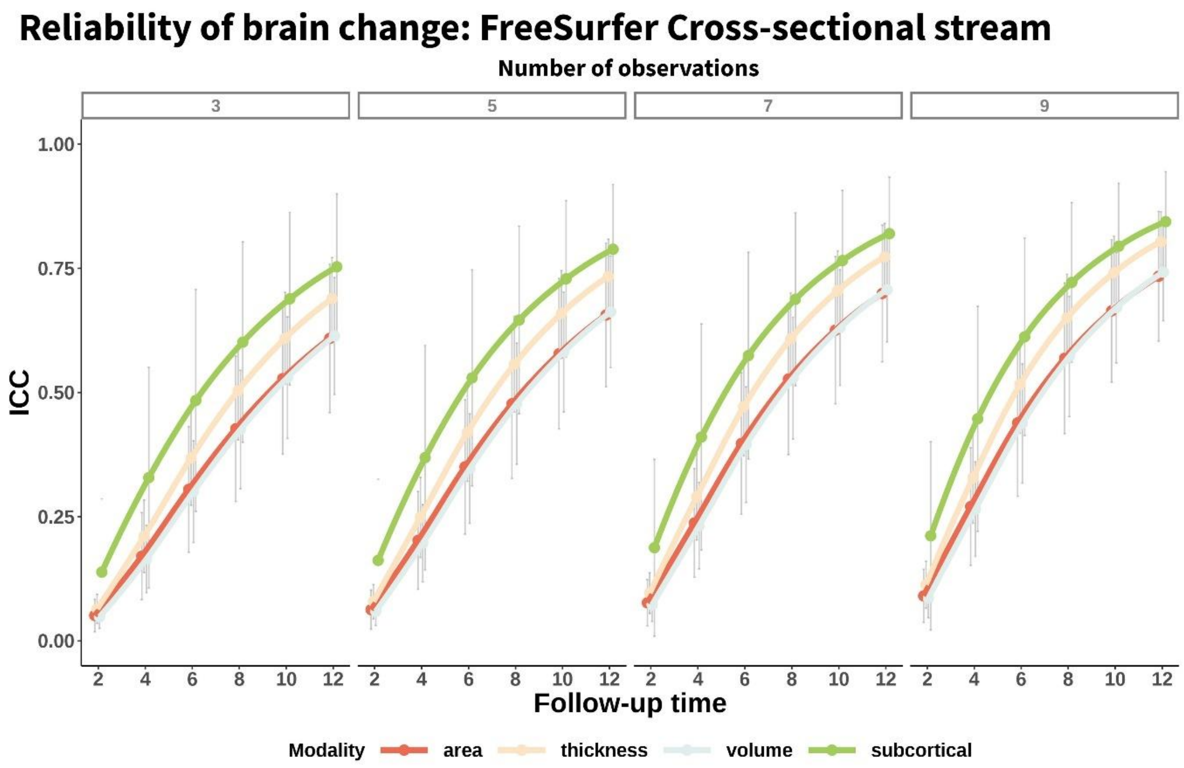


***Supplementary Figure 10. Reliability of brain change: FreeSurfer Cross-sectional Stream.*** *Mean – across features - reliability (ICC) of structural brain change as a function of total follow-up time, modality, and number of equispaced observations. Data was obtained from the FreeSurfer cross-sectional stream. Error bars represent 1 SD. See also* ***Figure 7*** *and* ***Supplementary Tables 17, 18*** *for additional statistics.*

#### Determinants of longitudinal reliability (III): Global versus regional features.

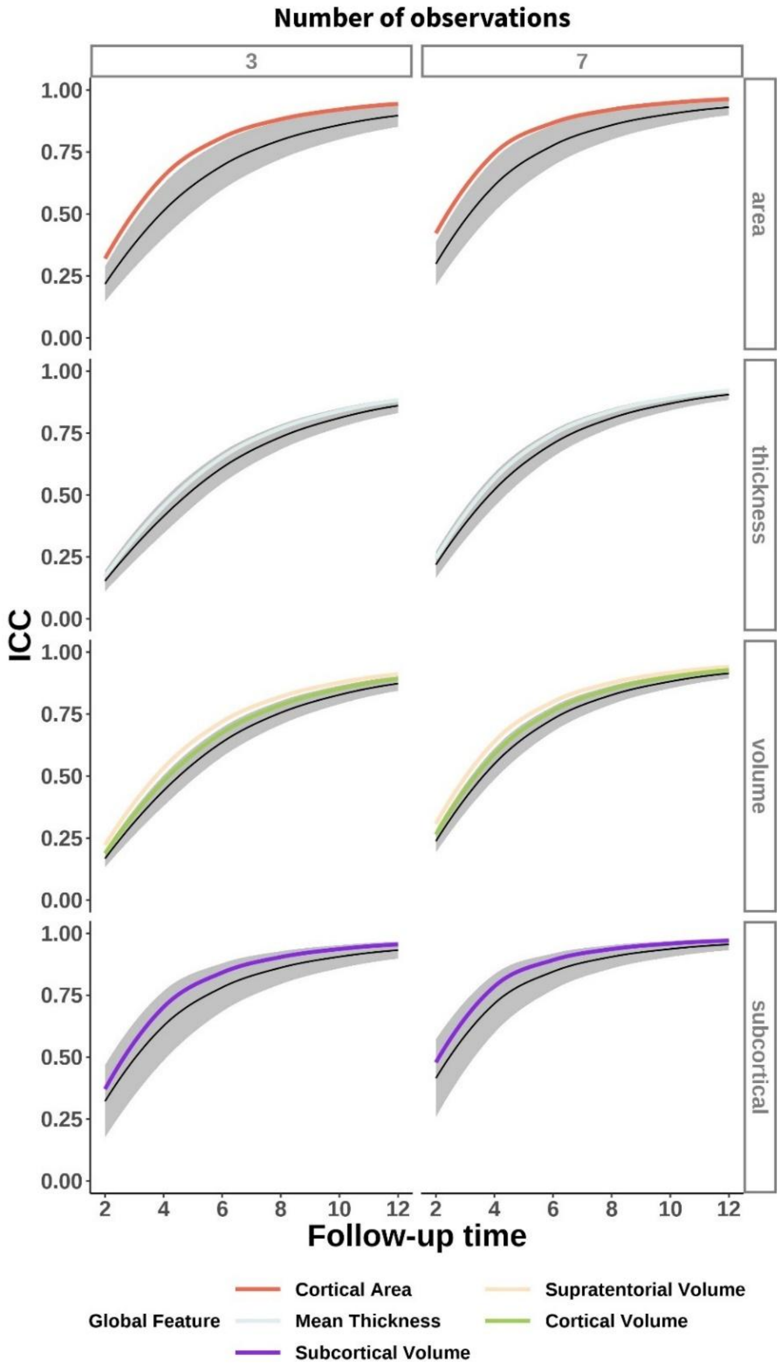


***Supplementary Figure 11.*** *Longitudinal reliability for whole-brain, summary features. Longitudinal reliability of whole-brain summary features overlaid with mean – across features, within modality – reliability as a function of total time and number of observations. The black line represents the mean and shades the SD. For visualization purposes, only 3 and 7 number of observations are shown.*

#### Reliability of longitudinal brain change: Estimations based on empirical data.

| *term* | *df* | *F-stat* | *p-value* | *partial Eta squared (η2)* |
|---|---|---|---|---|
| *Mod* | 3 | 798.347 | 0.000 | 0.318 |
| *Obs* | 3 | 625.163 | 0.000 | 0.267 |
| *Follow-Up* | 5 | 14449.280 | 0.000 | 0.934 |
| *Mod×Obs* | 9 | 5.570 | 0.000 | 0.010 |
| *Mod×Follow-Up* | 15 | 46.673 | 0.000 | 0.120 |
| *Obs×Follow-Up* | 15 | 5.890 | 0.000 | 0.017 |
| *Mod×Obs×Follow-Up* | 45 | 1.643 | 0.004 | 0.014 |

***Supplementary Table 19**. **ANOVA summary**. Modality × Follow-up Time × Number of Observations on the reliability of brain change as estimated "empirically" from the longitudinal aging dataset. See also **Figure 8** and **Supplementary Figure 12** for visualization and **Supplementary Table 20** for additional statistics. Mod = Modality. Obs. = Number of Observations.*

| Modality | Follow-up | Obs = 3 | | Obs = 5 | | Obs = 7 | | Obs = 9 | |
|---|---|---|---|---|---|---|---|---|---|
| | | mean | SD | mean | SD | mean | SD | mean | SD |
| area | 2 | 0.30 | 0.06 | 0.31 | 0.06 | 0.33 | 0.07 | 0.33 | 0.10 |
| | 4 | 0.41 | 0.06 | 0.45 | 0.05 | 0.47 | 0.07 | 0.48 | 0.09 |
| | 6 | 0.54 | 0.05 | 0.58 | 0.04 | 0.62 | 0.05 | 0.64 | 0.07 |
| | 8 | 0.66 | 0.05 | 0.71 | 0.03 | 0.75 | 0.04 | 0.77 | 0.05 |
| | 10 | 0.77 | 0.05 | 0.81 | 0.03 | 0.84 | 0.03 | 0.86 | 0.03 |
| | 12 | 0.85 | 0.04 | 0.88 | 0.02 | 0.91 | 0.02 | 0.92 | 0.03 |
| subcortical | 2 | 0.46 | 0.18 | 0.54 | 0.17 | 0.60 | 0.16 | 0.66 | 0.16 |
| | 4 | 0.58 | 0.14 | 0.65 | 0.13 | 0.71 | 0.12 | 0.75 | 0.11 |
| | 6 | 0.69 | 0.10 | 0.75 | 0.09 | 0.80 | 0.08 | 0.83 | 0.07 |
| | 8 | 0.79 | 0.07 | 0.83 | 0.06 | 0.87 | 0.05 | 0.89 | 0.04 |
| | 10 | 0.86 | 0.05 | 0.89 | 0.03 | 0.91 | 0.03 | 0.93 | 0.03 |
| | 12 | 0.91 | 0.03 | 0.93 | 0.02 | 0.95 | 0.02 | 0.95 | 0.03 |
| thickness | 2 | 0.29 | 0.06 | 0.31 | 0.05 | 0.32 | 0.07 | 0.32 | 0.09 |
| | 4 | 0.42 | 0.05 | 0.45 | 0.05 | 0.47 | 0.07 | 0.48 | 0.09 |
| | 6 | 0.56 | 0.04 | 0.60 | 0.04 | 0.62 | 0.05 | 0.64 | 0.07 |
| | 8 | 0.69 | 0.04 | 0.73 | 0.03 | 0.76 | 0.03 | 0.78 | 0.04 |
| | 10 | 0.79 | 0.03 | 0.83 | 0.02 | 0.86 | 0.02 | 0.88 | 0.03 |
| | 12 | 0.87 | 0.03 | 0.90 | 0.02 | 0.92 | 0.01 | 0.93 | 0.02 |
| volume | 2 | 0.32 | 0.05 | 0.35 | 0.05 | 0.37 | 0.07 | 0.39 | 0.10 |
| | 4 | 0.45 | 0.04 | 0.49 | 0.04 | 0.52 | 0.06 | 0.54 | 0.09 |
| | 6 | 0.58 | 0.04 | 0.63 | 0.03 | 0.67 | 0.04 | 0.69 | 0.06 |
| | 8 | 0.70 | 0.04 | 0.75 | 0.02 | 0.79 | 0.02 | 0.81 | 0.03 |
| | 10 | 0.79 | 0.04 | 0.84 | 0.02 | 0.87 | 0.02 | 0.89 | 0.02 |
| | 12 | 0.87 | 0.03 | 0.90 | 0.02 | 0.93 | 0.01 | 0.94 | 0.01 |

***Supplementary Table 20*. *Cell means***. *Average reliability of brain change as estimated "empirically" in the different Modality × Follow-up Time × Number of Observations cells. See also* ***Figure 8*** *and* ***Supplementary Figure 12*** *for visualization and* ***Supplementary Table 19*** *for additional statistics. Obs. = Number of Observations.*

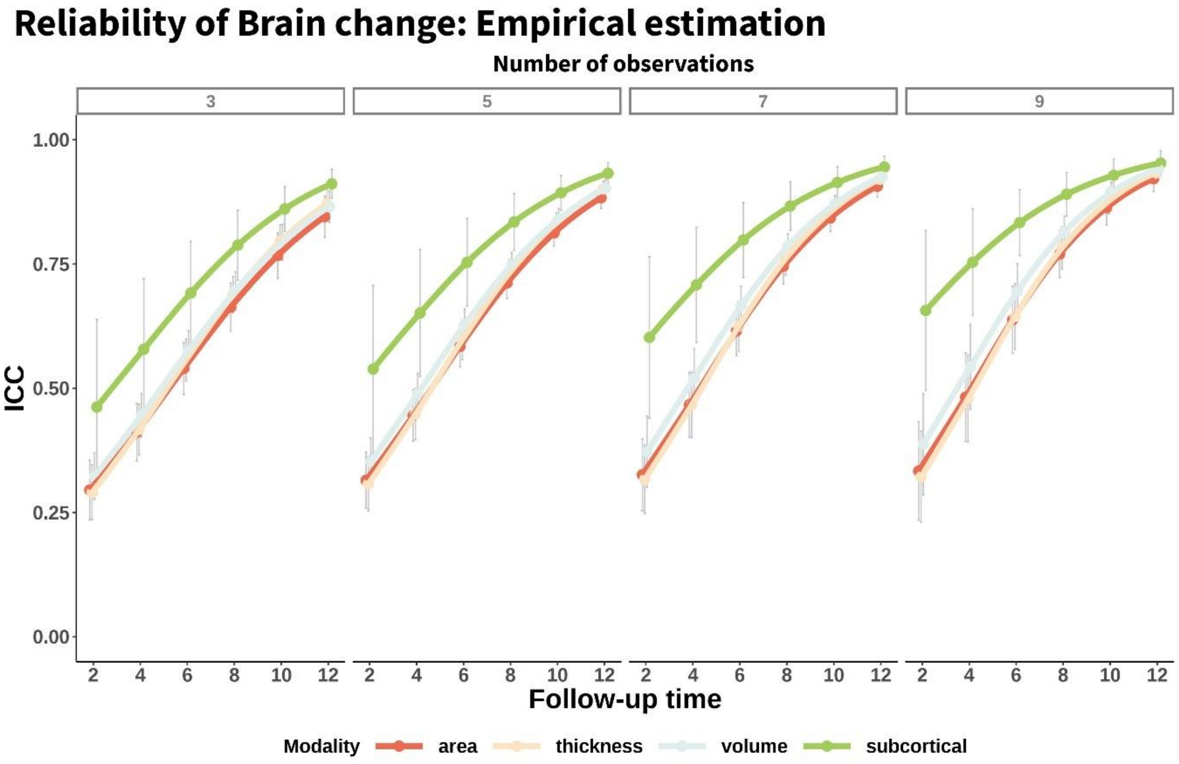


***Supplementary Figure 12. Reliability of brain change: "Empirical estimation".*** *Mean – across features - reliability (ICC) of structural brain change as a function of total follow-up time, modality, and number of equispaced observations. Reliability estimated "empirically" from the longitudinal aging dataset. Error bars represent 1 SD. See also* ***Figure 8*** *and* ***Supplementary Tables 19, 20*** *for additional statistics.*

#### Sample trends: measurement error, mean decline, age, and skewness

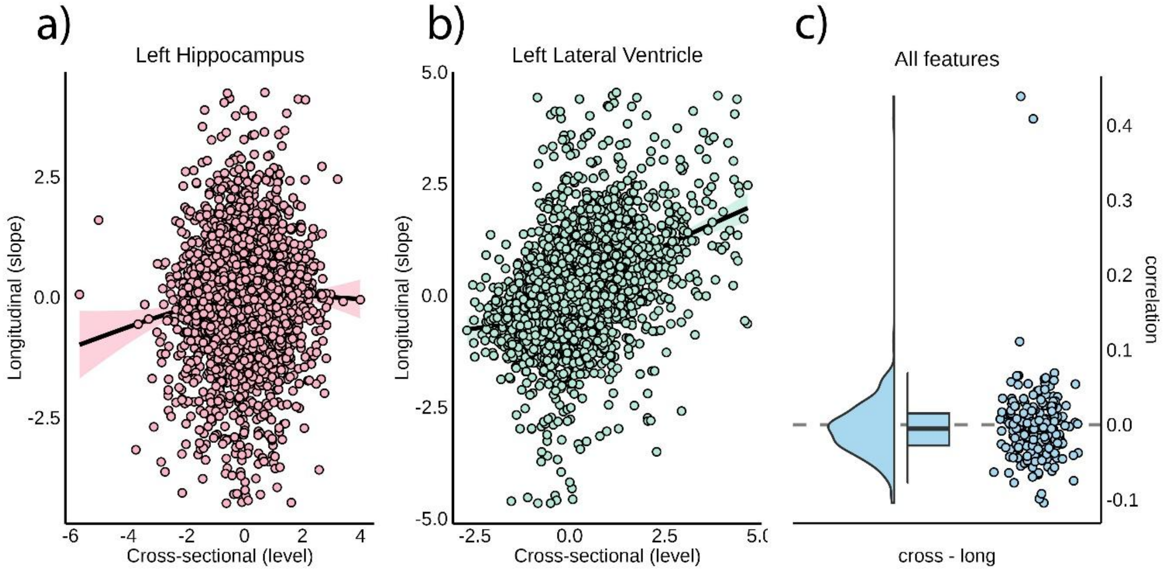


***Supplementary Figure 13. Level – Change correlation of brain change.*** *Correlation between level (cross-sectional) and change/slope (longitudinal) estimates for the a) Left Hippocampus and b) Left Lateral Ventricle. Each point represents an individual. For visualization purposes, we include GAM prediction with its confidence intervals. Cross-sectional levels are estimated as the individual random intercepts while change is computed from the model residuals as detailed in the main manuscript. Hence these measures represent level and change relative to age- and sex- matched peers. c) Correlation between level and slope across all brain features with each point representing a feature. The two outlier features are the Left and Right Lateral Ventricles (r = .44, .41, respectively). The remaining features showed correlations abs(r) ≤ .11.*

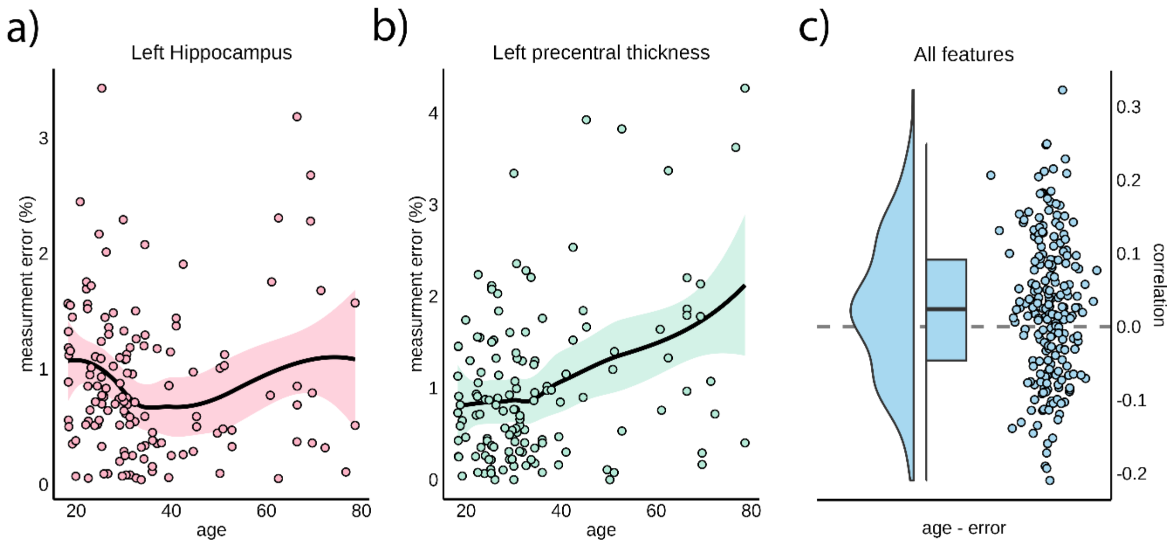


***Supplementary Figure 14. Measurement error and age.*** *Association between measurement error (percentage of test-retest change) and age for a) Left Hippocampus and b) Left precentral thickness in the S2C test-retest dataset. For visualization purposes, data was fitted with GAMs using age as a smoothing term. Ribbons represent confidence intervals. c) Pearson's correlation between measurement error and age, with each point represents a feature. Left precentral thickness is the only feature showing a significant association between measurement error and age (FDR-corrected, p < .05). The dashed line represents a null correlation.*

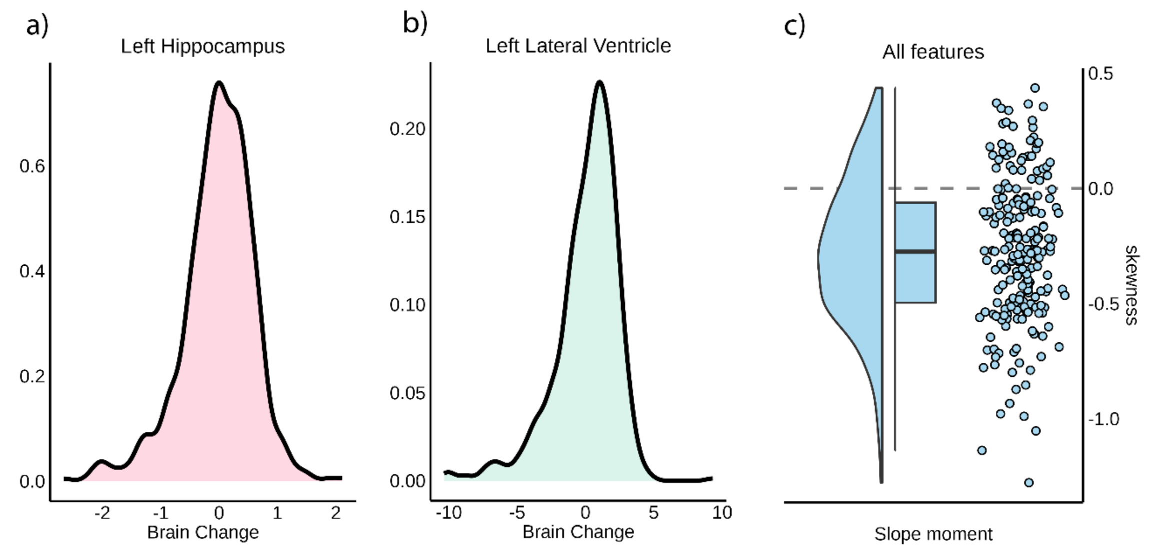


***Supplementary Figure 15. Skewness of brain change.*** *Density plots of brain change for the a) Left Hippocampus and b) Left lateral ventricle. Left lateral ventricle is one of the features with more pronounced skewness. Note that brain change is centered around 0, due to removal of aging trends in the preprocessing. c) Skewness of the distributions across all features, with each point representing a single feature. Negative skewness indicates longer right tails and vice versa. The dashed line represents no skewness.*

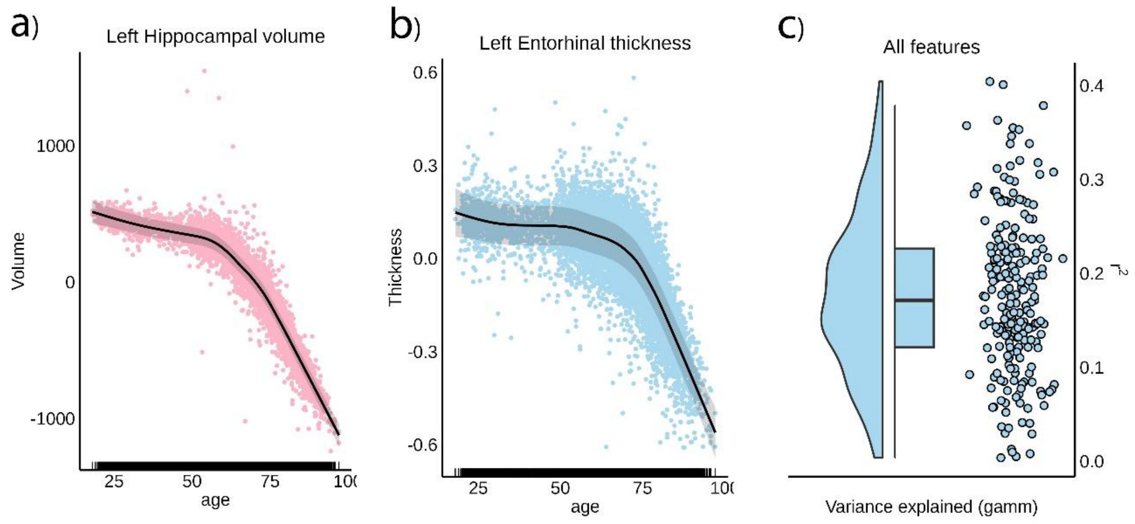


***Supplementary Figure 16. Brain trajectories in Adulthood.*** *Brain trajectories for a) Left Hippocampus and b) Left Entorhinal Thickness, modeled using generalized additive mixed models (GAMMs). Age was introduced as smoothing term, sex as factor and site, dataset, and subject as random intercepts. Points represents model residuals; ribbons indicate confidence intervals, and rug lines mark observations. c) Variance explained by the GAMMs, with each point representing a brain feature.*

#### References

Abellaneda-Pérez, K., Vaqué-Alcázar, L., Vidal-Piñeiro, D., Jannati, A., Solana, E., Bargalló, N., Santarnecchi, E., Pascual-Leone, A., Bartrés-Faz, D., 2019. Age-related differences in default-mode network connectivity in response to intermittent theta-burst stimulation and its relationships with maintained cognition and brain integrity in healthy aging. Neuroimage 188, 794–806. https://doi.org/10.1016/j.neuroimage.2018.11.036

Bates, D., Mächler, M., Bolker, B., Walker, S., 2015. Fitting Linear Mixed-Effects Models Using lme4. Journal of Statistical Software 67, 1–48. https://doi.org/10.18637/jss.v067.i01

Burgmans, S., van Boxtel, M.P.J., Gronenschild, E.H.B.M., Vuurman, E.F.P.M., Hofman, P., Uylings, H.B.M., Jolles, J., Raz, N., 2010. Multiple indicators of age-related differences in cerebral white matter and the modifying effects of hypertension. Neuroimage 49, 2083–2093. https://doi.org/10.1016/j.neuroimage.2009.10.035

Chen, B., Xu, T., Zhou, C., Wang, L., Yang, N., Wang, Z., Dong, H.-M., Yang, Z., Zang, Y.-F., Zuo, X.-N., Weng, X.-C., 2015. Individual Variability and Test-Retest Reliability Revealed by Ten Repeated Resting-State Brain Scans over One Month. PLOS ONE 10, e0144963. https://doi.org/10.1371/journal.pone.0144963

Dagley, A., LaPoint, M., Huijbers, W., Hedden, T., McLaren, D.G., Chatwal, J.P., Papp, K.V., Amariglio, R.E., Blacker, D., Rentz, D.M., Johnson, K.A., Sperling, R.A., Schultz, A.P., 2017. Harvard Aging Brain Study: Dataset and accessibility. NeuroImage, Data Sharing Part II 144, 255–258. https://doi.org/10.1016/j.neuroimage.2015.03.069

Dale, A.M., Fischl, B., Sereno, M.I., 1999. Cortical surface-based analysis. I. Segmentation and surface reconstruction. Neuroimage 9, 179–194. https://doi.org/10.1006/nimg.1998.0395

Daugherty, A.M., Raz, N., 2016. Accumulation of iron in the putamen predicts its shrinkage in healthy older adults: A multi-occasion longitudinal study. Neuroimage 128, 11–20. https://doi.org/10.1016/j.neuroimage.2015.12.045

Desikan, R.S., Ségonne, F., Fischl, B., Quinn, B.T., Dickerson, B.C., Blacker, D., Buckner, R.L., Dale, A.M., Maguire, R.P., Hyman, B.T., Albert, M.S., Killiany, R.J., 2006. An automated labeling system for subdividing the human cerebral cortex on MRI scans into gyral based regions of interest. Neuroimage 31, 968–980. https://doi.org/10.1016/j.neuroimage.2006.01.021

Ellis, K.A., Bush, A.I., Darby, D., De Fazio, D., Foster, J., Hudson, P., Lautenschlager, N.T., Lenzo, N., Martins, R.N., Maruff, P., Masters, C., Milner, A., Pike, K., Rowe, C., Savage, G., Szoeke, C., Taddei, K., Villemagne, V., Woodward, M., Ames, D., AIBL Research Group, 2009. The Australian Imaging, Biomarkers and Lifestyle (AIBL) study of aging: methodology and baseline characteristics of 1112 individuals recruited for a longitudinal study of Alzheimer's disease. Int Psychogeriatr 21, 672–687. https://doi.org/10.1017/S1041610209009405

Fischl, B., Salat, D.H., Busa, E., Albert, M., Dieterich, M., Haselgrove, C., van der Kouwe, A., Killiany, R., Kennedy, D., Klaveness, S., Montillo, A., Makris, N., Rosen, B., Dale, A.M., 2002. Whole Brain Segmentation. Neuron 33, 341–355. https://doi.org/10.1016/S0896-6273(02)00569-X

Fischl, B., Sereno, M.I., Dale, A.M., 1999. Cortical surface-based analysis. II: Inflation, flattening, and a surface-based coordinate system. Neuroimage 9, 195–207. https://doi.org/10.1006/nimg.1998.0396

Fjell, A.M., Sørensen, Ø., Wang, Y., Amlien, I.K., Baaré, W.F.C., Bartrés-Faz, D., Bertram, L., Boraxbekk, C.-J., Brandmaier, A.M., Demuth, I., Drevon, C.A., Ebmeier, K.P., Ghisletta, P., Kievit, R., Kühn, S., Skak Madsen, K., Mowinckel, A.M., Nyberg, L., Sexton, C.E., Solé-Padullés, C., Vidal-Piñeiro, D., Wagner, G., Watne, L.O., Walhovd, K.B., 2023. Sleep duration and brain atrophy – phenotypic associations and genotypic covariance. Nature Human Behaviour.

Gorgolewski, K.J., Auer, T., Calhoun, V.D., Craddock, R.C., Das, S., Duff, E.P., Flandin, G., Ghosh, S.S., Glatard, T., Halchenko, Y.O., Handwerker, D.A., Hanke, M., Keator, D., Li, X., Michael, Z., Maumet, C., Nichols, B.N., Nichols, T.E., Pellman, J., Poline, J.-B., Rokem, A., Schaefer, G., Sochat, V., Triplett, W., Turner, J.A., Varoquaux, G., Poldrack, R.A., 2016. The brain imaging data structure, a format for organizing and describing outputs of neuroimaging experiments. Sci Data 3, 160044. https://doi.org/10.1038/sdata.2016.44

Guggenheim, J.A., Williams, C., UK Biobank Eye and Vision Consortium, 2015. Role of Educational Exposure in the Association Between Myopia and Birth Order. JAMA Ophthalmol 133, 1408–1414. https://doi.org/10.1001/jamaophthalmol.2015.3556

Holmes, A.J., Hollinshead, M.O., O'Keefe, T.M., Petrov, V.I., Fariello, G.R., Wald, L.L., Fischl, B., Rosen, B.R., Mair, R.W., Roffman, J.L., Smoller, J.W., Buckner, R.L., 2015. Brain Genomics Superstruct Project initial data release with structural, functional, and behavioral measures. Sci Data 2, 150031. https://doi.org/10.1038/sdata.2015.31

Idland, A.-V., Sala-Llonch, R., Borza, T., Watne, L.O., Wyller, T.B., Brækhus, A., Zetterberg, H., Blennow, K., Walhovd, K.B., Fjell, A.M., 2017. CSF neurofilament light levels predict hippocampal atrophy in cognitively healthy older adults. Neurobiology of Aging 49, 138–144. https://doi.org/10.1016/j.neurobiolaging.2016.09.012

Idland, A.-V., Sala-Llonch, R., Watne, L.O., Brækhus, A., Hansson, O., Blennow, K., Zetterberg, H., Sørensen, Ø., Walhovd, K.B., Wyller, T.B., Fjell, A.M., 2020. Biomarker profiling beyond amyloid and tau: cerebrospinal fluid markers, hippocampal atrophy, and memory change in cognitively unimpaired older adults. Neurobiol. Aging 93, 1–15. https://doi.org/10.1016/j.neurobiolaging.2020.04.002

Kennedy, K.M., Raz, N., 2009. Aging white matter and cognition: differential effects of regional variations in diffusion properties on memory, executive functions, and speed. Neuropsychologia 47, 916–927. https://doi.org/10.1016/j.neuropsychologia.2009.01.001

LaMontagne, P., Benzinger, T. LS., Morris, J. C., Keefe, S., Hornbeck, Russ., Xiong, C., Grant, E., Hassenstab, J., Moulder, K., Vlassenko, A. G., Raichle, M. E., Cruchaga, C., Marcus, D., 2019. OASIS-3: Longitudinal Neuroimaging, Clinical, and Cognitive Dataset for Normal Aging and Alzheimer Disease. medRxiv 2019.12.13.19014902. https://doi.org/10.1101/2019.12.13.19014902

LaMontagne, P.J., Benzinger, T.L., Morris, J.C., Keefe, S., Hornbeck, R., Xiong, C., Grant, E., Hassenstab, J., Moulder, K., Vlassenko, A.G., Raichle, M.E., Cruchaga, C., Marcus, D., 2019. OASIS-3: Longitudinal Neuroimaging, Clinical, and Cognitive Dataset for Normal Aging and Alzheimer Disease. https://doi.org/10.1101/2019.12.13.19014902

Maclaren, J., Han, Z., Vos, S.B., Fischbein, N., Bammer, R., 2014. Reliability of brain volume measurements: A test-retest dataset. Sci Data 1, 140037. https://doi.org/10.1038/sdata.2014.37

Magnus, P., Birke, C., Vejrup, K., Haugan, A., Alsaker, E., Daltveit, A.K., Handal, M., Haugen, M., Høiseth, G., Knudsen, G.P., Paltiel, L., Schreuder, P., Tambs, K., Vold, L., Stoltenberg, C., 2016. Cohort Profile Update: The Norwegian Mother and Child Cohort Study (MoBa). Int J Epidemiol 45, 382–388. https://doi.org/10.1093/ije/dyw029

Marcus, D.S., Wang, T.H., Parker, J., Csernansky, J.G., Morris, J.C., Buckner, R.L., 2007. Open Access Series of Imaging Studies (OASIS): cross-sectional MRI data in young, middle aged, nondemented, and demented older adults. J Cogn Neurosci 19, 1498–1507. https://doi.org/10.1162/jocn.2007.19.9.1498

Markiewicz, C.J., Gorgolewski, K.J., Feingold, F., Blair, R., Halchenko, Y.O., Miller, E., Hardcastle, N., Wexler, J., Esteban, O., Goncavles, M., Jwa, A., Poldrack, R., 2021. The OpenNeuro resource for sharing of neuroscience data. eLife 10, e71774. https://doi.org/10.7554/eLife.71774

Miller, K.L., Alfaro-Almagro, F., Bangerter, N.K., Thomas, D.L., Yacoub, E., Xu, J., Bartsch, A.J., Jbabdi, S., Sotiropoulos, S.N., Andersson, J.L.R., Griffanti, L., Douaud, G., Okell, T.W., Weale, P., Dragonu, I., Garratt, S., Hudson, S., Collins, R., Jenkinson, M., Matthews, P.M., Smith, S.M., 2016. Multimodal population brain imaging in the UK Biobank prospective epidemiological study. Nature Neuroscience 19, 1523–1536. https://doi.org/10.1038/nn.4393

Morris, J.C., 1993. The Clinical Dementia Rating (CDR): current version and scoring rules. Neurology 43, 2412–2414. https://doi.org/10.1212/wnl.43.11.2412-a

Mueller, S.G., Weiner, M.W., Thal, L.J., Petersen, R.C., Jack, C., Jagust, W., Trojanowski, J.Q., Toga, A.W., Beckett, L., 2005. The Alzheimer's disease neuroimaging initiative. Neuroimaging Clin. N. Am. 15, 869–877, xi–xii. https://doi.org/10.1016/j.nic.2005.09.008

Nilsen, T., Brandt, I., Harris, J.R., 2019. The Norwegian Twin Registry. Twin Research and Human Genetics 22, 647–650. https://doi.org/10.1017/thg.2019.59

Nilsson, L.-G., Adolfsson, R., Bäckman, L., Frias, C.M. de, Molander, B., Nyberg, L., 2004. Betula: A Prospective Cohort Study on Memory, Health and Aging. Aging, Neuropsychology, and Cognition 11, 134–148. https://doi.org/10.1080/13825580490511026

Nilsson, L.-Gör., BÄCkman, L., Erngrund, K., Nyberg, L., Adolfsson, R., Bucht, Gös., Karlsson, S., Widing, M., Winblad, B., 1997. The betula prospective cohort study: Memory, health, and aging. Aging, Neuropsychology, and Cognition 4, 1–32. https://doi.org/10.1080/13825589708256633

Nyberg, L., Salami, A., Andersson, M., Eriksson, J., Kalpouzos, G., Kauppi, K., Lind, J., Pudas, S., Persson, J., Nilsson, L.-G., 2010. Longitudinal evidence for diminished frontal cortex function in aging. Proc. Natl. Acad. Sci. U.S.A. 107, 22682–22686. https://doi.org/10.1073/pnas.1012651108

Orban, P., Madjar, C., Savard, M., Dansereau, C., Tam, A., Das, S., Evans, A.C., Rosa-Neto, P., Breitner, J.C.S., Bellec, P., 2015. Test-retest resting-state fMRI in healthy elderly persons with a family history of Alzheimer's disease. Sci Data 2, 150043. https://doi.org/10.1038/sdata.2015.43

Petersen, R.C., Aisen, P.S., Beckett, L.A., Donohue, M.C., Gamst, A.C., Harvey, D.J., Jack, C.R., Jagust, W.J., Shaw, L.M., Toga, A.W., Trojanowski, J.Q., Weiner, M.W., 2010. Alzheimer's Disease Neuroimaging Initiative (ADNI): clinical characterization. Neurology 74, 201–209. https://doi.org/10.1212/WNL.0b013e3181cb3e25

R. A. Rigby, D. M. Stasinopoulos, 2005. Generalized additive models for location, scale and shape,(with discussion). Applied Statistics 54, 507–554.

Rajaram, S., Valls-Pedret, C., Cofán, M., Sabaté, J., Serra-Mir, M., Pérez-Heras, A.M., Arechiga, A., Casaroli-Marano, R.P., Alforja, S., Sala-Vila, A., Doménech, M., Roth, I., Freitas-Simoes, T.M., Calvo, C., López-Illamola, A., Haddad, E., Bitok, E., Kazzi, N., Huey, L., Fan, J., Ros, E., 2017. The Walnuts and Healthy Aging Study (WAHA): Protocol for a Nutritional Intervention Trial with Walnuts on Brain Aging. Front Aging Neurosci 8. https://doi.org/10.3389/fnagi.2016.00333

Raz, N., Yang, Y., Dahle, C.L., Land, S., 2012. Volume of White Matter Hyperintensities in Healthy Adults: Contribution of Age, Vascular Risk Factors, and Inflammation-Related Genetic Variants. Biochim Biophys Acta 1822, 361–369. https://doi.org/10.1016/j.bbadis.2011.08.007

Reuter, M., Rosas, H.D., Fischl, B., 2010. Highly accurate inverse consistent registration: A robust approach. NeuroImage 53, 1181–1196. https://doi.org/10.1016/j.neuroimage.2010.07.020

Reuter, M., Schmansky, N.J., Rosas, H.D., Fischl, B., 2012. Within-subject template estimation for unbiased longitudinal image analysis. Neuroimage 61, 1402–1418. https://doi.org/10.1016/j.neuroimage.2012.02.084

Routier, A., Burgos, N., Díaz, M., Bacci, M., Bottani, S., El-Rifai, O., Fontanella, S., Gori, P., Guillon, J., Guyot, A., Hassanaly, R., Jacquemont, T., Lu, P., Marcoux, A., Moreau, T., Samper-González, J., Teichmann, M., Thibeau-Sutre, E., Vaillant, G., Wen, J., Wild, A., Habert, M.-O., Durrleman, S., Colliot, O., 2021. Clinica: An Open-Source Software Platform for Reproducible Clinical Neuroscience Studies. Front Neuroinform 15, 689675. https://doi.org/10.3389/fninf.2021.689675

Sajjad, M.U., Blennow, K., Knapskog, A.B., Idland, A.-V., Chaudhry, F.A., Wyller, T.B., Zetterberg, H., Watne, L.O., 2020. Cerebrospinal Fluid Levels of Interleukin-8 in Delirium, Dementia, and Cognitively Healthy Patients. J Alzheimers Dis 73, 1363–1372. https://doi.org/10.3233/JAD-190941

Samper-González, J., Burgos, N., Bottani, S., Fontanella, S., Lu, P., Marcoux, A., Routier, A., Guillon, J., Bacci, M., Wen, J., Bertrand, A., Bertin, H., Habert, M.-O., Durrleman, S., Evgeniou, T., Colliot, O., 2018. Reproducible evaluation of classification methods in Alzheimer's disease: Framework and application to MRI and PET data. NeuroImage 183, 504–521. https://doi.org/10.1016/j.neuroimage.2018.08.042

Tremblay-Mercier, J., Madjar, C., Das, S., Pichet Binette, A., Dyke, S.O.M., Étienne, P., Lafaille-Magnan, M.-E., Remz, J., Bellec, P., Louis Collins, D., Natasha Rajah, M., Bohbot, V., Leoutsakos, J.-M., Iturria-Medina, Y., Kat, J., Hoge, R.D., Gauthier, S., Tardif, C.L., Mallar Chakravarty, M., Poline, J.-B., Rosa-Neto, P., Evans, A.C., Villeneuve, S., Poirier, J., Breitner, J.C.S., PREVENT-AD Research Group, 2021. Open science datasets from PREVENT-AD, a longitudinal cohort of pre-symptomatic Alzheimer's disease. Neuroimage Clin 31, 102733. https://doi.org/10.1016/j.nicl.2021.102733

Uribe, C., Segura, B., Baggio, H.C., Abos, A., Marti, M.J., Valldeoriola, F., Compta, Y., Bargallo, N., Junque, C., 2016. Patterns of cortical thinning in nondemented Parkinson's disease patients. Mov Disord 31, 699–708. https://doi.org/10.1002/mds.26590

Vidal-Piñeiro, D., Martin-Trias, P., Arenaza-Urquijo, E.M., Sala-Llonch, R., Clemente, I.C., Mena-Sánchez, I., Bargalló, N., Falcón, C., Pascual-Leone, Á., Bartrés-Faz, D., 2014. Task-dependent activity and connectivity predict episodic memory network-based responses to brain stimulation in healthy aging. Brain Stimul 7, 287–296. https://doi.org/10.1016/j.brs.2013.12.016

Walhovd, K.B., Fjell, A.M., Westerhausen, R., Nyberg, L., Ebmeier, K.P., Lindenberger, U., Bartrés-Faz, D., Baaré, W.F.C., Siebner, H.R., Henson, R., Drevon, C.A., Strømstad Knudsen, G.P., Ljøsne, I.B., Penninx, B.W.J.H., Ghisletta, P., Rogeberg, O., Tyler, L., Bertram, L., Lifebrain Consortium, 2018. Healthy minds 0-100 years: Optimising the use of European brain imaging cohorts ("Lifebrain"). Eur. Psychiatry 50, 47–56. https://doi.org/10.1016/j.eurpsy.2017.12.006

Walhovd, K.B., Krogsrud, S.K., Amlien, I.K., Bartsch, H., Bjørnerud, A., Due-Tønnessen, P., Grydeland, H., Hagler, D.J., Håberg, A.K., Kremen, W.S., Ferschmann, L., Nyberg, L., Panizzon, M.S., Rohani, D.A., Skranes, J., Storsve, A.B., Sølsnes, A.E., Tamnes, C.K., Thompson, W.K., Reuter, C., Dale, A.M., Fjell, A.M., 2016. Neurodevelopmental origins of lifespan changes in brain and cognition. Proc. Natl. Acad. Sci. U.S.A. 113, 9357–9362. https://doi.org/10.1073/pnas.1524259113

Walhovd, K.B., Krogsrud, S.K., Amlien, I.K., Sørensen, Ø., Wang, Y., Bråthen, A.C.S., Overbye, K., Kransberg, J., Mowinckel, A.M., Magnussen, F., Herud, M., Håberg, A.K., Fjell, A.M., Vidal-Piñeiro, D., 2024. Back to the future: omnipresence of fetal influence on the human brain through the lifespan. eLife 12. https://doi.org/10.7554/eLife.86812.2

Wood, S.N., 2017. Generalized Additive Models: An Introduction with R, 2nd ed. Chapman and Hall/CRC.

Zuo, X.-N., Anderson, J.S., Bellec, P., Birn, R.M., Biswal, B.B., Blautzik, J., Breitner, J.C.S., Buckner, R.L., Calhoun, V.D., Castellanos, F.X., Chen, A., Chen, B., Chen, J., Chen, X., Colcombe, S.J., Courtney, W., Craddock, R.C., Di Martino, A., Dong, H.-M., Fu, X., Gong, Q., Gorgolewski, K.J., Han, Y., He, Ye, He, Yong, Ho, E., Holmes, A., Hou, X.-H., Huckins, J., Jiang, T., Jiang, Y., Kelley, W., Kelly, C., King, M., LaConte, S.M., Lainhart, J.E., Lei, X., Li, H.-J., Li, Kaiming, Li, Kuncheng, Lin, Q., Liu, D., Liu, J., Liu, X., Liu, Y., Lu, G., Lu, J., Luna, B., Luo, J., Lurie, D., Mao, Y., Margulies, D.S., Mayer, A.R., Meindl, T., Meyerand, M.E., Nan, W., Nielsen, J.A., O'Connor, D., Paulsen, D., Prabhakaran, V., Qi, Z., Qiu, J., Shao, C., Shehzad, Z., Tang, W., Villringer, A., Wang, H., Wang, K., Wei, D., Wei, G.-X., Weng, X.-C., Wu, X., Xu, T., Yang, N., Yang, Z., Zang, Y.-F., Zhang, L., Zhang, Q., Zhang, Zhe, Zhang, Zhiqiang, Zhao,

K., Zhen, Z., Zhou, Y., Zhu, X.-T., Milham, M.P., 2014. An open science resource for establishing reliability and reproducibility in functional connectomics. Sci Data 1, 140049. https://doi.org/10.1038/sdata.2014.49
